# Perturb-Seqr: Signature Search Engine Integrating Connectivity Mapping Resources

**DOI:** 10.64898/2026.10.06.757137

**Authors:** Lily D. Taub, John Gardner, Ido Diamant, Daniel J.B. Clarke, Avi Ma’ayan

**Author notes:** Contributed equally (Lily D. Taub, John Gardner).

## Abstract

High-throughput transcriptomics can be used to develop Connectivity Mapping resources for identifying potential reversers and mimickers of disease signatures for drug and target discovery. Recently, several new Connectivity Mapping datasets have emerged, but no efforts were made so far to evaluate and integrate them. Motivated by this opportunity, we have developed Perturb-Seqr, a comprehensive Connectivity Mapping search engine that integrates nine small molecule transcriptomics Connectivity Maps and seven single gene perturbation followed by expression Connectivity Maps. Specifically, Perturb-Seqr contains processed data from LINCS (L1000), Tahoe-100M, Novartis Institute for Biomedical Research (DRUG-seq), the original CMap (cDNA microarrays), Ginkgo Bioworks (DRUG-seq), SciPlex (scRNA-seq), DeepCover MoA (PISA proteomics), Perturb Atlas (RNA-seq), Replogle et al. (Perturb-seq), CREEDS (cDNA microarrays), CM4AI (Perturb-seq), and RummaGEO (RNA-seq). To serve these Connectivity Maps in one unifying search engine, we converted the expression signatures from these resources into 450,271 gene sets of up- and down-regulated genes measuring the effects of 9,856 unique drug and 17,643 unique gene perturbations, targeting 917 cell lines and cell types. We demonstrate how Perturb-Seqr, and the gene sets it serves, can be used to identify targets for drugs and drugs for targets, uncover new pathways, and identify potential side effects for preclinical compounds. Overall, Perturb-Seqr is a comprehensive up-to-date Connectivity Mapping search engine and database, enabling the discovery of new drugs and targets. Perturb-Seqr is accessible from: https://perturbseqr.maayanlab.cloud/.

## Introduction

Data generated by high-throughput perturbations followed by genome-wide mRNA and protein expression can help elucidate small molecules’ mechanisms of action, identify novel bioactive compounds, enable drug repurposing, predicted side effects for preclinical compounds, as well as identify targets for drugs and drugs for targets (*1–6*). The first implementation of a massive collection of single perturbation followed by genome-wide expression to develop a community resource was published in 2000 by Hughes et al. (*7*). The authors generated 300 transcriptomic profiles using cDNA microarrays by measuring the response of *S. cerevisiae* to 267 gene deletions and mutations, as well as 13 small molecule perturbagens. Using these profiles, unknown gene functions and novel drug targets were discovered. Several years later, the landmark “Connectivity Map” publication described a collection of high-throughput drug perturbations followed by genome-wide gene expression applied to four human cancer cell lines (*8*). This original Connectivity Map, also known as CMAP, contained 1,309 small molecules and drugs, including most of the FDA-approved drugs, applied in different concentrations, measuring gene expression after 6 hours of the initial perturbation. The CMAP resource has been widely used for drug discovery and drug repurposing, and this success prompted the NIH to establish the Library of Integrated Network-Based Cellular Signatures (LINCS) Common Fund program(*9*).

A cornerstone dataset that was produced by the LINCS program was the L1000 chemical and gene perturbation Connectivity Maps (*10*). The chemical perturbations dataset has been widely utilized by the community to construct models and predict compounds to reverse disease conditions. For example, the L1000 data has been used to predict chemical reversers of an Amyotrophic lateral sclerosis (ALS) disease signature derived from whole blood collected from a large disease cohort (*11*), and to identify candidate compounds to target a pan-cancer expression pattern of stemness (*12*). Additionally, several predictive models have been developed to infer the effects of perturbations in untested conditions. The LINCS L1000 dataset was used in these studies for training and benchmarking (*11–13*). In addition, in order to make gene signatures derived from perturbation followed by expression experiments available to the community, several L1000 signature search engines have been developed (*14–19*). Notably, the LINCS L1000 gene knockout perturbation followed by expression dataset was never published, and as a result was not widely utilized by the community.

To create other Connectivity Mapping resources, not all the data needed to come from the same source. For example, a crowd-sourcing project called CRowd Extracted Expression of Differential Signatures (CREEDS) (*20*) created a Connectivity Mapping resource by mining published cDNA microarray studies from the Gene Expression Omnibus (*21*). A later effort called RummaGEO (*22*) automatically identified and computed signatures from gene and drug perturbations followed by expression from human and mouse bulk RNA-seq studies published in the Gene Expression Omnibus (GEO) (*23*). These and other similar efforts complement existing Connectivity Mapping efforts and have the advantage of heterogeneity of signatures and the opportunity for continual growth. More recently, the Perturb-seq assay (*24*) and the DRUG-seq platform (*25*) have been developed. As a result, this significantly expanded the availability of Connectivity Mapping resources. Perturb-seq combines high-throughput single-cell RNA-sequencing with CRISPR-based gene editing to enable large-scale gene perturbation followed by expression screens. DRUG-seq optimizes multi-well processing to enable large compound perturbation followed by expression screens across a variety of dosages and time points. Together, these technologies have been used to create several new Connectivity Mapping datasets that capture transcriptional responses to hundreds of thousands of perturbations (*10*, *26–29*).

Here we present Perturb-Seqr, a unified signature search engine that integrates 16 Connectivity Maps generated from 12 resources (Table 1). By harmonizing expression signature data from across 12 resources, Perturb-Seqr enables users to identify consistent matching perturbations across assay technologies, experimental protocols, and biological contexts. In addition, by comparing the signatures created from the different sources, previously unknown drug-drug, target-target, and drug-target similarity can be discovered. Furthermore, we demonstrate the utility of Perturb-Seqr in two use cases. For the first use case, we identify potential genetic and chemical mimickers of exercise, by analyzing signatures generated by the MoTrPAC adult acute exercise study (*30*, *31*). For the second use case, signatures computed from the lung squamous cell carcinoma (LSCC) cohort from the CPTAC3 study (*32*) are analyzed by Perturb-Seqr to suggest drugs and targets. In addition, we demonstrate that the data collected for Perturb-Seqr can be used as training data for developing adverse drug reaction prediction models.

**Table 1.**
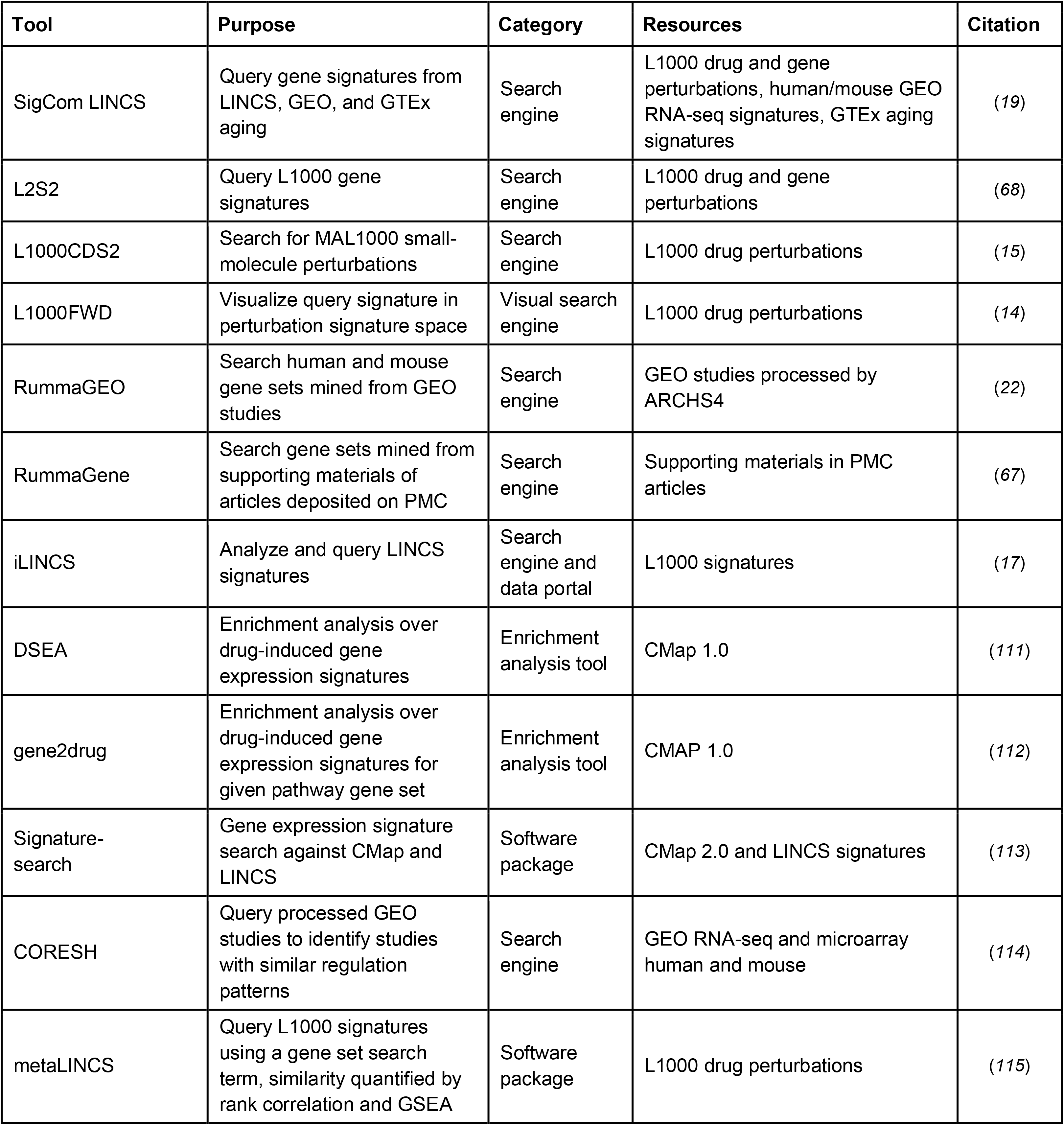
Other gene expression signature search engines and relevant gene set enrichment analysis tools serving gene sets derived from Connectivity Map resources.

| Tool | Purpose | Category | Resources | Citation |
| --- | --- | --- | --- | --- |
| SigCom LINCS | Query gene signatures from LINCS, GEO, and GTEx aging | Search engine | L1000 drug and gene perturbations, human/mouse GEO RNA-seq signatures, GTEx aging signatures | (19) |
| L2S2 | Query L1000 gene signatures | Search engine | L1000 drug and gene perturbations | (68) |
| L1000CDS2 | Search for MAL1000 small-molecule perturbations | Search engine | L1000 drug perturbations | (15) |
| L1000FWD | Visualize query signature in perturbation signature space | Visual search engine | L1000 drug perturbations | (14) |
| RummaGEO | Search human and mouse gene sets mined from GEO studies | Search engine | GEO studies processed by ARCHS4 | (22) |
| RummaGene | Search gene sets mined from supporting materials of articles deposited on PMC | Search engine | Supporting materials in PMC articles | (67) |
| iLINCS | Analyze and query LINCS signatures | Search engine and data portal | L1000 signatures | (17) |
| DSEA | Enrichment analysis over drug-induced gene expression signatures | Enrichment analysis tool | CMap 1.0 | (111) |
| gene2drug | Enrichment analysis over drug-induced gene expression signatures for given pathway gene set | Enrichment analysis tool | CMAP 1.0 | (112) |
| Signature-search | Gene expression signature search against CMap and LINCS | Software package | CMap 2.0 and LINCS signatures | (113) |
| CORESH | Query processed GEO studies to identify studies with similar regulation patterns | Search engine | GEO RNA-seq and microarray human and mouse | (114) |
| metaLINCS | Query L1000 signatures using a gene set search term, similarity quantified by rank correlation and GSEA | Software package | L1000 drug perturbations | (115) |

## Methods

### Data Processing and Signature Generation

#### LINCS L1000

Gene expression signatures from the LINCS L1000 chemical perturbations (*10*) were derived by utilizing the characteristic direction (CD) (*33*) coefficients computed on batch-corrected L1000 data measuring the expression of the 978 landmark genes and the 11,350 inferred genes. Signatures were filtered to optimize replicate consistency using an empirical permutation test (*14*). The gene expression signatures for LINCS L1000 single-gene CRISPR knockouts (*10*) were computed following the same procedure. To evaluate whether filtering for replicate consistency had improved performance, the Dex-Benchmark (*34*) and the gene known-relationship recall benchmark were used.

#### Tahoe-100M

The dataset was retrieved from the HuggingFace repository on October 6th, 2025. Pseudobulked differential expression (*35*) was computed with DESeq2 (*36*). Gene sets were created by filtering the pseudobulked differential expression for adjusted p-values of <= 0.05 and the top 100 up- and 100 down-regulated protein-coding genes were extracted to create gene sets.

#### Novartis DRUG-seq MoABox

Limma-trend (*37*) was used to compute differential expression from the Novartis Institutes for BioMedical Research (NIBR) DRUG-seq MoABox dataset (*38*). This dataset was obtained from the Novartis/DRUG-seq U2OS MoABox repository (*26*). Gene sets were constructed by filtering for genes with adjusted p-values < 0.01.

#### Original Microarray CMAP

Gene expression signatures from the. original Connectivity Mapping dataset (*8*) were processed from the provided Z-scores using a z-score threshold of 0.95. Further filtering was applied to include only protein-coding genes, and capping gene set size at 100 genes per set with a minimum of 5.

#### Ginkgo Bioworks GDPx1 and GDPx2

Differential expression analysis for the Ginkgo Bioworks GDPx1 (*39*) and GDPx2 (*27*) was performed with limma-voom (*40*) as implemented in the Limma R package (v3.62.2). Raw counts were normalized using the Trimmed Mean of M-values in edgeR (v4.4.2), and gene sets were created by filtering for genes with a differential expression p-value of < 0.05.

#### SciPlex

Raw UMI count matrices for SciPlex3 (*41*) scRNA-seq chemical perturbation screens in A549, MCF7, and K562 cell lines were obtained from GEO (GSE139944). To enable the use of CD analysis on single-cell data, pseudobulk expression profiles were generated by randomly binning cells into replicates. Subsequently, top 500 up- and down-regulated genes were taken to define the gene sets, with a minimum of 5 genes per set, filtered to only include protein-coding genes.

#### DeepCover MoA

Pre-computed differential protein abundance scores for 875 drug perturbations in HCT116 cells were obtained from the DeepCover MoA (*42*) proteomics resource. Gene sets were constructed by selecting the top 100 proteins with positive scores as up-regulated and the top 100 with negative scores as down-regulated.

#### CREEDS

Gene set signatures for CREEDS Chem and CREEDS Gene were both obtained from CREEDS (*20*), where differential expression was computed from curated GEO microarray studies using the CD method. The Gene Matrix Transposed (GMT) files for these datasets were supplemented with additional GEO (*21*) signatures from Harmonizome (*43*). These additional signatures were processed by the same CD pipeline and merged after computing the set difference of perturbation names between Harmonizome GEO and CREEDS libraries. Only microarray-platform signatures from human, mouse, and rat were unified into the final gene sets.

#### RummaGEO

Drug and gene perturbation gene sets were obtained from RummaGEO (*22*) directly without any processing. RummaGEO mines signatures automatically and computes differential expression signatures from GEO RNA-seq studies for both humans and mice using limma-voom (*40*) applied to the uniformly aligned samples from ARCHS4 (*44*).

#### PerturbAtlas

The PerturbAtlas (*45*) resource provides pre-computed DESeq2 (*36*) differential expression signatures for bulk RNA-seq gene perturbation followed by expression data from human and mouse. The human data was filtered by applying a threshold p-value of < 0.05, and transcripts were mapped to protein-coding gene symbols. The mouse data followed the same process, but genes were additionally mapped to their human orthologs. Gene sets were created from the top 100 up- and 100 down-regulated genes ranked by the signed significance score (log2FC) * -log10(padj), with a minimum of 5 genes per set.

#### Replogle et al

(*29*): Z-normalized pseudobulk expression matrices were downloaded from Figshare (*46*). Gene sets were created by filtering to include only protein-coding genes, and by applying a z-score threshold of |z| >= 0.5 for the two K562 screens, and |z| >= 1.0 for the RPE1 screen. Positive z-scores represent up-regulation and negative z-scores down-regulation. Perturbation signatures with fewer than 5 genes per set were discarded. The K562 genome-wide dataset was combined with the K562 essential, and RPE1 essential Perturb-seq datasets into a single gene set library.

#### CM4AI

To generate gene sets for CM4AI (*47*), pseudobulked expression profiles were created from the single-cell data following the process described by the authors of the CM4AI study. Pseudo-replicates were compared to plate-matched control profiles generated by sampling single cells treated with non-targeting control sgRNAs. Limma-voom (*40*) was used to generate differential expression signatures from the pseudobulked profiles. The up- and down-regulated gene sets were created by setting a threshold of an adjusted p-value < 0.05 and gene set size was capped at 2000 genes per perturbation.

### Benchmarking the Signature and Gene Set Generation Pipelines

Signature computation workflows for creating the gene sets from the drug and small chemical perturbation resources were benchmarked to optimize each workflow. The benchmarking included comparing the choice of differential expression analysis method, filtering decisions, and gene set extraction size thresholds. Most of the benchmarks applied to the chemical perturbations followed by expression resources utilized the Dex-Benchmark (*34*) resource. To compare data processing pipelines, signature computation, and gene set extraction strategies for the gene perturbation followed by expression Connectivity Mapping resources, the known relationship recall benchmark from Celik et al. (*48*) was adapted. Pairs of gene sets whose perturbation genes have a known relationship in CORUM (*49*), HuMAP (*50*), Reactome (*51*), SIGNOR (*52*), and StringDB (*53*) were annotated as “expected” relationships. Cosine similarity was used to compute similarity between all pairs of signatures, and the fraction of “expected” relationship pairs that was recovered in the top or bottom 5% of the cosine similarity distribution formed by all cross-target signature pairs was recorded.

### Gene ID Mapping

Gene sets generated from all the resources were post-processed to align gene identifiers. First, genes annotated as non-protein-coding in the National Center for Biotechnology Information’s (NCBI) Gene database (*54*) were removed, and only human and mouse signatures were retained from sources that had other species. Mouse genes were translated to their human orthologs using the Mouse Genome Informatics (MGI) homology table (*55*).

### Drug ID Mapping

To ensure consistency across the drugs and small molecules that are used across the Connectivity Mapping resources, all terms used to describe drugs and other compounds were normalized by their PubChem CID. The PubChem CID normalization was used to provide consistent mechanism of action (MoA) labels for drug perturbation gene set signatures from the Drug Repurposing Hub (*56*).

### Calculating Signature Similarity

The signed Jaccard index was used to evaluate the similarity between two signatures as defined below:

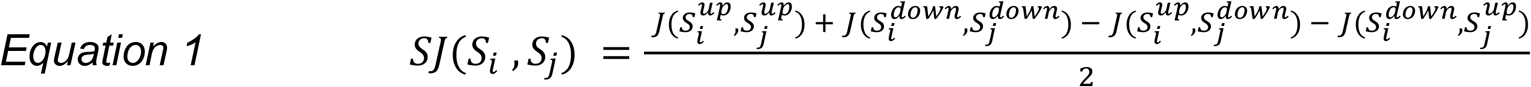

where *S^up^* and S^down^ denote the up- and down-regulated gene sets. The signed Jaccard index is bound between [-1,1] where 1 represents signatures with the same transcriptional effect, and -1 represents signatures with opposite transcriptional effects. A signed Jaccard index of 0 indicates unrelated signatures. Within each Connectivity Map gene set library, the signed Jaccard index was calculated for each pairwise combination of signatures. Individual perturbations, defined by the perturbagen ID, cell line, dosage (if applicable), and exposure time, that were not associated with a pair of up and down gene sets in the Connectivity Map library were excluded.

### The Dexamethasone, Bortezomib, and HDAC Inhibitors Benchmarks

Dexamethasone, bortezomib, and HDAC inhibitor perturbation signatures were extracted from each Connectivity Map gene set library. Small molecule perturbagens were classified as HDAC inhibitors based on OpenTargets drug MoA data (version 26.03) (*57*). Signatures were ordered based on differential expression significance, determined by the adjusted p-value computed with limma-voom (*40*) and DESeq2 (*36*), or the Characteristic Direction (*33*) coefficient. Ranked signatures for dexamethasone perturbations were submitted for gene set enrichment analysis against the ChEA (*58*) 2022 library in Enrichr (*59*) and the ranks of gene sets associated with NR3C1 were examined as described in the Dex-Benchmark (*34*). Ranked signatures for bortezomib perturbations were compared to the Reactome (*60*) 2024 apoptosis pathway from Enrichr (*59*). HDAC inhibitor ranked signatures were compared to consensus gene sets created from each Connectivity Map library. These consensus gene sets were constructed from all the HDAC inhibitors up and down gene sets. If the consensus gene set constructed from the union of all HDAC inhibitors was more than 1,000 genes, the consensus was trimmed by iteratively eliminating genes found in fewest sets. For each benchmark, a running sum bridge plot was generated by walking through the ranked genes in a signature and incrementing the sum if a ranked gene is in the reference gene set and decrementing the sum otherwise. For each drug benchmark, running sums were averaged over all reference gene sets.

### Benchmarking Gene Perturbation Followed by Expression Signatures

Gene set enrichment analysis was also applied to benchmark the gene perturbation followed by expression Connectivity Mapping gene set libraries. Each gene perturbation followed by expression signature was compared to a reference gene set composed of the 1,000 genes that had the most correlated expression with the perturbed gene based on data from ARCHS4 (*44*). The ARCHS4 human gene-gene co-expression correlation matrix (v2.4) was used to obtain these gene sets. The matrix consists of pairwise Pearson correlation coefficients computed for all human genes based on 10,000 randomly selected ARCHS4 samples. All benchmarking analyses were implemented as described in the Dex-Benchmark protocol (*34*).

### Identifying Drug-Gene Perturbational Mimickers

Perturbational mimickers are defined as a pair of signatures where the overlaps between the up- and the down-regulated gene sets are both significant at an FDR corrected p-value of < 1×10^-5^. Statistical significance of the overlap is calculated with Fisher’s exact test. Small molecule and gene perturbagen mimicking pairs were ranked based on the product of Fisher’s exact p-value for the overlaps among their up- and down-regulated genes. Mimicking pairs were classified as “unknown” if the perturbed gene was not a known target of the drug as defined in the OpenTargets database version 26.03 (*57*).

### Pathway Discovery by Constructing Gene-Gene Mimicker Subnetworks

Mimicking gene perturbations were identified using the same method as described above for identifying drug-gene perturbational mimickers. Communities of mimicking gene perturbagens were identified using the Louvain community detection algorithm (*61*) as implemented in the NetworkX (*62*) Python package. Protein-protein interaction evidence for pairs of identified mimicking genes was downloaded from the STRING database v12.0 (*53*).

### Gene Set Library Visualization with UMAP

Uniform Manifold Approximation and Projection (UMAP)(*63*) plots of gene set libraries were generated by first mapping the gene sets into Term Frequency-Inverse Document Frequency (TF-IDF) vectors (*64*). For libraries where the number of unique genes across all sets exceeded the number of sets, the resulting sparse matrix was further reduced with Singular Value Decomposition (SVD). Matrices were then embedded using UMAP (min_dist=0.1, spread=1.0) and clusters were assigned by applying the Leiden algorithm (*61*) (resolution=1.0) on a k-nearest-neighbors graph (n_neighbors=30). For gene set libraries constructed from single gene perturbation datasets, each cluster was annotated with the GO Biological Process (*65*) term for which it was most specifically enriched. Enrichment was assessed by Fisher’s exact test on a contingency table comparing, for a given GO Biological Process term *A*, the number of sets for which *A* was the most significantly enriched term inside versus outside the cluster. The Benjamini-Hochberg p-value correction was applied. Broad terms, as defined by the GOATOOLS (*66*) package (v 1.5.2) were filtered out. Outlier signatures beyond three standard deviations from the mean of the UMAP coordinates were removed.

### Developing the Perturb-Seqr Signature Search Engine

The Perturb-Seqr enrichment analysis search engine inherits its architecture from the previously published search engines Rummagene (*67*), RummaGEO (*22*), and L2S2 (*68*). Perturb-Seqr stores user queries, gene backgrounds, and all the gene sets from the ingested libraries as bit-vectors in random access memory (RAM) for efficient processing of the Fisher’s exact test via a standalone Rust API. The Perturb-Seqr search engine extends the previously published enrichment analysis search engines to enable users to select from the 16 distinct gene set libraries. The gene set libraries were created from different resources that use different assays. Therefore, each library has a different gene set background determined by the union of all genes present in each library. Hence, gene sets are represented as separate bit-vectors per library in the memory of the Rust API. Fisher’s exact test is then computed on a contingency table whose entries are filled from a user query of the gene set, a reference gene set arising from one of 16 distinct libraries, the overlap between these two sets, and the library-specific background. The query gene set is filtered to only include genes that are present in the library-specific background so only genes found in the library enter the test. Benjamini–Hochberg (*69*) (BH) correction is applied to the raw Fisher p-values independently for each library, using the total number of gene sets in each library as the number of independent tests. For up- and down-gene-set queries, Perturb-Seqr returns mimickers and reversers from each Connectivity Mapping resource following a similar procedure. Mimickers are defined as the sum of the intersection between the user-submitted and Perturb-Seqr up gene sets, and the intersection between the user-submitted and Perturb-Seqr down gene sets. A reverser is analogously defined in the opposite direction. These overlaps fill the entries in the contingency table for computing the Fisher’s exact test, and Benjamini-Hochberg adjusted p-values are computed to correct for multiple hypotheses testing. When a single gene set is submitted as a query, the results report the drug and gene perturbations that collectively up- or down-regulate the expression of the overlapping genes using the standard Fisher’s exact test with BH correction.

### Developing the Perturb-Seqr Search Engine Website

Gene sets from all Connectivity Mapping libraries are ingested into a Postgres Database accessible to the Perturb-Seqr frontend and the companion enrichment engine API. The Perturb-Seqr web server application is containerized into different independent components with Docker Compose and deployed on a Kubernetes cluster. The frontend is implemented using TypeScript and Tailwind CSS in the Next.js web framework. Interactive butterfly bar charts to visualize top enrichment results are implemented using components from the visx JavaScript library with colors and bar scaling defined using D3.js (*70*). The frontend of the application is connected to a PostgreSQL database through a GraphQL endpoint enabled by PostGraphile so that the database schema is exposed to users as a type-safe GraphQL API, which additionally provides programmatic access to Perturb-Seqr. A Python package available from the project’s GitHub repository wraps this API, returning results from single and paired (up/down) gene set queries and gene set term searches are ingested as pandas DataFrames. The frontend and the PostgreSQL database are connected to a standalone Rust microservice through a PL/Python wrapper function embedded in PostgreSQL to process gene set and metadata queries. Gene set libraries and pre-computed UMAP visualizations are served from an S3-compatible object storage via MinIO and all assets are available for download via links displayed on the frontend.

## Results

### A Large Scale, Context Diverse Gene Set Compendium

Perturb-Seqr provides access to 450,271 gene sets extracted from nine small molecule perturbations followed by expression resources, and seven gene perturbations followed by expression resources (Fig. 1A). Drug perturbation followed by expression gene sets were extracted from differential expression signatures created from LINCS L1000 (*10*), Tahoe-100M (*35*), Novartis Institutes for Biomedical Research (NIBR) DRUG-seq (*26*), CMap (*8*), RummaGEO (*22*), SciPlex (*41*), Ginkgo Bioworks (*27*, *39*), CREEDS (*20*) and DeepCover MoA (*42*) (Table 2). Gene perturbation followed by expression gene sets were extracted from differential expression signatures created from LINCS L1000 (*10*), Perturb Atlas (*45*), RummaGEO (*22*), CREEDS (*20*), CM4AI KOLF2.1J genome-wide Perturb-Seq (*47*), and Replogle at al. Perturb-Seq (*29*) (Table 3). Gene identifiers across libraries were normalized to NCBI gene symbols and pruned to only include protein-coding genes (Fig. S1). In total, the collection captures transcriptomics responses to 17,643 unique gene perturbations and 9,856 unique small molecule perturbations across 917 cell lines, cell types, and tissues (Fig. 1B). The chemical perturbations were applied in a variety of concentrations and time points across the resources (Tables 2 and 3). The most applied chemical and gene perturbagens, and the most used cell lines, are related to cancer (Fig. 1C-D). The genes that are most differentially expressed after single gene perturbations are involved in a range of cellular functions (Fig. S2). UMAP (*63*) visualization of each processed Connectivity Mapping library shows distinct MoA clustering for the drug perturbations followed by expression resources, and distinct GO term enrichment clusters for the gene perturbation followed by expression resources (Fig. S3). These clusters highlight functional relationships among the perturbagens captured by each resource. The MoA clustering of groups of drugs and other small molecules with known MoAs near compounds with unknown MoAs suggest that the data can be used to assign MoAs to small molecules that do not have a known MoA based on their transcriptional responses. The same concept can be applied for assigning functions for genes based on clustering by function for the gene perturbation signatures, as well as for assigning new targets for drugs and drugs for targets.

**Fig. 1.**
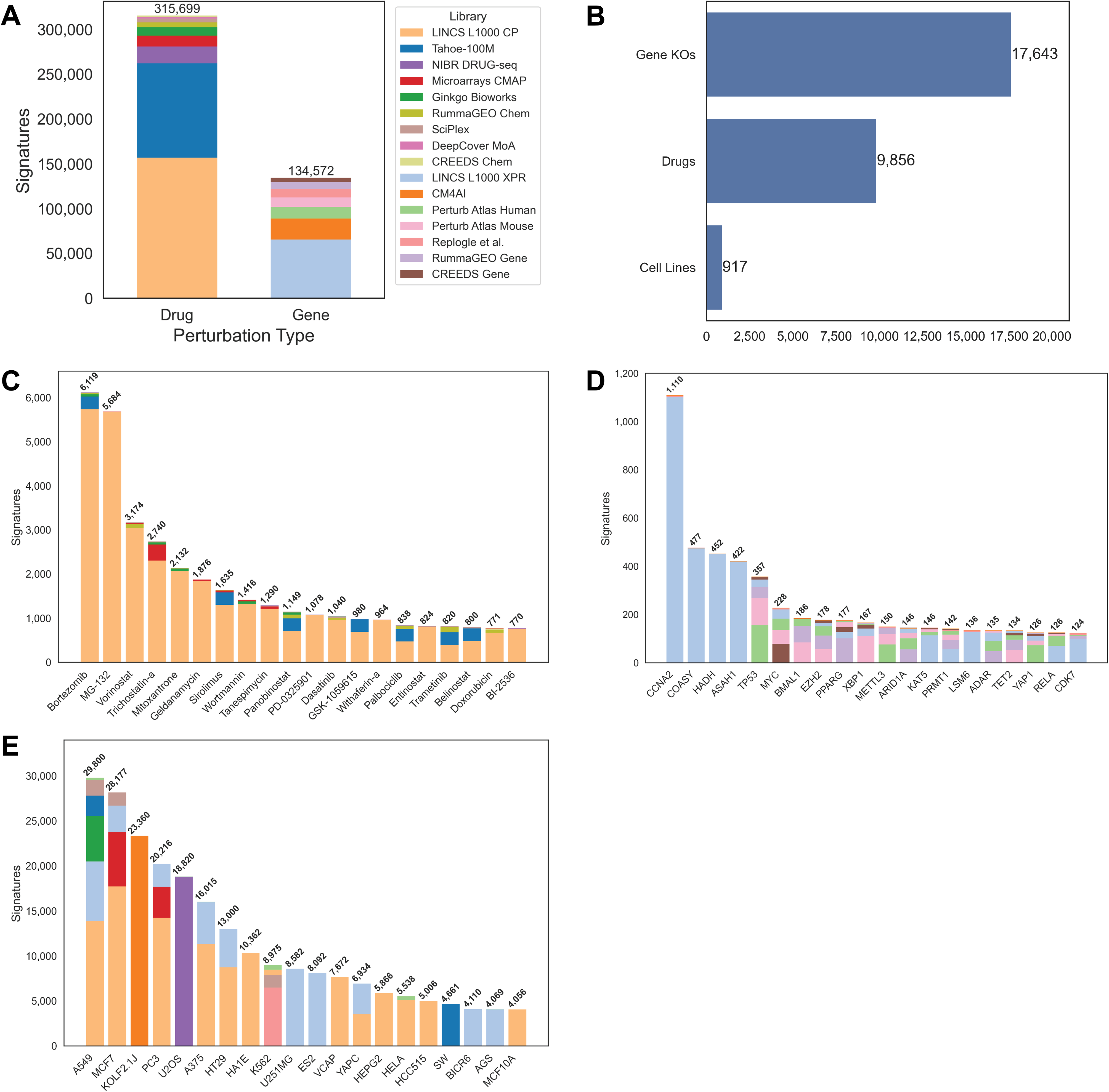
Peturb-Seqr descriptive statistics. (A) Distribution of drug and gene perturbation gene sets across the Connectivity Mapping resources. (B) Unique gene perturbations, drug perturbations, and cell lines. (C) Most common single gene perturbations. (D) Most common single drug perturbations. (E) Most common cell.

**Table 2.**
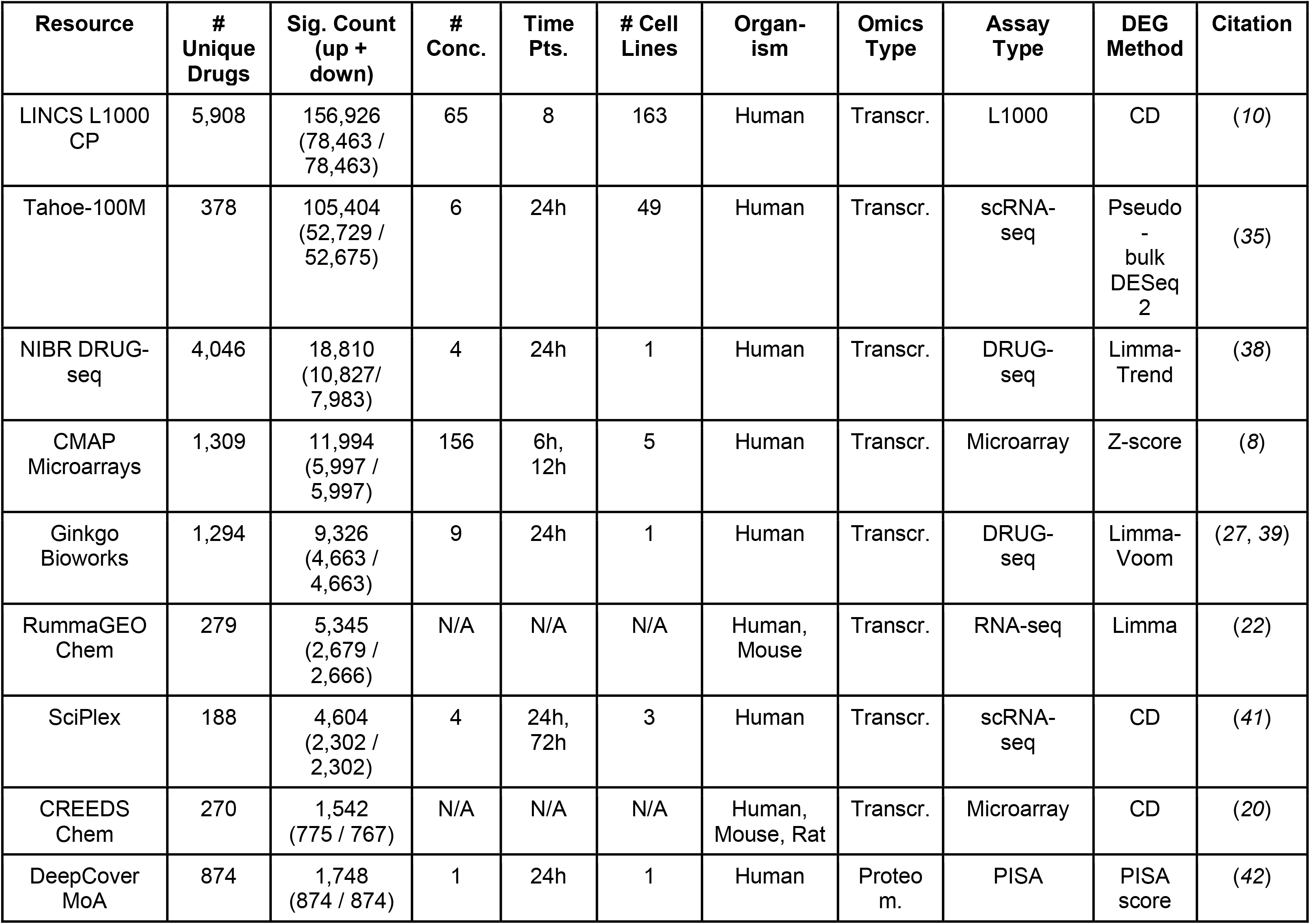
Drug perturbation followed by expression Connectivity Mapping resources processed for Perturb-Seqr. Due to unstandardized metadata formats, cell lines, and perturbation metadata for RummaGEO Chem and CREEDS Chem was not quantified.

**Table 3.** Gene perturbation followed by expression Connectivity Mapping resources processed for Perturb-Seqr. Due to unstandardized metadata formats in community derived resources, cell lines, and perturbation metadata for Perturb Atlas, RummaGEO Gene, and CREEDS Gene was not quantified.

| Resource | # Unique Genes | Sig Count (up + dn) | Pert Type | Time Pts. | # Cell Lines | Organism | Omics Type | Assay Type | DEG Method | Cites |
| --- | --- | --- | --- | --- | --- | --- | --- | --- | --- | --- |
| LINCS L1000 XPR | 6,962 | 65,672 (32,836 / 32,836) | CRISPR KO | 96h | 21 | Human | Transcr . | L1000 | CD | (10) |
| Perturb Atlas Human | 2,657 | 12,852 (6,438 / 6,414) | KD, KO, OE, Mut | N/A | N/A | Human | Transcr . | Bulk RNA-seq | DESeq2 | (45) |
| Perturb Atlas Mouse | 2,485 | 10,743 (5,367 / 5,376) | KD, KO, OE, Mut | N/A | N/A | Mouse | Transcr . | Bulk RNA-seq | DESeq2 | (45) |
| Replogle et al. | 3,098 | 9,207 (5,583 / 3,624) | CRISPRi | 6-8 days | 2 | Human | Transcr . | Perturb-seq | Z-score | (29) |
| RummaGEO Gene | 1,002 | 7,960 (4,017 / 3,943) | Various | N/A | N/A | Human, Mouse | Transcr . | Bulk RNA-seq | Limma | (22) |
| CREEDS Gene | 1,122 | 4,778 (2,388 / 2,390) | KO, KD, OE | N/A | N/A | Human, Mouse, Rat | Transcr . | Microarray | CD | (20) |
| CM4AI | 11,680 | 23,360 (11,680 / 11,680) | CRISPRi | 6 days | 1 | Human | Transcr . | scRNA-seq | Pseudo-bulk Limma-Voom | (47) |

### Benchmarking Perturbational Consistency

Within each Connectivity Map resource gene set similarity among matched perturbations was quantified with the signed Jaccard index (*20*). The distribution of signed Jaccard indexes for matched perturbagens was compared to the distribution for randomly paired perturbagens in all resources except for the CM4AI KOLF2.1J genome-wide perturbation atlas (*47*), which has a single replicate per perturbation (Fig. 2). Across all small molecule and gene perturbation libraries, the average similarity of matched perturbagen gene sets was significantly higher than the average similarity of random perturbagen pairs. Despite this, the median similarity between gene sets for matched perturbagens was below 0.5 on a -1 to 1 scale, where -1 indicates opposite effect and 1 indicates the same effect. This could be explained by the diverse cellular contexts captured in each library. The libraries with the fewest distinct cell lines had the highest average consistency among perturbation replicates, namely, Replogle et al. (*29*) (K562 and RPE1 cell lines) and NIBR DRUG-seq (*38*) (U2OS cell line). The UMAP visualization of the libraries also explains this by showing that signatures commonly cluster by cell line (Fig. S2). Despite this, there were some small molecule and gene perturbagens that had highly consistent transcriptional effects across cell lines and conditions (Table S1, S2), suggesting that these perturbagens induce context agnostic cellular effects. The small molecule perturbagens that showed the most consistent genome-wide gene expression changes were enriched for HDAC inhibitors, while the compounds that displayed the most idiosyncratic effects were enriched for DNA methyltransferase inhibitors (Fig. S4). It is known that genes silenced by DNA methylation are tissue specific and inherited within cell lineages (*71*), leading to context-dependent transcriptional responses. In contrast, HDACs target proteins in well-preserved cellular processes such as proliferation, differentiation, and cell death (*72*). HDAC inhibitors have been also shown to regulate a core set of genes across distinct tumor types (*73*).

**Fig. 2.**
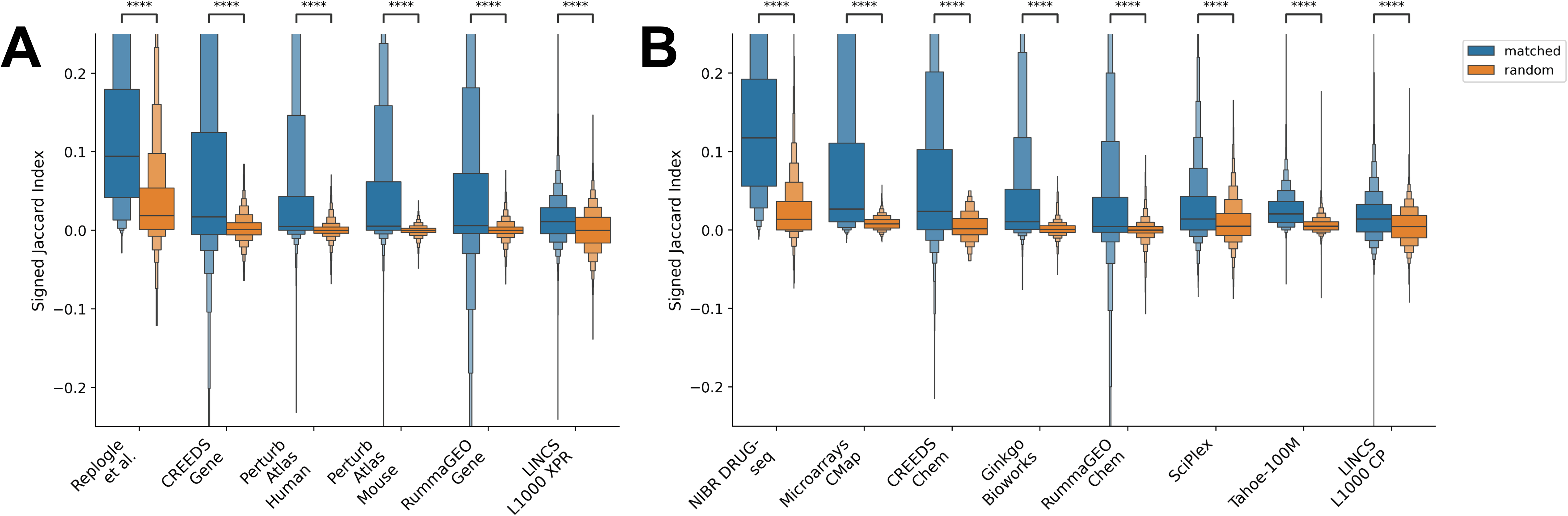
Transcriptional response consistency among the same perturbations within each Connectivity Map resource. Consistency is measured as the signed Jaccard index between two signatures. (A) Consistency among the same gene perturbations. Consistency among the same drug perturbations. Matching signatures for the same gene and drug perturbations are compared to consistency between random pairs within each resource. ****: BH-corrected p-value <= 1.00e-04 for Welch’s t-test.

### Benchmarking the Quality of the Connectivity Mapping Resources with Silver Standards

To evaluate how well signatures from each small molecule perturbation Connectivity Map produced expected known biological relationships, we first employed the Dex-Benchmark (*34*). The Dex-Benchmark is a “silver standard” benchmark that asks how well differentially expressed genes induced by the drug dexamethasone are also established targets of the glucocorticoid receptor (NR3C1) based on ChIP-seq profiling of this nuclear receptor which is the known target of dexamethasone. Since some Connectivity Mapping resources did not contain dexamethasone perturbations, we also developed a similar “silver standard” benchmark by asking how differentially expressed genes induced by the drug bortezomib are known members of the apoptosis pathway (*74–76*). Bortezomib is a known inducer of apoptosis across human cancer cell lines. All available signatures induced by dexamethasone and bortezomib perturbations were extracted from all the chemical perturbations Connectivity Mapping resources and ordered based on differential expression significance. The ranked differentially expressed genes for the dexamethasone perturbations were compared to gene sets associated with the NR3C1 transcription factor (TF), and the ranked differentially expressed genes for bortezomib perturbations were compared to apoptosis pathway gene sets. The relationship between bortezomib and apoptosis genes is recovered from all resources better than seen for randomly ordered gene sets, and the Ginkgo Bioworks (*27*, *39*), NIBR DRUG-seq (*38*), and RummaGEO (*22*) gene sets show the most recovery of the expected apoptosis genes (Fig. 3A). The relationship between dexamethasone and NR3C1 was recovered best by the Tahoe-100M (*35*) and the LINCS L1000 (*10*) gene sets, and was not captured in the NIBR DRUG-seq dataset (Fig. 3B). The NIBR DRUG-seq dataset resource exclusively perturbs the U2OS osteosarcoma cell line with dexamethasone, so it is likely that its primary target, NR3C1, is not active in this cell line. This assertion is supported by the low basal expression levels of glucocorticoid receptors in the U2OS cell line (*77*).

**Fig. 3.**
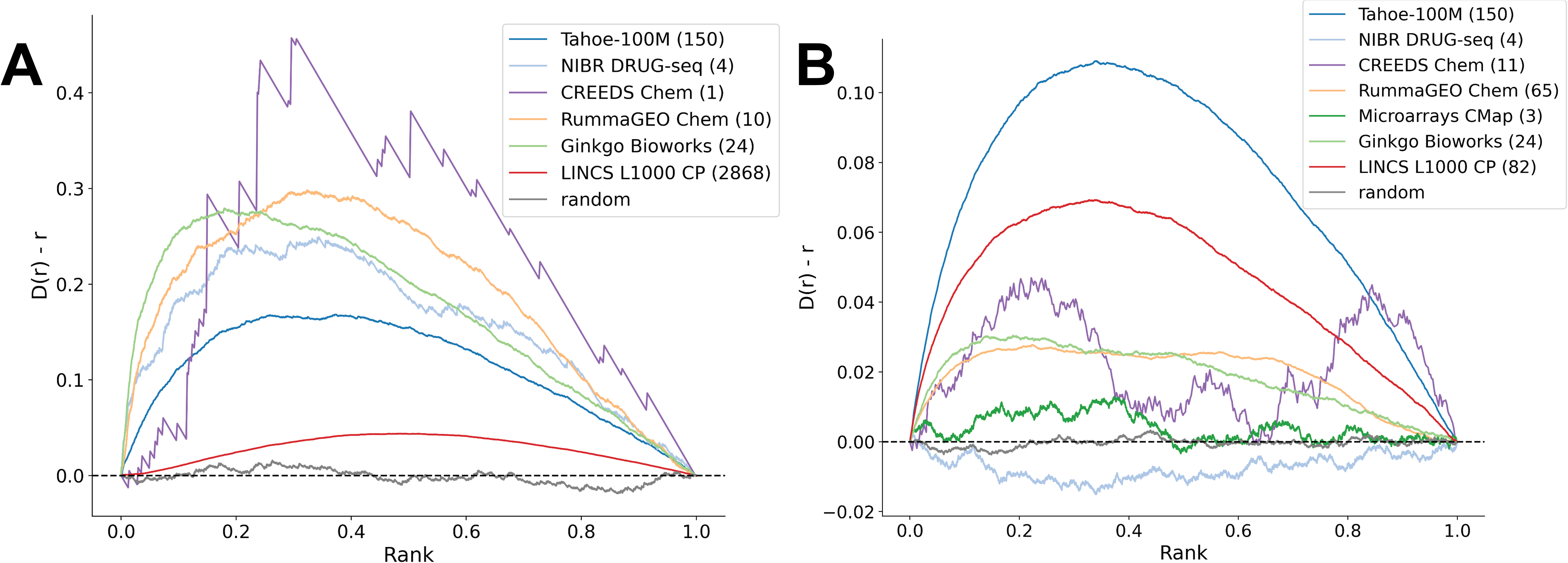
Enrichment analysis results for dexamethasone and bortezomib signatures from Perturb-Seqr datasets against ChEA 2022 NR3C1 TF gene sets for dexamethasone, and Reactome 2024 apoptosis gene sets for bortezomib. (A) Bridge plot of average enrichment of bortezomib signatures against the apoptosis pathway gene sets. (B) Bridge plot of average enrichment of dexamethasone signatures against NR3C1 TF gene sets. Higher peaks indicate better recovery of expected genes.

Since HDAC inhibitors show the most consistent signatures across dosages, time points, cell lines, and gene set libraries, we also developed a similar “silver standard” benchmark for known HDAC inhibitors. Specifically, we compared the consistency of signatures induced by HDAC inhibitors across all the small molecule perturbation Connectivity Maps. Ranked gene sets for HDAC inhibitors from eight of the nine small molecule perturbation resources were enriched against a consensus HDAC inhibitor gene set created from the left-out resource (Fig. S5). The RummaGEO (*22*) HDAC inhibitor consensus gene set generated the highest average enrichment scores across the query HDAC inhibitor gene sets, and the Tahoe-100M (*35*) HDAC inhibitor consensus gene set generated the lowest average enrichment scores (Fig. 4). These results suggest that the HDAC inhibitor gene sets from RummaGEO capture best the global HDAC inhibitors effect, whereas the Tahoe-100M dataset captures a more idiosyncratic response, likely due to the the few cell lines profiled by the Tahoe-100M dataset, and the diversity of cell lines and other conditions captured by RummaGEO.

**Fig. 4.**
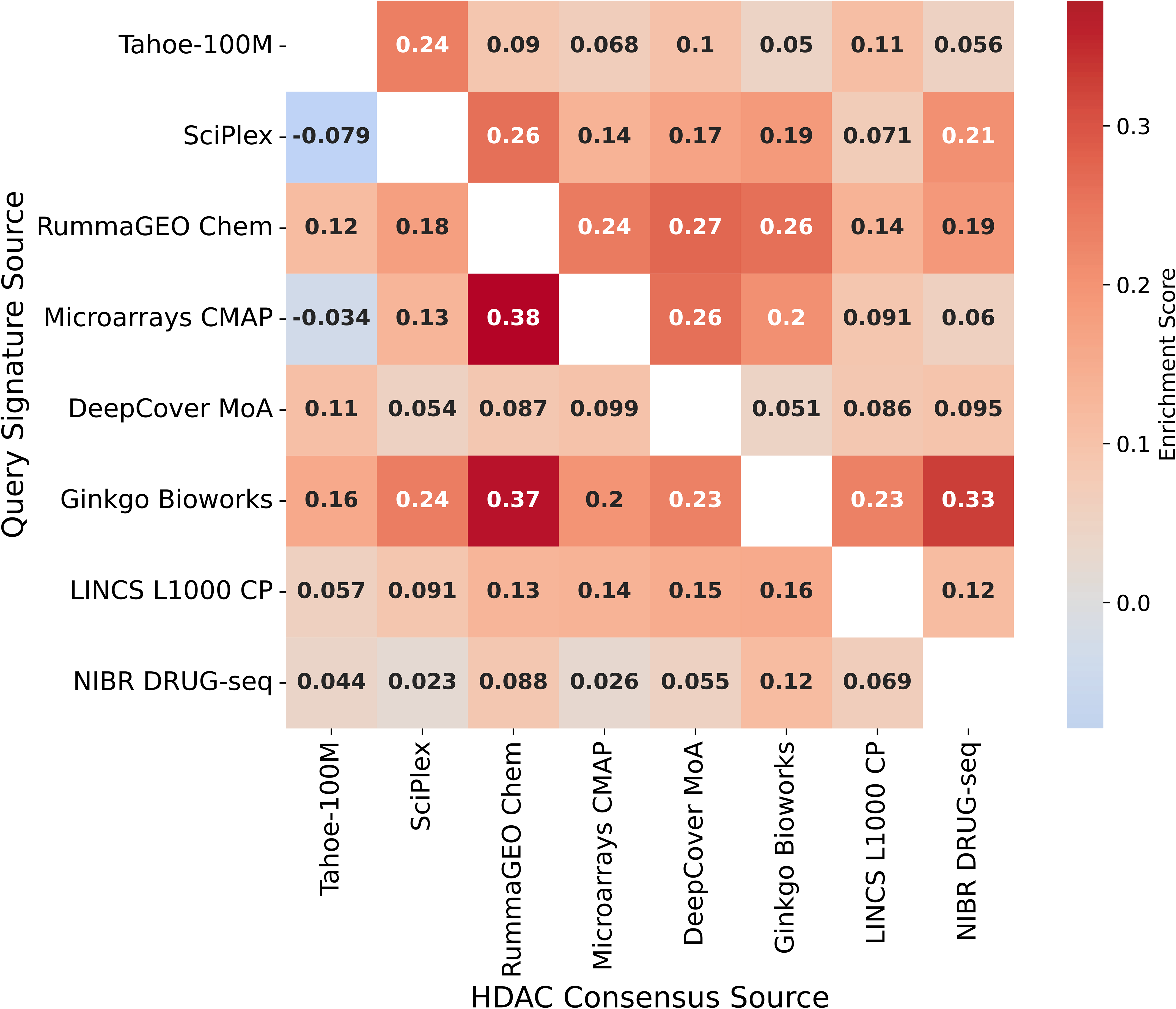
Enrichment analysis for HDAC inhibitors signatures from all drug perturbation Connectivity Maps compared to HDAC inhibitors consensus gene sets generated from a left-out Connectivity Mapping resource.

To benchmark signatures derived from the gene perturbation followed by expression Connectivity Maps, we compared these signatures to gene sets created from the 1,000 most highly correlated genes for each perturbed gene based on mRNA gene-gene co-expression correlations computed from 10,000 randomly selected samples from the ARCHS4 resource (*44*). The ARCHS4 resource provides uniformly aligned samples from all the human and mouse FASTQ files deposited to the Gene Expression Omnibus (GEO) (*21*). Ranked genes from each gene perturbation Connectivity Map recovered the gene sets created from the gene-gene coexpression associations for the same gene (Fig. S6). From across all human genes, the perturbation followed by expression for the gene NPM1 significantly recovers NPM1 co-expressed genes across all the gene perturbation resources except for the LINCS L1000 XPR dataset. NPM1 encodes the nucleophosmin protein, which plays a role in ribosome biogenesis, centrosome deduplication, stress response, and p53-mediated growth suppression (*78*). The role of NPM1, and its gene co-expression module, is well-conserved across cell types. The collection of independent Connectivity Maps can help distinguish genes with consistent transcriptional effects when knocked out from those genes with cell and tissue-specific effects. In addition, consistency across resources, as observed for the NPM1 knockout and co-expressed genes increases confidence of this relationship because it is observed across contexts, assay types, and experiments conducted by different groups.

### Identifying Unknown Drug Targets from Drug-Gene Perturbation Set Crosses

The drug and gene perturbation signatures in the Perturb-Seqr database can be used to discover potentially novel drug-target relationships. To demonstrate this, we crossed gene sets from all the small molecule perturbation libraries with all gene sets from the gene perturbation libraries in Perturb-Seqr. This crossing can uncover pairs of drug/gene perturbation signatures with high similarity. These pairs are referred to as drug-gene mimickers. Cross referencing mimickers with known drug-target associations can confirm that this approach can recover known associations and potentially uncover novel drug targets (Fig. S7). Using this approach with the OpenTargets (*57*) and DGIdb (*79*) resources, we identified known and unknown drug-target associations. The novel mimicking pairs were ranked according to their consistency across resources, and their consistency across conditions such as cell line, dosage, and assay type. Among the top 1,000 mimickers, ARID1A and dasatinib appear at the top of the table (Table S3). ARID1A is part of the SWI/SNF chromatin remodeling complex, whose member genes are mutated in 20% of malignancies (*80*). Specifically, the ARID1A gene is found to be mutated in 6% of all cancer cases (*81*). Experiments in ovarian clear cell carcinoma cell lines showed loss of ARID1A induced sensitivity to dasatinib(*82*). In our analysis, ARID1A knockout, and the knockdown of ARID1A mimicked the effects of dasatinib in a variety of cancer cell lines (Table S4). This suggests that the synthetic lethality of ARID1A is highly similar to the effect of dasatinib, a drug that is likely relevant to treating many cancer types.

In addition to reporting the top mimickers identified in multiple drug-gene pair libraries, we also ranked the top 1000 individual signature crosses with the highest matching overlap, regardless of whether such pair is supported by more than a single drug-gene library pair (Table S5). These individual signature crosses preserve perturbational information such as cell line, dosage, time exposure, and Connectivity Map resource. These crosses were ranked by the product of the Fisher’s p-values for the overlap of the up- and down-regulated gene sets from each signature. We identified small molecule perturbations with the largest number of distinct gene knockout followed by expression mimickers, as well as gene perturbation signatures with the largest number of distinct small molecule mimickers (Fig. S8-S9). A single gene perturbation that induces gene expression changes that are like many different small molecules could indicate that the gene is a hub downstream of many cells signaling pathways. It is also possible that the gene is a bona fide target of many existing drugs, and the compound directly binds to it. These two possibilities are supported by the observation that the gene perturbations with the most matching compounds are well-known cancer targets with known mutations and known existing drugs that bind to them. For example, EGFR, PIK3CA, PIK3R, HSAP9, ADAR, and PHB are some of the gene knockouts that induce transcriptomics changes like many different drug perturbations (Fig. S8). On the other hand, genes such as INTS2, a subunit of the Integrator Complex (*83*), is not a well-documented master regulator. However, its perturbation signatures are like that of many drug perturbations (Fig. S8, Table S6). As a member of the Integrator Complex, which is involved in transcription of many different RNA classes (*83*), INTS2 has broad transcriptional reach and could be a promising new pharmacological target.

### Identifying Functional Communities from Gene-Gene Perturbation Set Crosses

Instead of crossing drug perturbations followed by expression with gene perturbations followed by expression from the various integrated Connectivity Mapping resources, we could also compare only the gene perturbation signatures. Such analysis could lead to the identification of functional modules and new pathways. Employing this approach, we identified several communities of genes whose perturbation causes similar transcriptional changes. These communities can be used to make hypotheses about the membership of genes in undiscovered pathways. When crossing all the LINCS L1000 (*10*) single gene knockout signatures with the Replogle et al. Perturb-seq resource (*29*), nine distinct mutually exclusive communities are identified (Fig. 5). From these communities, we extracted several highly connected subnetworks by only retaining triangles, or in other words, three-node loops (Fig. S10). The genes in the extracted triangle subnetworks (Fig. S10A-B) are enriched for MYC transcription factor targets (*58*), tRNA processing and protein translation. Both subnetworks are significantly enriched for various KEGG pathways (*84*), GO Biological Process terms (*65*, *85*), ChEA TF gene sets (*58*), GWAS Catalog associations (*86*) and MGI Mammalian phenotypes (*87*) related to protein translation and tRNA processing (Table S8). For example, mimicker community 1 (Fig. S10A) is enriched for tRNA translation processes such as the KEGG aminoacyl t-RNA biosynthesis pathway and tRNA aminoacylation GO term (Table S8). Eukaryotic initiation factors such as EIF1AX, EIF2B1, EIF2B2, EIF2B3, EIF2B4, EIF2B5, EIF2S1, EIF2S2, EIF2S3 are also found in this highly connected subnetwork. Although these are not annotated as tRNA translation genes, it is known that EIF2 and tRNAs interact in translational processes and are reprogrammed under stress conditions (*88*). Hence, the perturbational mimicker subnetworks capture known pathways and can discover new members of such pathways or even entirely new pathways. For example, mimicker community 2 (Fig. S10B) is enriched for ribosomal biogenesis processes (Fig. S10B and Table S8). Highlighted in this subnetwork is C12ORF45. C12ORF45 knockout mimics the transcriptomics changes induced by RSL24D1 and XPO1 knockouts, suggesting that C12ORF45 may be functionally related to them. This hypothesis is partially supported experimentally, because C12ORF45 has been shown to bind to NOP58, which is a core component of small nucleolar ribonucleoprotein (snoRNP) complexes involved in ribosome biogenesis (*89*). However, more targeted experiments are needed to further confirm the unknown functions of C12ORF45.

**Fig. 5.**
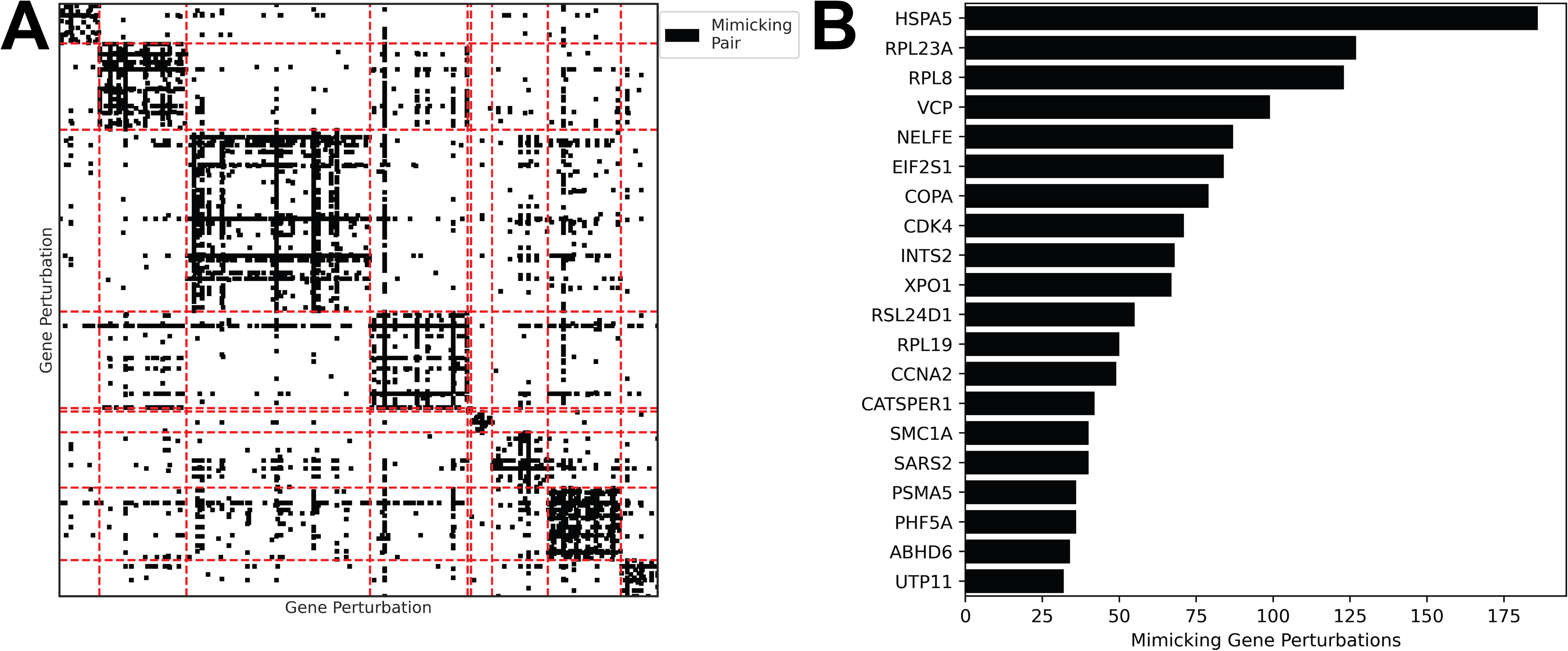
Mimicker gene perturbation modules from crossing LINCS L1000 XPR and Replogle et al. gene perturbation followed by expression gene set libraries. (A) Densely connected regions of mimicking pairs, grouped using the Louvain community detection algorithm on the adjacency matrix of the top 5000 mimicking pairs by significance of up and down-regulated gene set overlap. Full Enrichr enrichment analysis results for each community are available as links provided in Table S7. (B) Most commonly mimicked gene perturbations.

### Using the Perturb-Seqr Search Engine and Website

All the gene expression signatures assembled for Perturb-Seqr are served for enrichment analysis through the Perturb-Seqr signature search engine (Fig. 6). Through a web interface, users can submit either paired up- and down-regulated gene sets, or a single gene set, to identify perturbation signatures like their query (Fig. 6A). Users can restrict the search to only search against a subset of the 16 libraries included in Perturb-Seqr (Fig. 6A). The search engine results are returned to the user as per-library interactive butterfly charts and as paginated tables (Figs. 6B-C). The interactive butterfly charts enable visual comparison of the most similar and opposite perturbations across all libraries by displaying the top significant reversers and mimickers for paired gene set input, or the top up- and down-regulated matches for single gene set input. Complementary to the butterfly charts, paginated tables enable granular exploration of the results by reporting the most similar or opposite perturbations along with their library name, cell line, time point, concentration, directionality, mechanism of action, FDA-approval status, gene set size, overlap size with the query gene set, overlapping genes, and enrichment statistics. Within each table, compounds are linked to PubChem (*90*) and perturbed genes are linked to their NCBI Gene entries (*91*), Harmonizome (*43*), ARCHS4 (*44*), GDLPA (*92*), and GSFM (*93*).

**Fig. 6.**
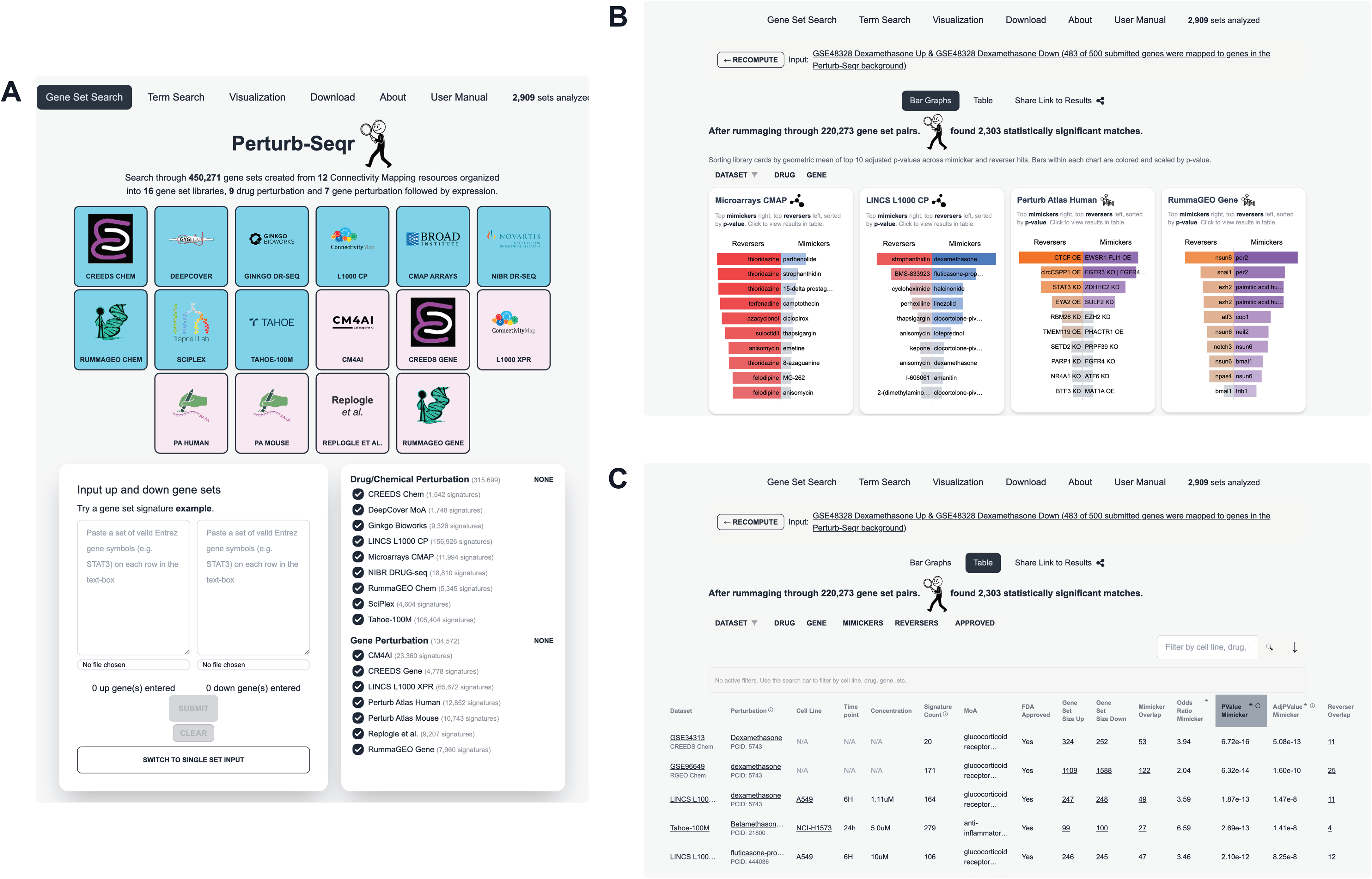
The Perturb-Seqr web-based search engine application. (A) The Perturb-Seqr web-based application landing page accepts input gene sets and filtering options. (B) Interactive butterfly plots assist users to quickly identify top mimickers and reversers across Connectivity Mapping resources. C) Combined tabular view of Connectivity Mapping results. These tables are available for download and display granular information about the matching perturbations.

Additional filtering can be applied to restrict the search to perturbations of interest, such as only searching for drug or gene perturbations, only listing FDA-approved drugs in the results, as well as searching the results to find specific terms. Multiple free-text terms can be combined, for example, to only list results for specific cell lines at once. In addition to the gene set search, a term search page is provided with the same filtering options. In this interface users can submit text-based queries and receive a table of only the signatures that match the queried term and other applied filters. Both gene set and term search results can be downloaded from the paginated tables as Tab-Separated Values (TSV) files. By clicking the gene set overlap size, users can explore the overlapping genes between a Pertrub-Seqr signature, and the user submitted query gene set. Users can copy these overlapping genes as gene symbols to their clipboard, or submit the overlapping genes directly to Enrichr (*59*), Rummagene (*67*), RummaGEO (*22*), or GSFM (*93*). UMAP (*63*) visualizations of the 16 gene set libraries within Perturb-Seqr can be explored on a dedicated data visualization page. Source data for each gene set library is made available for download as compressed Gene Matrix Transposed (GMT) files, or as full genome-wide differential expression signatures. The gene set and term search functions are also available programmatically through a GraphQL API and an accompanying Python package. In addition, Perturb-Seqr is also provided as an agent skill markdown file, and as a Model Context Protocol (MCP) server.

### Use Case #1: Mimickers of Endurance and Resistance Exercise Signatures from the MoTrPAC Adult Acute Exercise Study

Exercise has been shown to mitigate the disease burden of diabetes and various other metabolic disorders as well as provide health benefits at the molecular and systemic levels (*94*). While physical activity may be an effective intervention, there are people that cannot exercise, and it is possible that exercise mimetics will improve the overall health status of a person regardless of whether they exercise or not. The Molecular Transducers of Physical Activity Consortium (MoTrPAC) (*31*) is an NIH Common Fund program that has generated multi-omics data that captures the molecular response to endurance and resistance exercise in a group of 175 normally sedentary adults (*30*) and aerobic exercise of rats (*95*). For this use case, exercise-induced up and down gene sets were extracted from tables of differentially abundant molecular features downloaded from the MoTrPAC data portal. Differential abundance tables were derived from RNA-seq data generated from skeletal muscle tissue biospecimens of study participants collected 3.5 hours after an acute or endurance bout of exercise. The extracted up and down gene sets represent changes observed because of exercise compared to a non-exercise control group. These gene sets were submitted to Perturb-Seqr to find chemical and gene perturbation mimickers of exercise (Fig. 7).

**Fig. 7.**
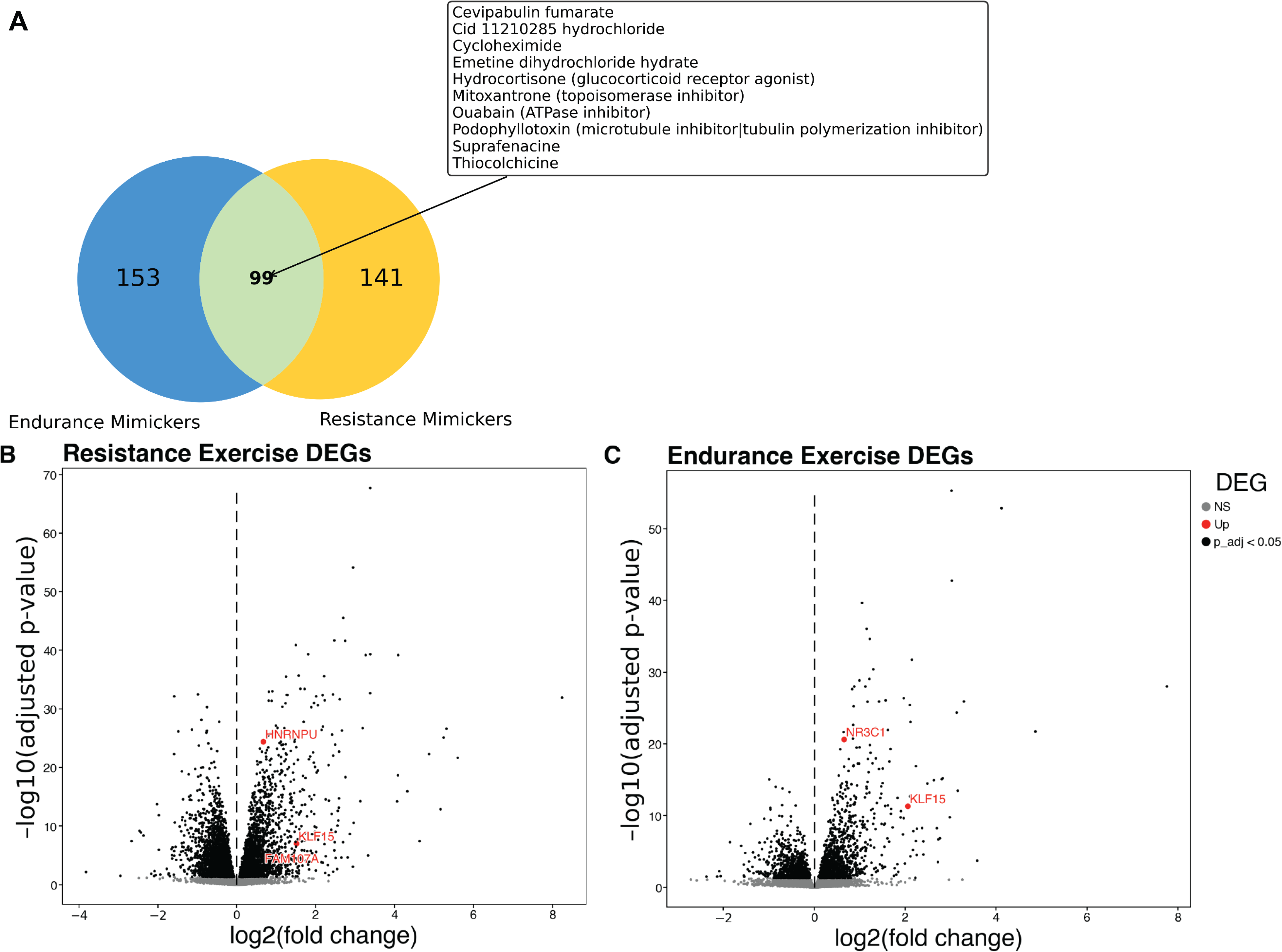
Chemical mimickers of endurance and resistance exercise-induced signatures in muscle samples from the MoTrPAC adult acute exercise study. (A) Overlap among significant chemical mimickers of the endurance and resistance exercise signatures. The 10 most significant overlapping mimickers by the product of Fisher’s exact test p-value are highlighted. (B) Differentially abundant genes in the muscle samples taken 3.5 hours after the exercise period for the resistance exercise vs. control group; and the (C) endurance exercise vs. the control group. Genes in the GO “Cellular Response to Glucocorticoid Stimulus” gene set (GO:0071385) are highlighted in both volcano plots.

The glucocorticoid receptor agonist hydrocortisone is among the most significant mimickers of the MoTrPAC acute endurance and resistance exercise gene signatures. Hydrocortisone is an FDA approved cortisol replacement treatment. Cortisol is a glucocorticoid that increases after acute physical exercise (*96*). In addition to the observation that glucocorticoids can be molecular by-products of exercise, treatment with dexamethasone, another glucocorticoid receptor agonist, increased the exercise capacity of mice by increasing the expression of the metabolic transcription factor KLF15 (*97*). KLF15 is up-regulated in the endurance and resistance exercise signatures in muscle as well as other genes in the GO “Cellular Response to Glucocorticoid Stimulus” pathway (GO:0071385) (Fig. 7B-C). The expression patterns observed and the subsequent Perturb-Seqr analysis suggest that hydrocortisone could be further investigated as an exercise mimicker. In addition, many other pre-clinical compounds and approved drugs can be explored, as well as gene knockouts as potential new targets (Table S9). It should be noted that it is likely that many of these mimickers may not be relevant in-vivo since the data in Perturb-Seqr is collected in cell lines. Moreover, the concentration and time points may differ. Finally, exercise appears to induce an inflammatory response that may not be beneficial to be exacerbated with drugs.

### Use Case #2: Reversing Nf-kB Pathway Activation in the CPTAC3 Lung Squamous Cell Carcinoma Cohort

Lung squamous cell carcinoma (LSCC) accounts for 20% of primary lung neoplasms in the United States (*98*). We used Perturb-Seqr to identify chemical reversers of an NF-kB activation signature derived from the LSCC cohort in the CPTAC-3 project, accessed from the database of Genotypes and Phenotypes (dbGaP accession phs001287.v1.p1) (*99*). NF-kB activation promotes tumorigenesis in multiple malignancies and is associated with chemoresistance (*100*). The NF-kB activation signature was created by comparing the bulk RNA-seq profiles between high and low NF-kB PROGENy score (*101*) groups in the CPTAC-3 LSCC cohort (Fig. 8A). To identify chemical and gene perturbations that reverse the NF-kB PROGENy score signature, the up- and down-regulated genes were submitted to Perturb-Seqr via the paired gene set query.

**Fig. 8.**
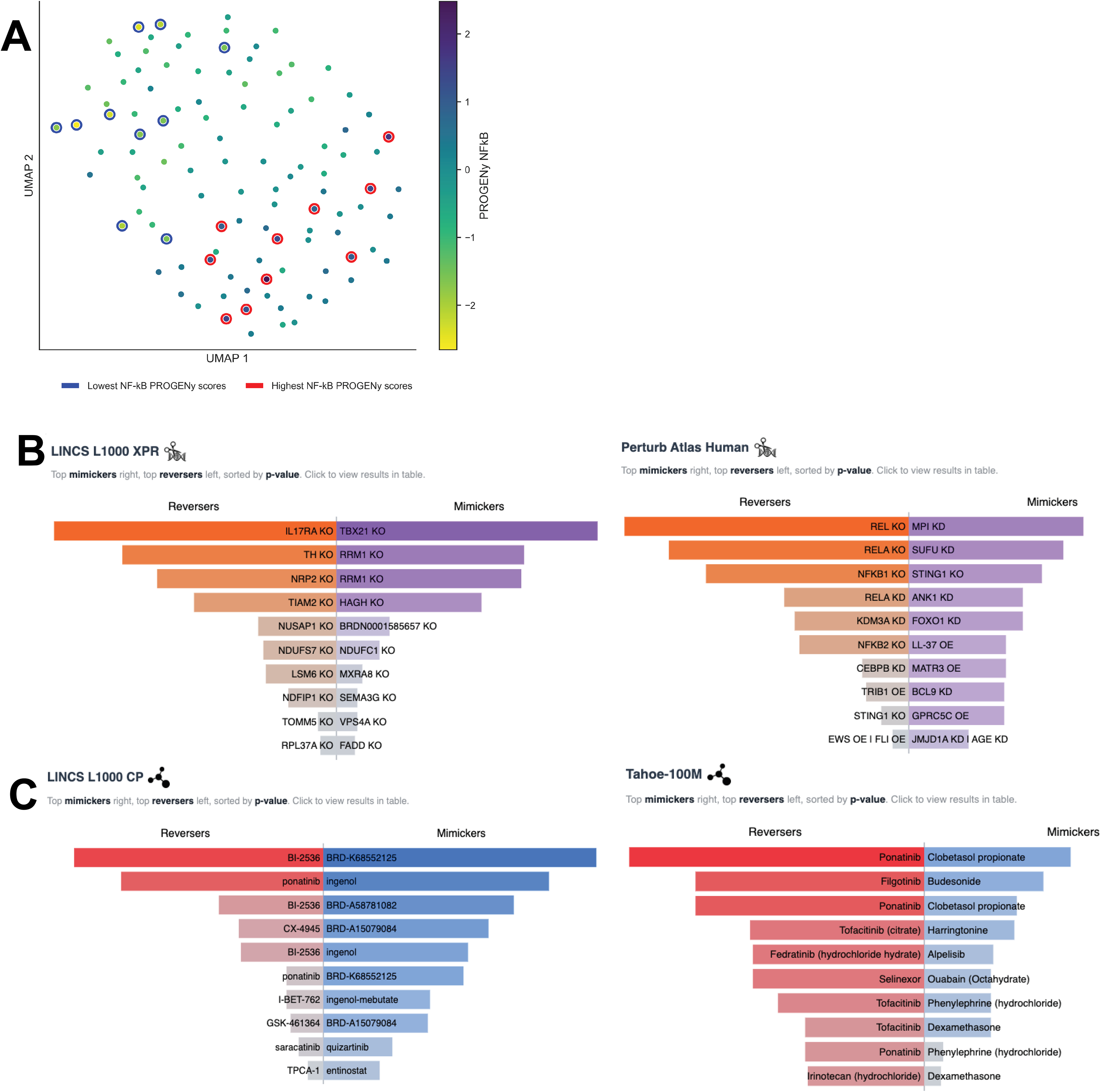
Small molecule and gene perturbagens that reverse the NF-kB activation signature in the CPTAC3 LSCC cohort. (A) PROGENy NF-kB activation scores for CPTAC3 lung squamous cell carcinoma cohort. Transcriptomics profiles of samples with the highest and lowest NF-kB activation scores were compared to create the activation signature. (B) Top gene perturbation reversers of the NF-kB activation signature returned from a Perturb-Seqr query. (C) Top drug perturbation reversers.

The most significant reversing gene perturbations are canonical NF-kB genes and IL-17 knockouts (Fig. 8B). IL-17 has been shown to induce the epithelial to mesenchymal transition in the A549 lung cancer cell line partially through NF-kB activation (*102*). Biologics that target IL-17 have been developed for the treatment of inflammatory diseases like psoriasis and arthritis (*103*), and could be repurposed for LSCC. Among the most significant chemical reversers of the NF-kB activation pattern was Ponatinib, which was identified in multiple Connectivity Mapping resources as a top highly significant reverser (Fig. 8C). Ponatinib is a tyrosine kinase inhibitor that targets the growth factor receptor FGFR1 (*104*) . FGFR1 is highly expressed in lung cancers (*105*), suggesting that Ponatinib might be an excellent candidate for drug repurposing for LSCC.

## Discussion

The Perturb-Seqr signature search engine contains a comprehensive collection of gene set signatures extracted from publicly available Connectivity Mapping resources. The collection includes 450,271 gene sets that capture 47,499 unique experimental perturbations of either small molecules or single genes. Leveraging the diversity of this integrated Connectivity Map, we examined perturbational consistency, identified known and new macromolecular pathways, and discovered unestablished drug-gene and gene-gene associations. Comparing perturbation of the same drug or gene across conditions and platforms reveal significant differences across dosages, time exposures, and cell lines for most perturbations. However, consistency is also observed for few compounds and knockouts. This suggests that these perturbations are cell type, concentration, and dosage agnostic. Furthermore, we observed that each Connectivity Mapping resource captured distinct biological relationships. Gene perturbation Connectivity Maps recovered different relationships captured by the ARCHS4 gene-gene co-expression matrix (Fig. S6) and drug perturbation Connectivity Maps recovered different known relationships between drugs, their targets, and the target genes of the known drug targets (Fig. 3). The inconclusive results of the benchmarks suggest that the different resources may contribute different aspects of real biology, and each platform also has some technical biases. Models that could transform the data from one assay type to another (*106*) may be able to separate the experimental noise from the biological variability. Foundation models trained on diverse datasets such as GSFM (*93*), BulkFormer (*107*) and Nicheformer (*108*), could be used to learn biological relationships among genes and perturbations across conditions. Such models achieve better performance when trained on large collections of diverse experimental conditions and assay technologies.

In the future, we plan to reuse the collection of the Connectivity Maps in Perturb-Seqr to train foundation models. Such foundation models could be applied to a variety of tasks, including predicting the effects for unseen conditions such as cell lines, concentrations, and time points; and predicting the synergistic effects of drug-drug, drug-gene, and gene-gene combinations. Currently, many models that predict the effects of unseen perturbations on gene expression do not outperform a linear baseline (*109*). The diverse assay technologies, cellular contexts, and perturbations collected and organized for Perturb-Seqr could offer improvement in this space. A foundation model created from multiple Connectivity Maps could also be used to create digital twin cellular models. Dynamical models that capture the response to perturbations can potentially replace animal and organoid testing of pre-clinical compounds and new industrial chemicals (*110*). Many of the Connectivity Map resources harmonized for creating Perturb-Sqr focus on therapeutics and cell lines relevant to cancer (Fig. 1). The prevalence of the cancer-focused context may impact the ability of Perturb-Seqr and foundation models trained using the Perturb-Seqr collection may not generalize well to other diseases. While the focus on cancer genetics could reflect research priorities in the field, we hope to increase the diversity of conditions captured in Perturb-Seqr as additional Connectivity Mapping resources become available.

## Data and Code Availability

The Perturb-Seqr signature search engine website provides access to gene set matrices, differential gene expression tables, and perturbation metadata. Perturb-Seqr can be found at https://perturbseqr.maayanlab.cloud. The search engine and website source code are available on GitHub at: https://github.com/MaayanLab/PerturbSeqr. The Perturb-Seqr search engine can be integrated into AI applications through an MCP server that exposes Perturb-Seqr’s enrichment and database query functionality as agent-callable tools. The MCP server is hosted at https://perturbseqr.maayanlab.cloud/mcp and can be installed locally from https://github.com/MaayanLab/PerturbSeqr/tree/main/mcp. An agent skill file that provides agents with instructions on how to use the Perturb-Seqr graphQL API effectively is available from the same repository.

## Acknowledgements

This work was partially funded by NIH grants OT2OD036435, U24CA264250, R01DK131525, and U24CA271114.

