## Supplementary Figures and Tables for "Perturb-Seqr: Signature Search Engine Integrating Connectivity Mapping Resources"

[*]Contributed equally

### Supplemental Figures

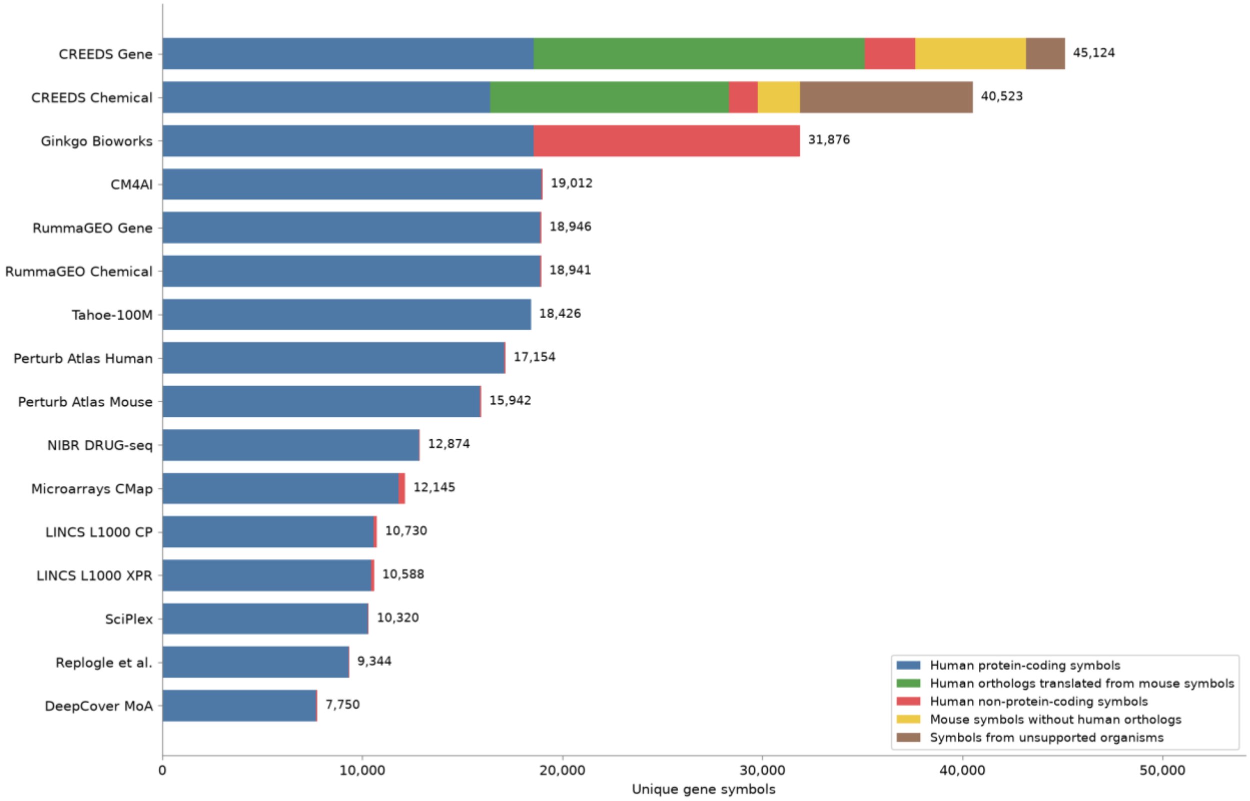


**Fig. S1.** Unique gene identifiers in each Connectivity Mapping resource. Gene sets are filtered to retain only human protein-coding genes. Mouse symbols are translated into their human orthologs using the Mouse Genome Informatics (MGI) homology table. All other species are dropped.

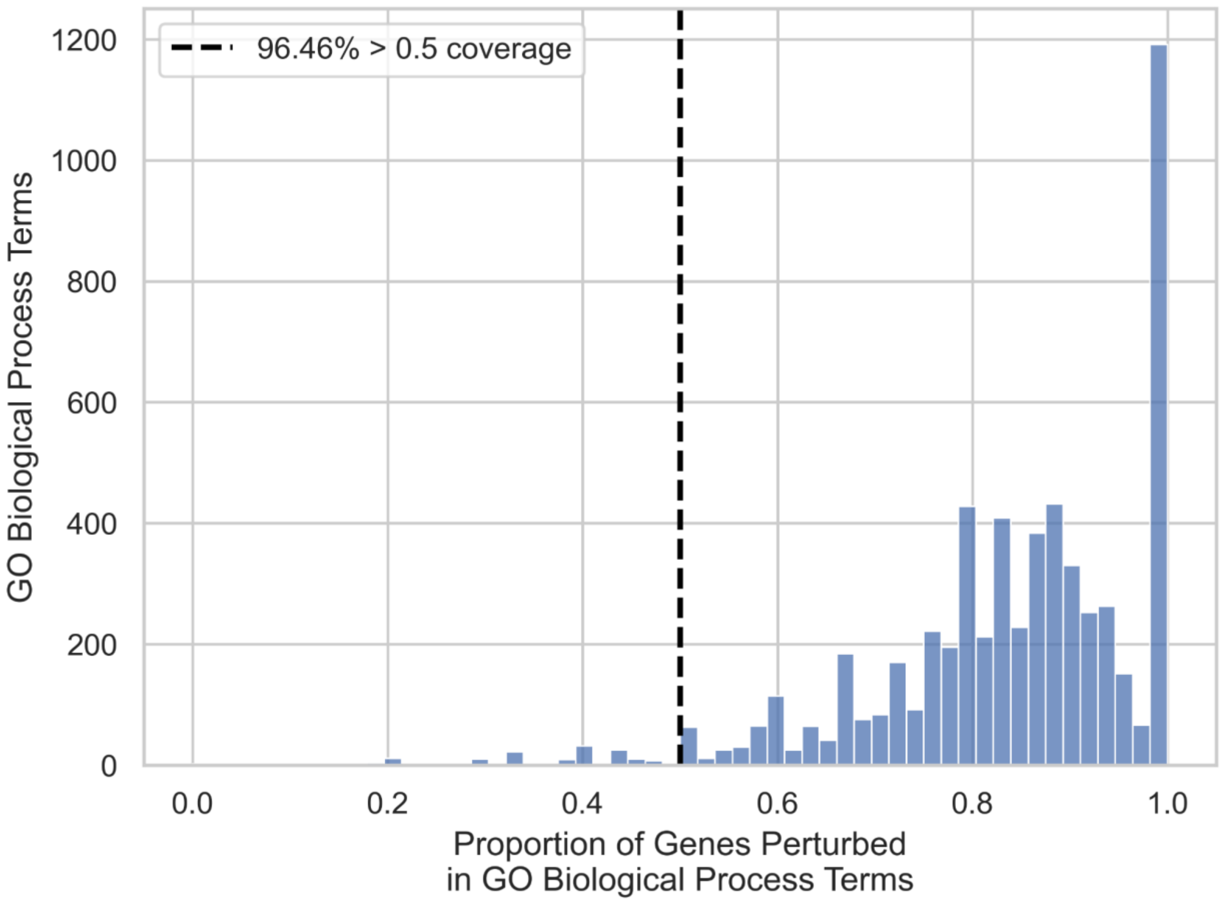


**Fig. S2.** Coverage of GO Biological Processes (BP) by single gene perturbation signatures in Perturb-Seqr. For each term in the GO BP library, coverage was calculated as the fraction of the term's genes that are the target of at least one single gene perturbation in Perturb-Seqr. The distribution of coverage across all GO BP terms is shown, and more than 95% of terms have at least half of their genes represented by one or more perturbation signatures in the Perturb-Seqr collection. GO BP terms were extracted from the 2026 version of the GO BP library in Enrichr.

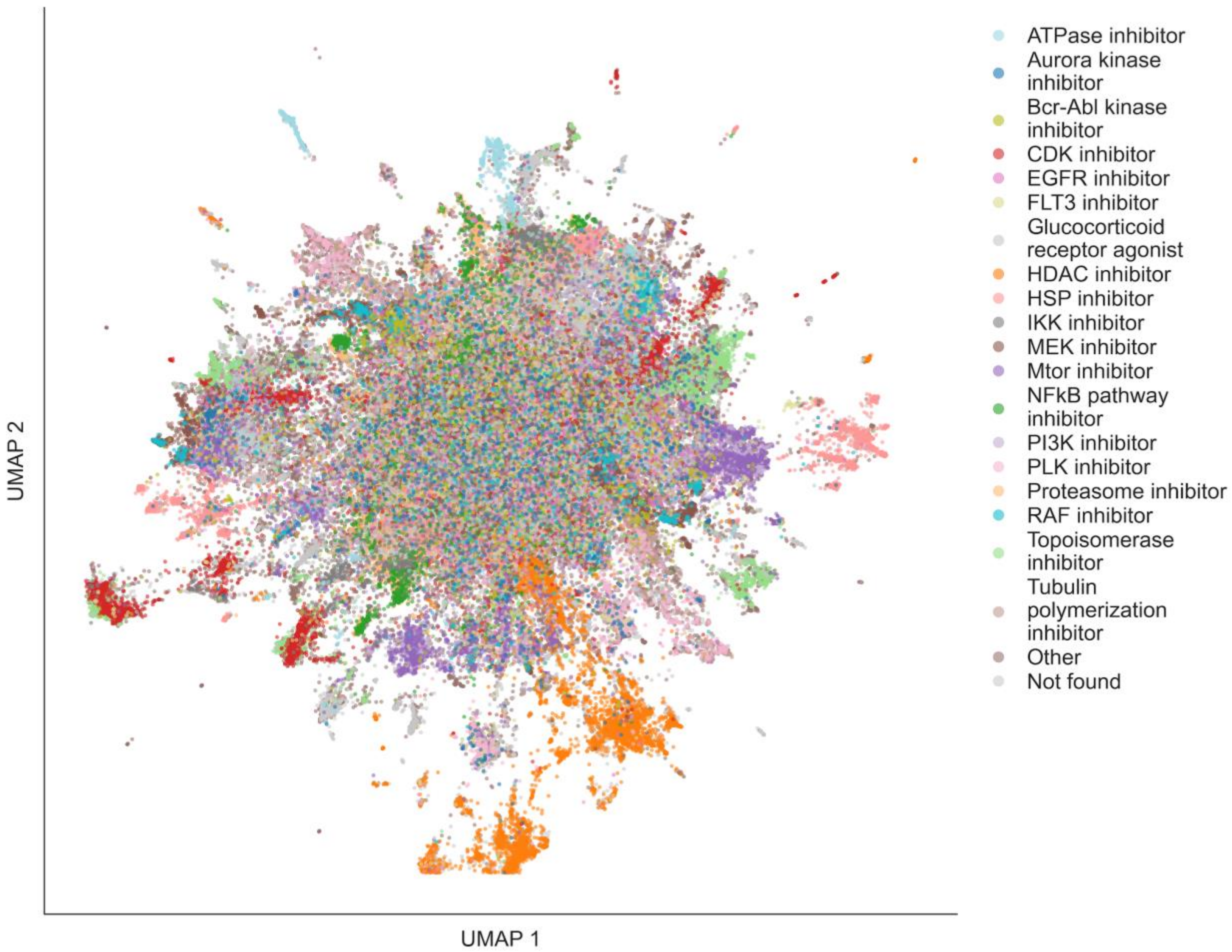


**Fig. S3-A1. LINCS L1000 Chemical Perturbation (MoA)**

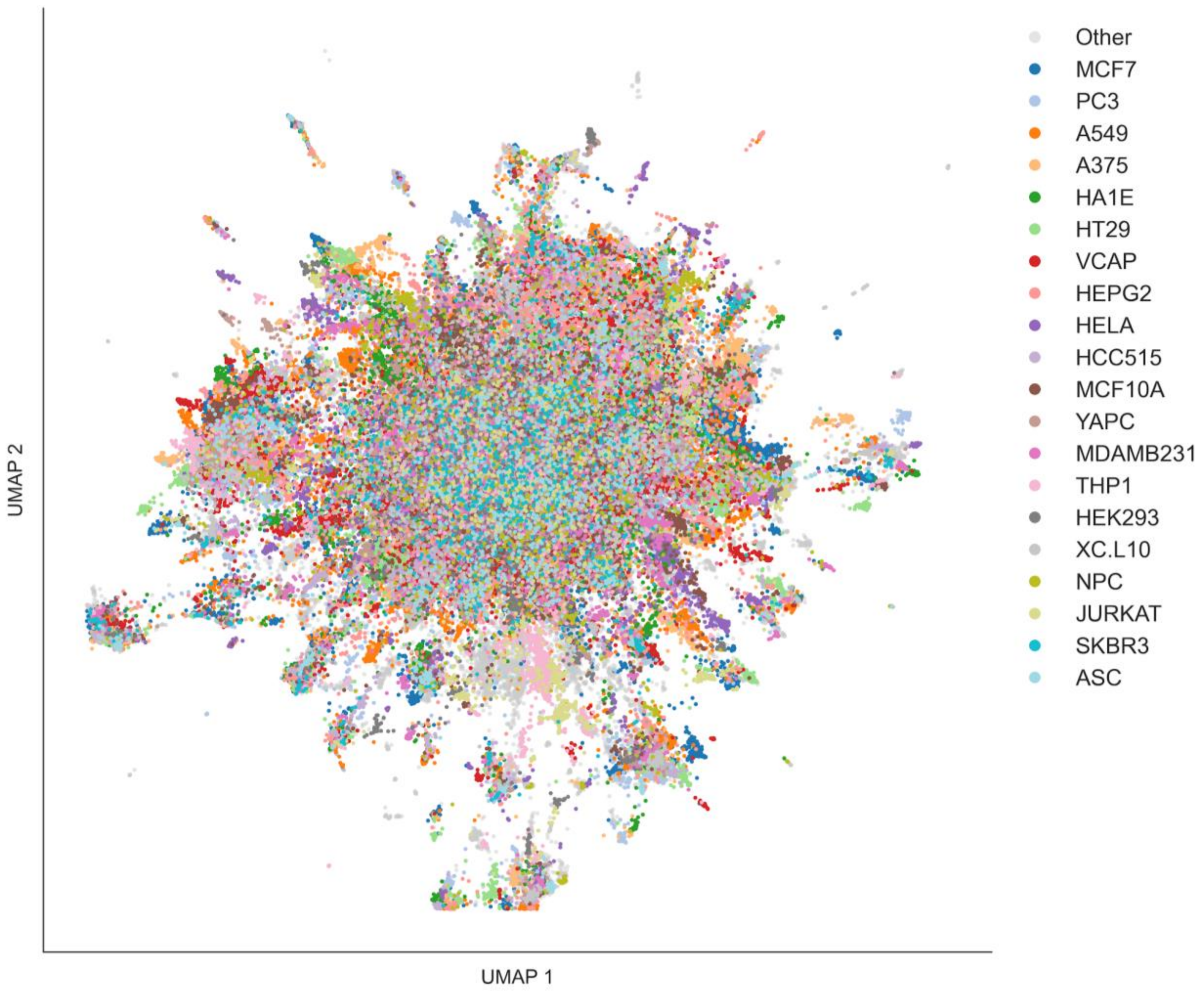


**Fig. S3-A2. LINCS L1000 Chemical Perturbation (Cell Line)**

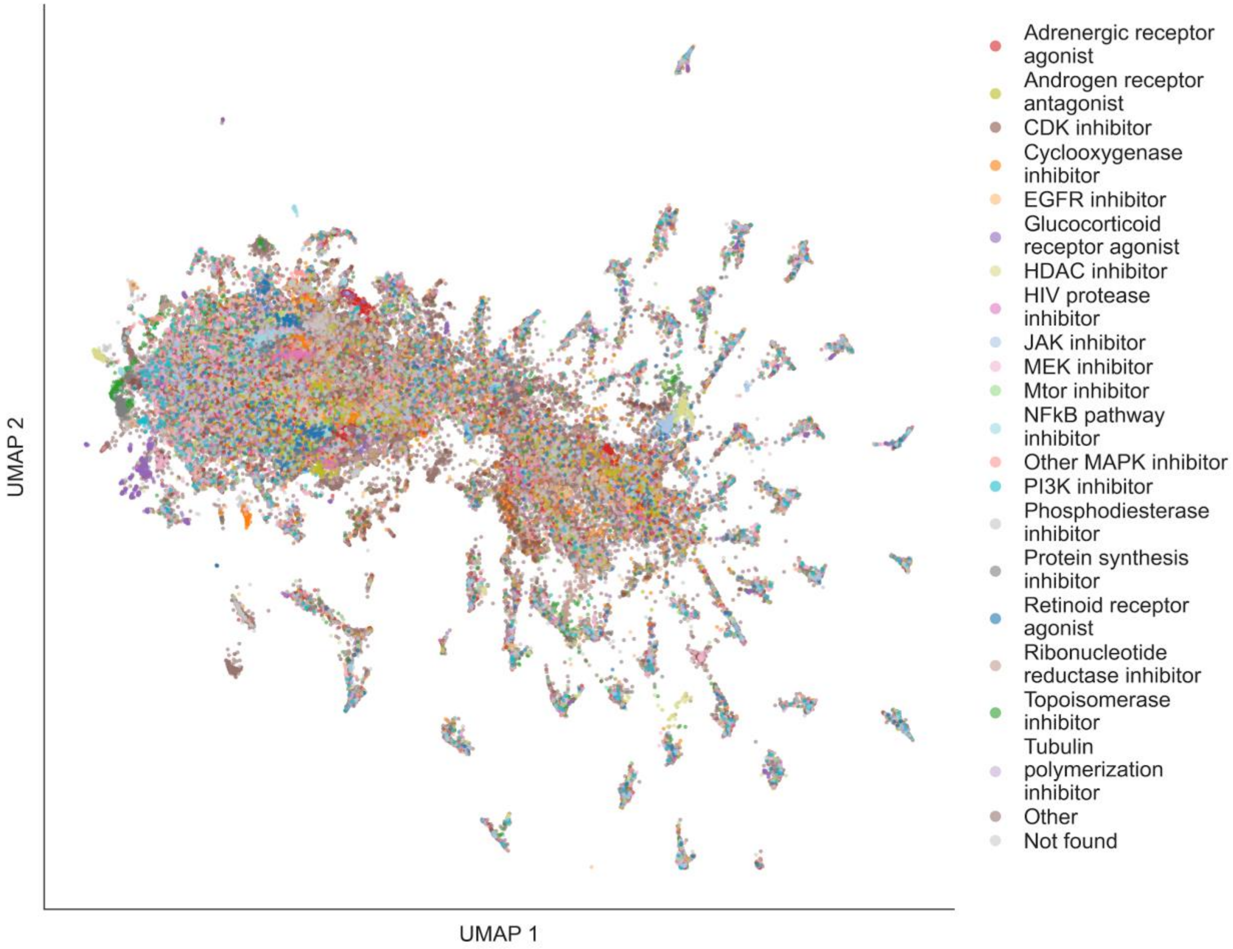


**Fig. S3-B1. Tahoe-100M (MoA)**

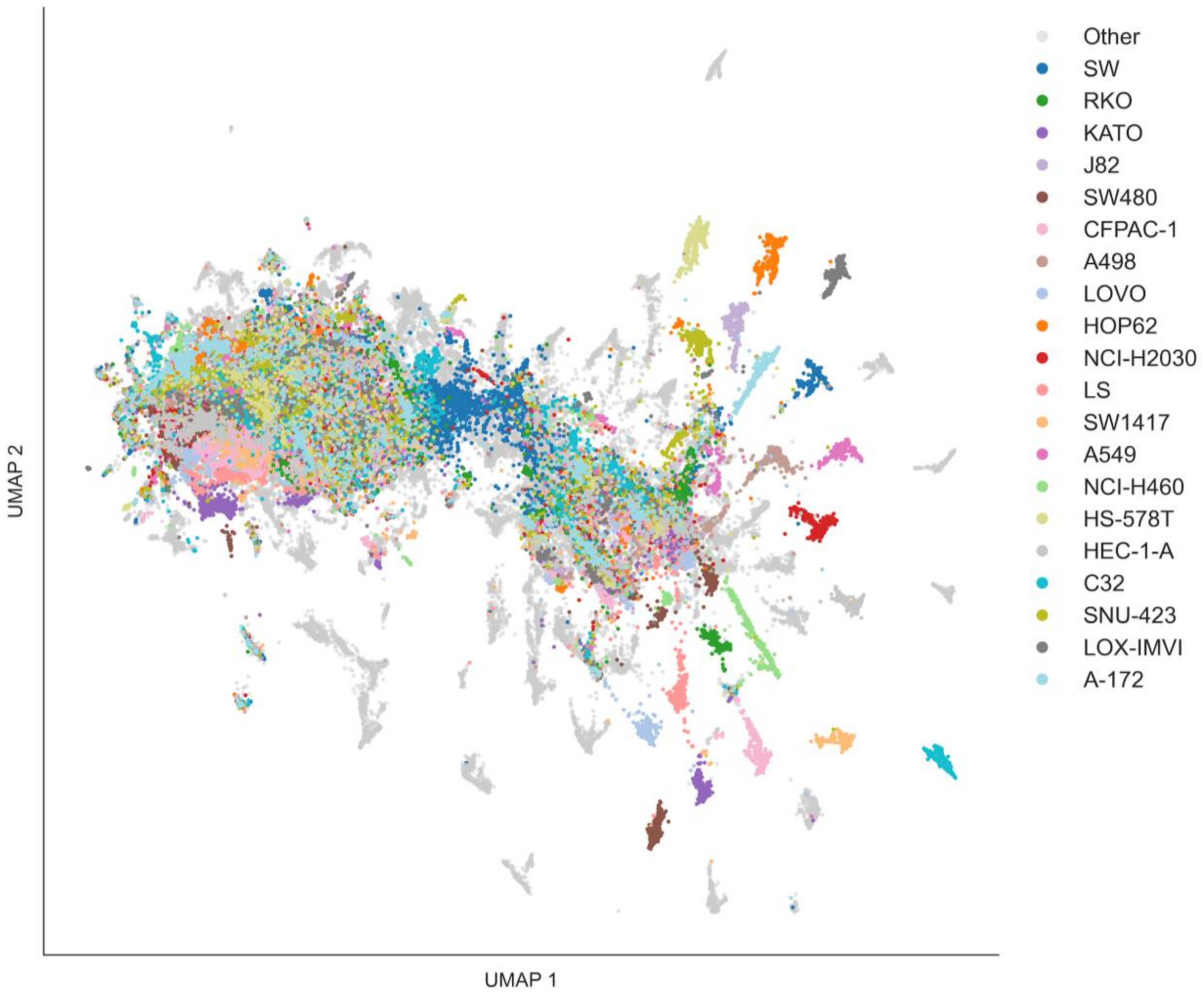


**Fig. S3-B2. Tahoe-100M (Cell Line)**

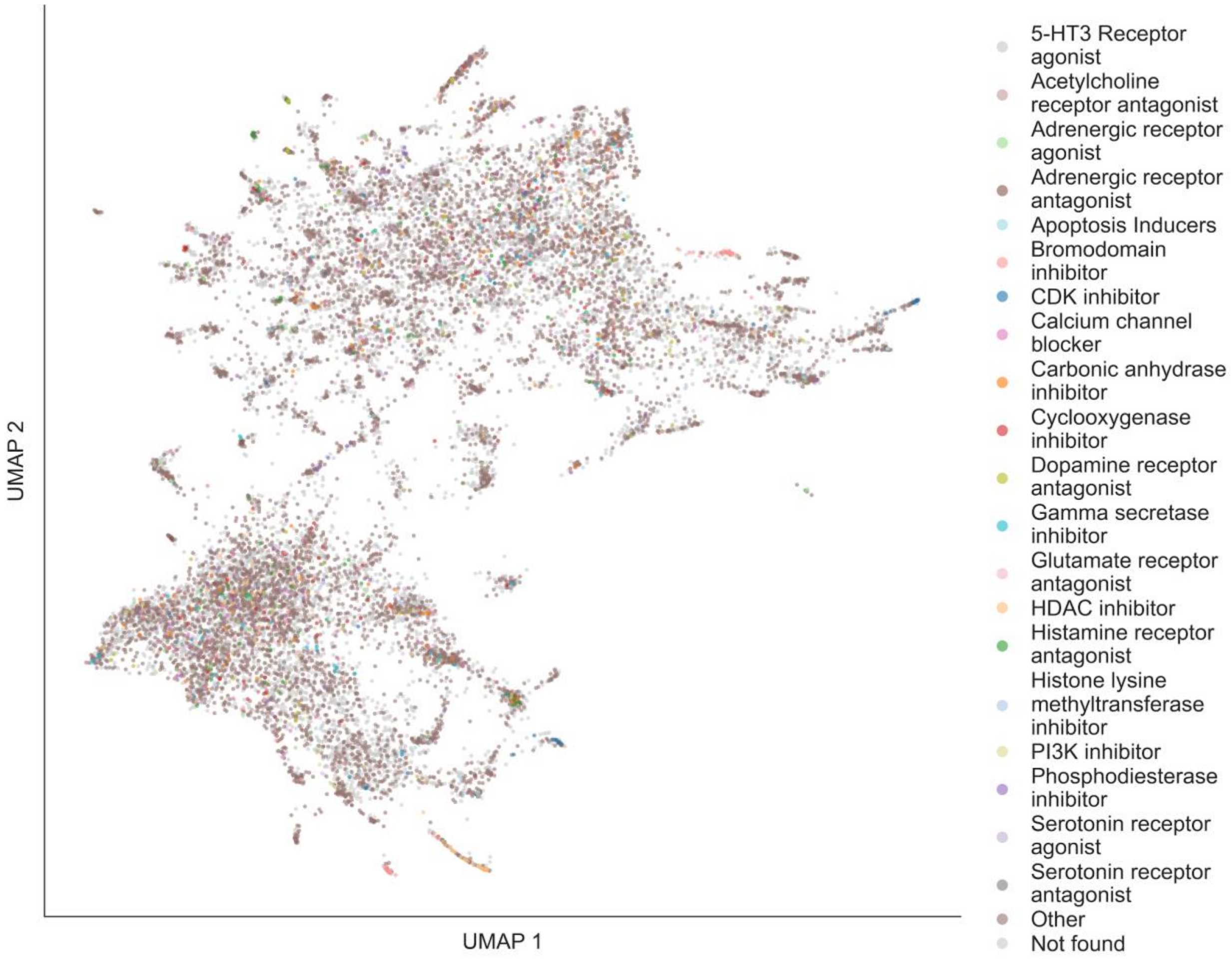


**Fig. S3-C1. NIBR DRUG-seq (MoA)**

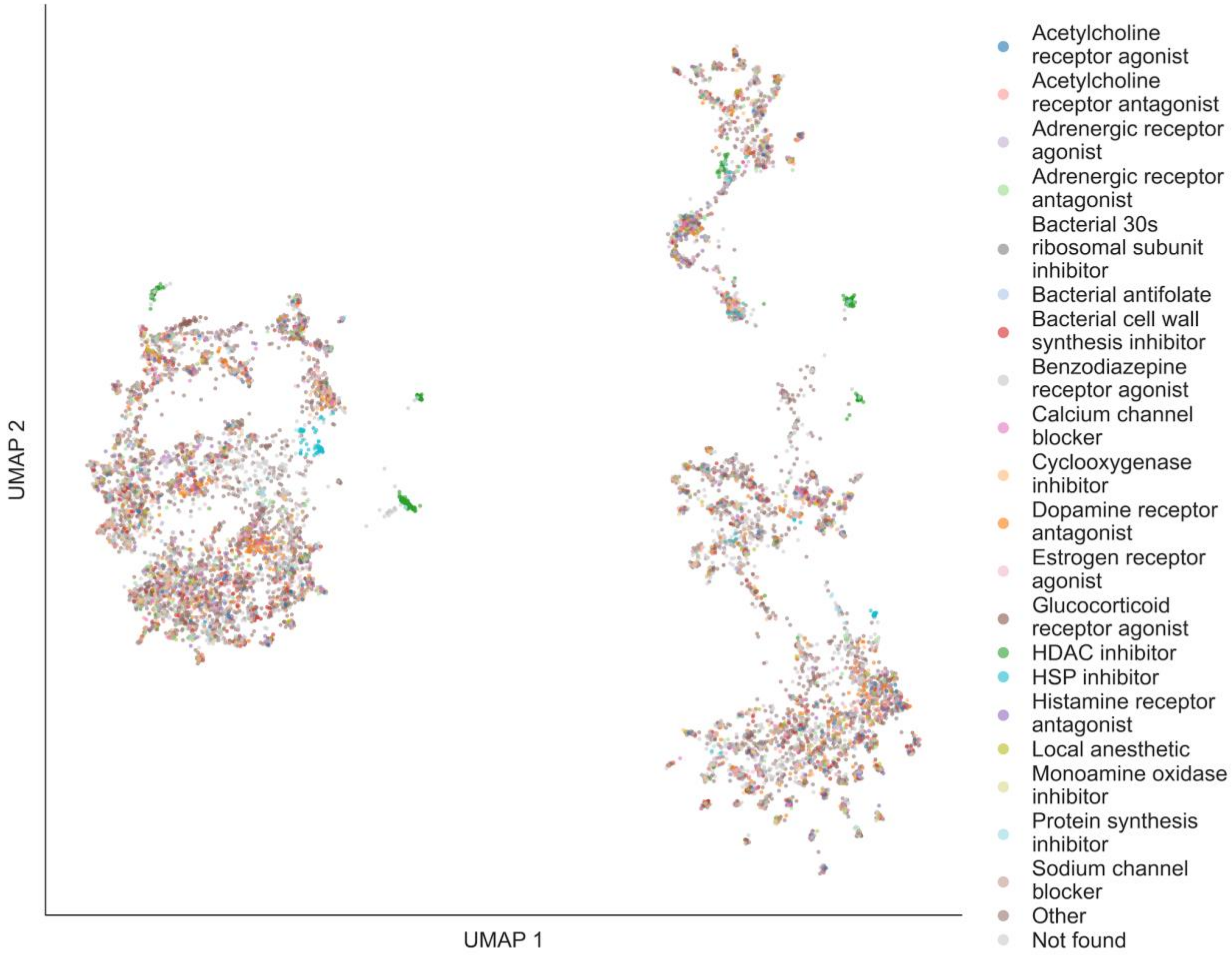


**Fig. S3-D1. Microarrays CMAP (MoA)**

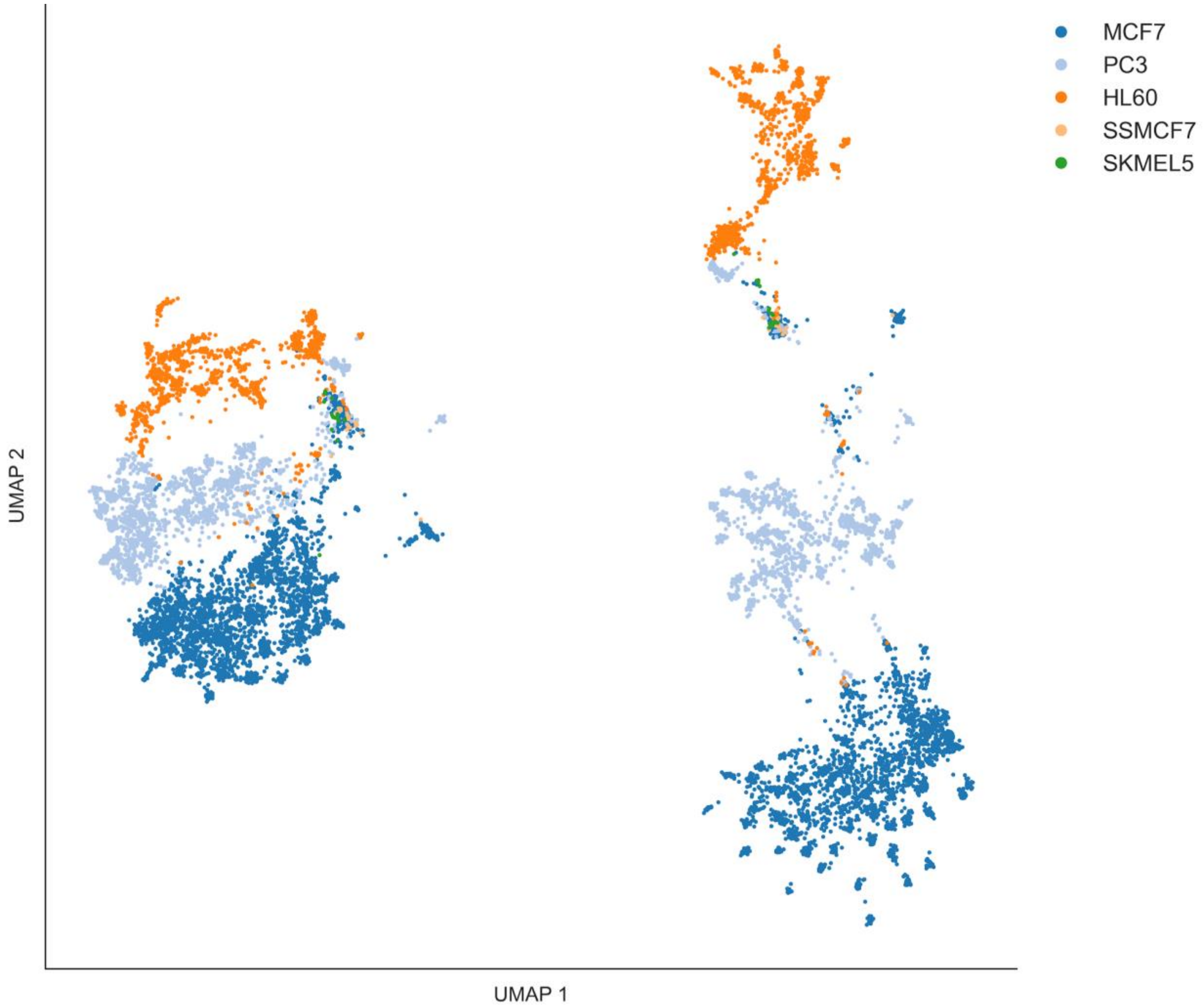


**Fig. S3-D2. Microarrays CMAP (Cell Line)**

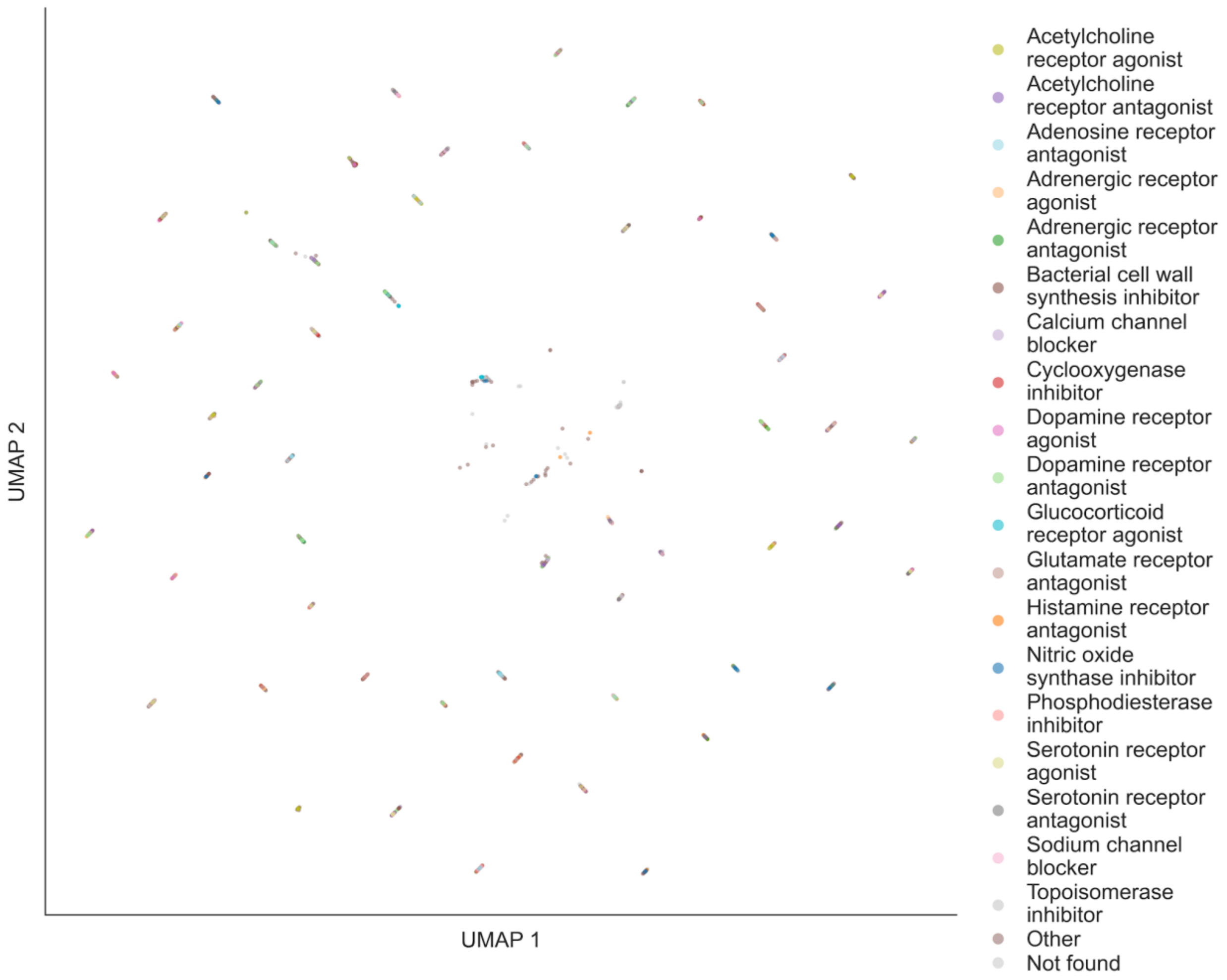


**Fig. S3-E1. Ginkgo Bioworks GDPx1 (MoA)**

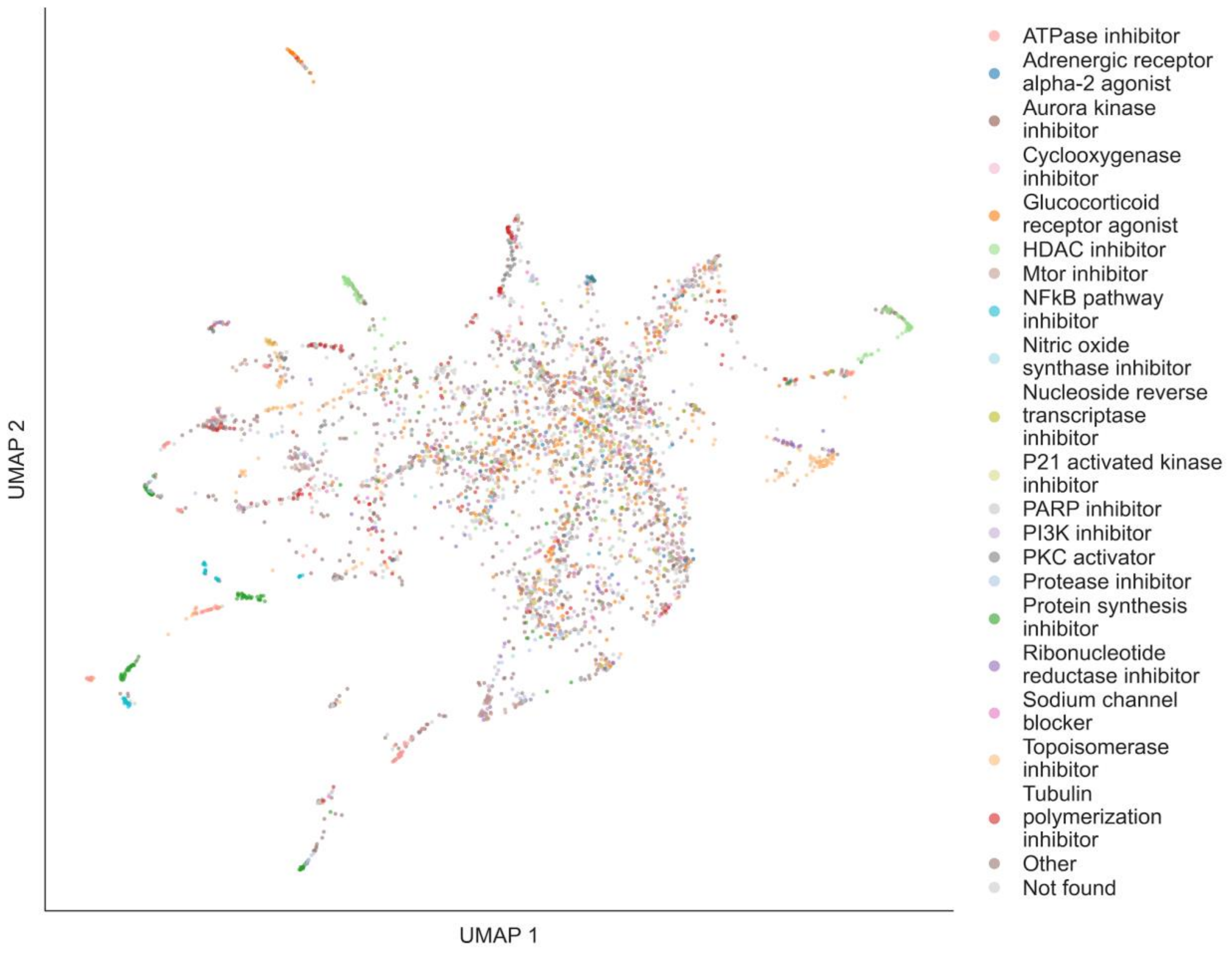


**Fig. S3-E2. Ginkgo Bioworks GDPx2 (MoA)**

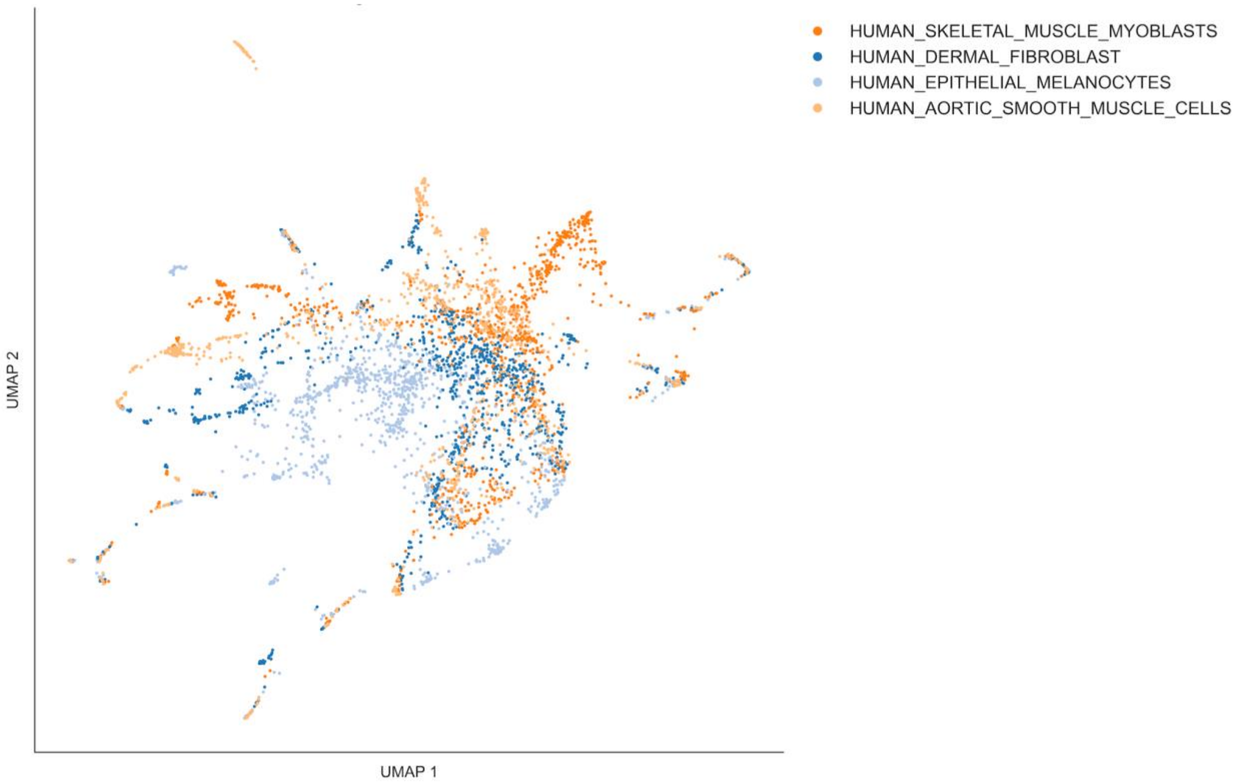


**Fig. S3-E3. Ginkgo Bioworks GDPx2 (Cell Line)**

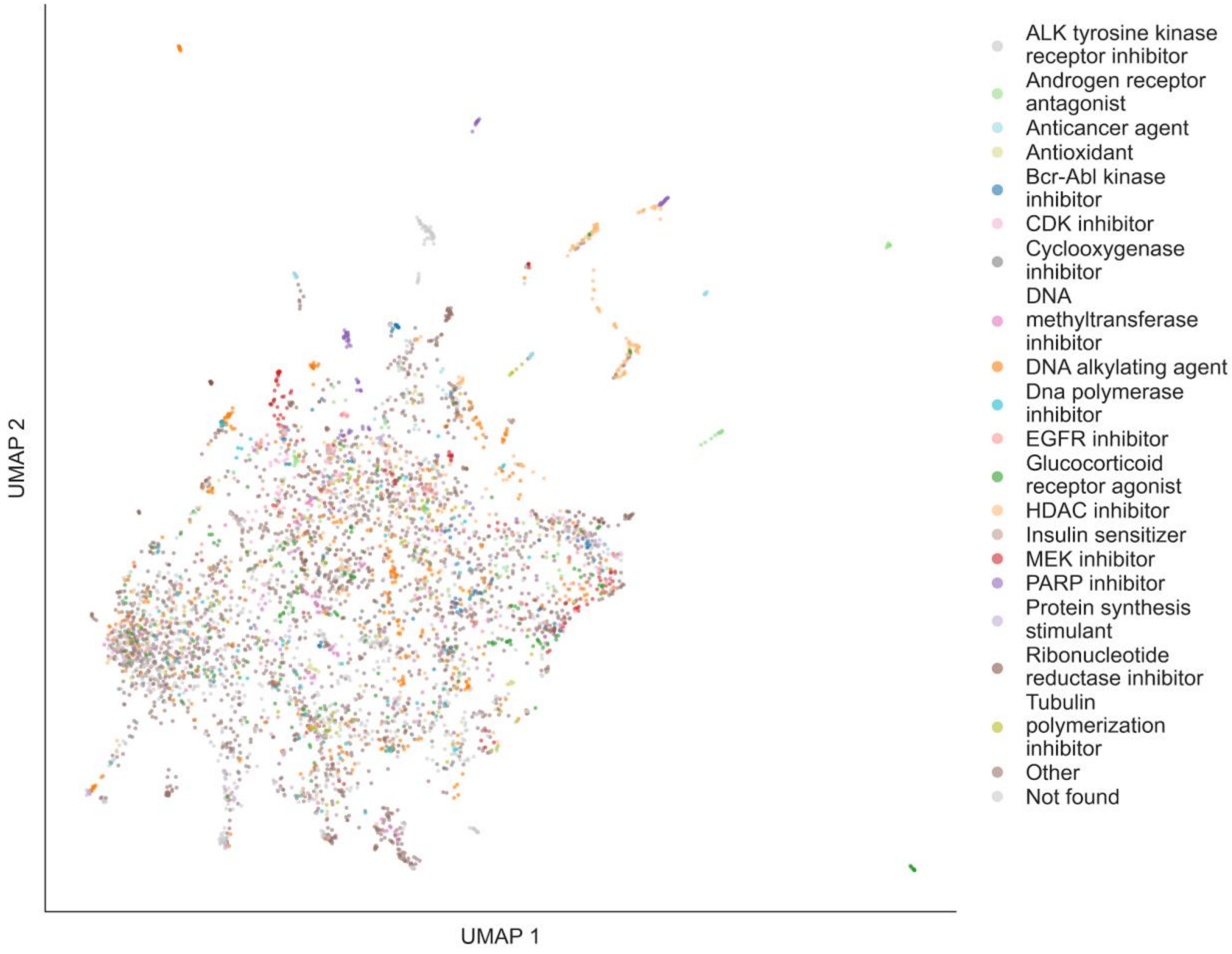


**Fig. S3-F1. RummaGEO Chemical Perturbations (MoA)**

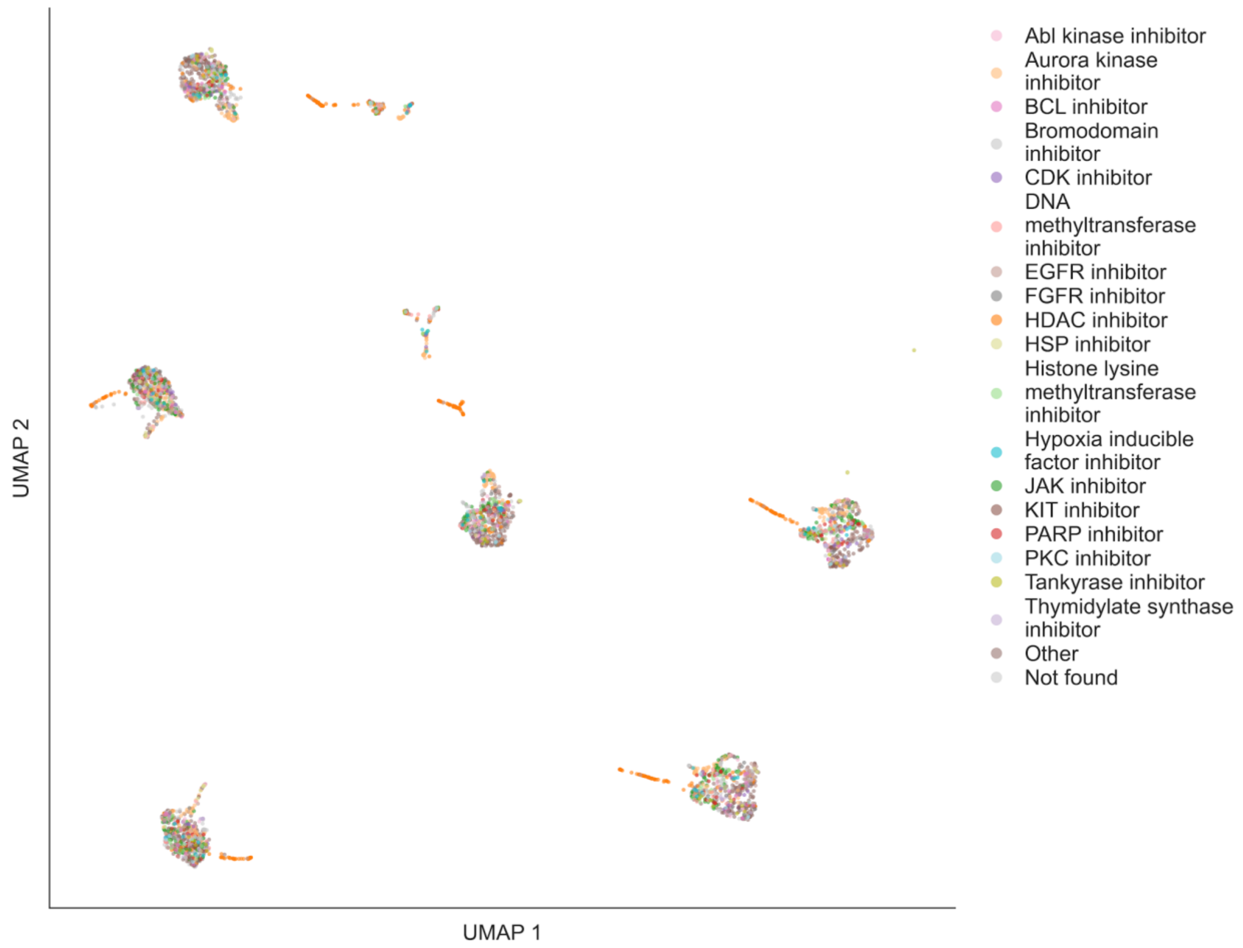


**Fig. S3-G1. SciPlex (MoA)**

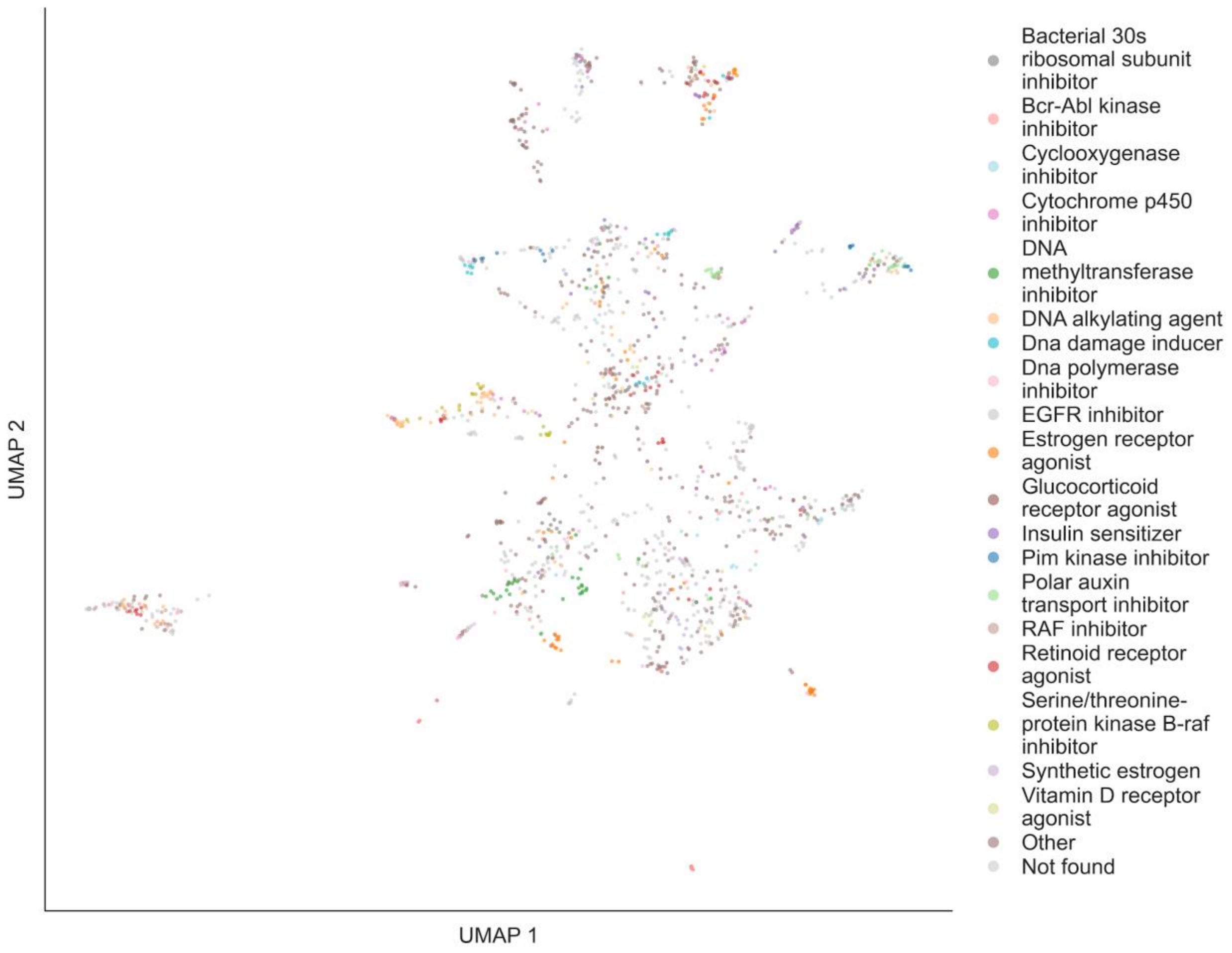


**Fig. S3-H1. CREEDS Chemical Perturbations (MoA)**

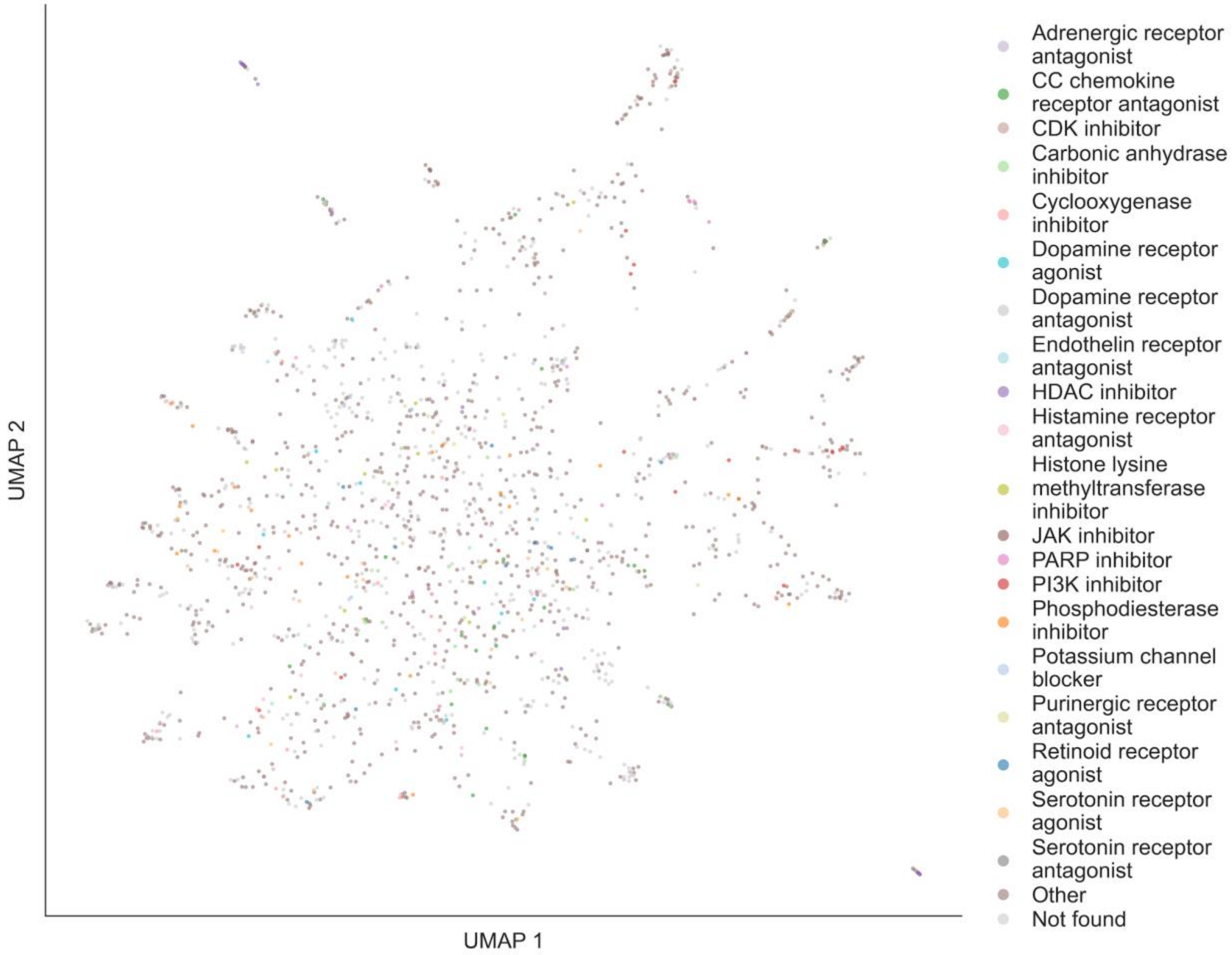


**Fig. S3-I1. DeepCover MoA (MoA)**

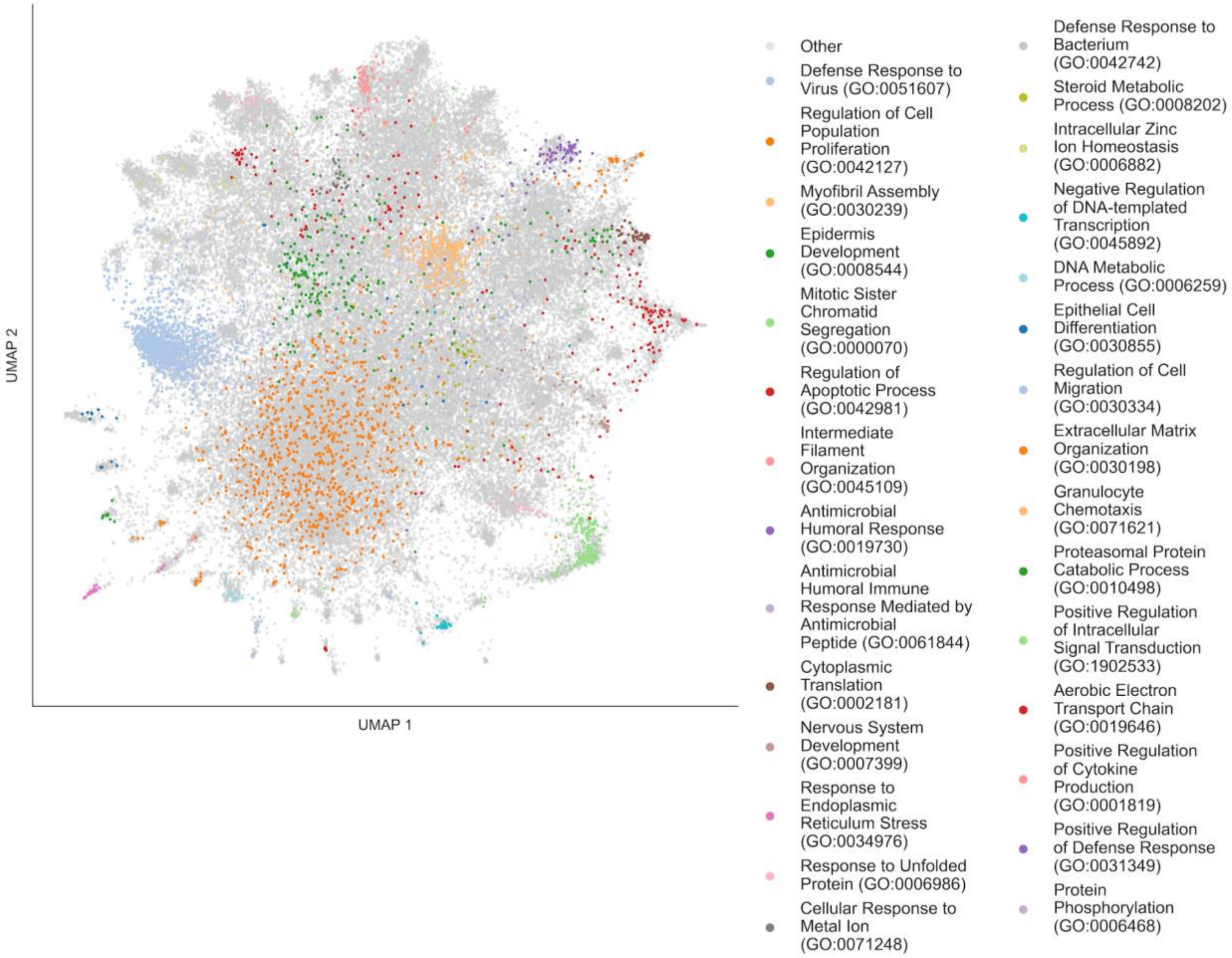


**Fig. S3-J1. LINCS L1000 Gene Perturbations (Enriched GO Term)**

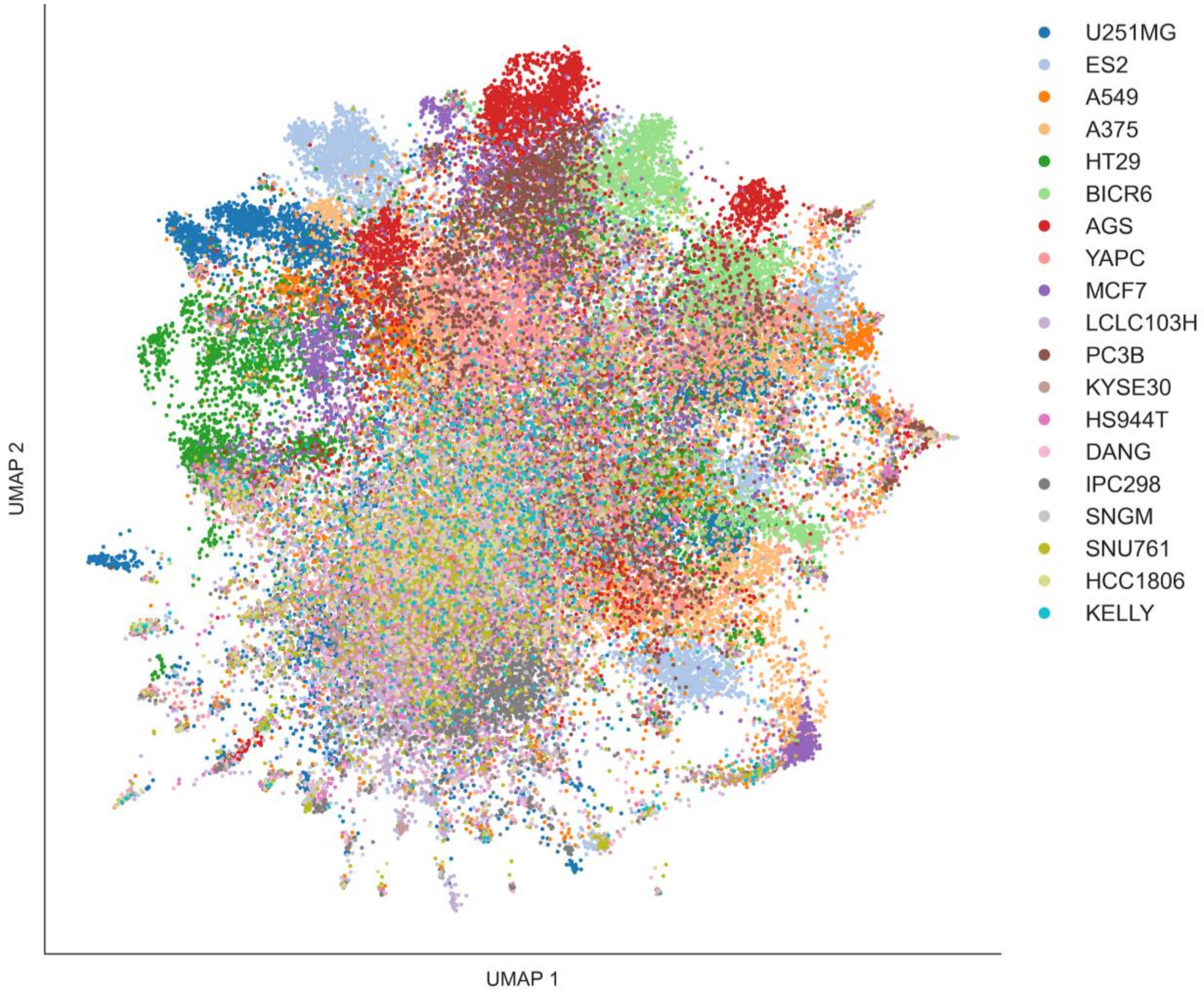


**Fig. S3-J2. LINCS L1000 Gene Perturbations (Cell Line)**

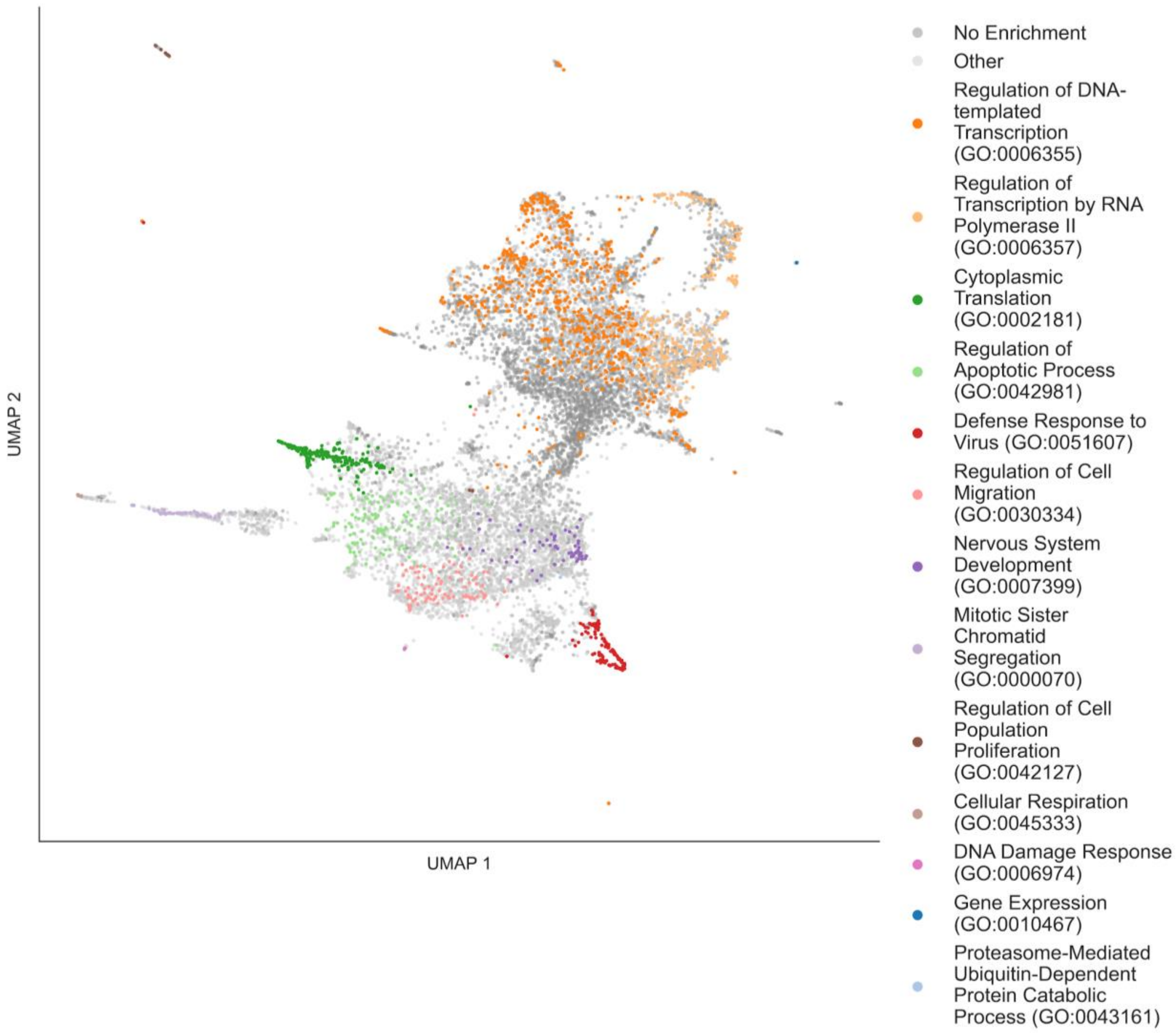


**Fig. S3-K1. Perturb Atlas Human (Enriched GO Term)**

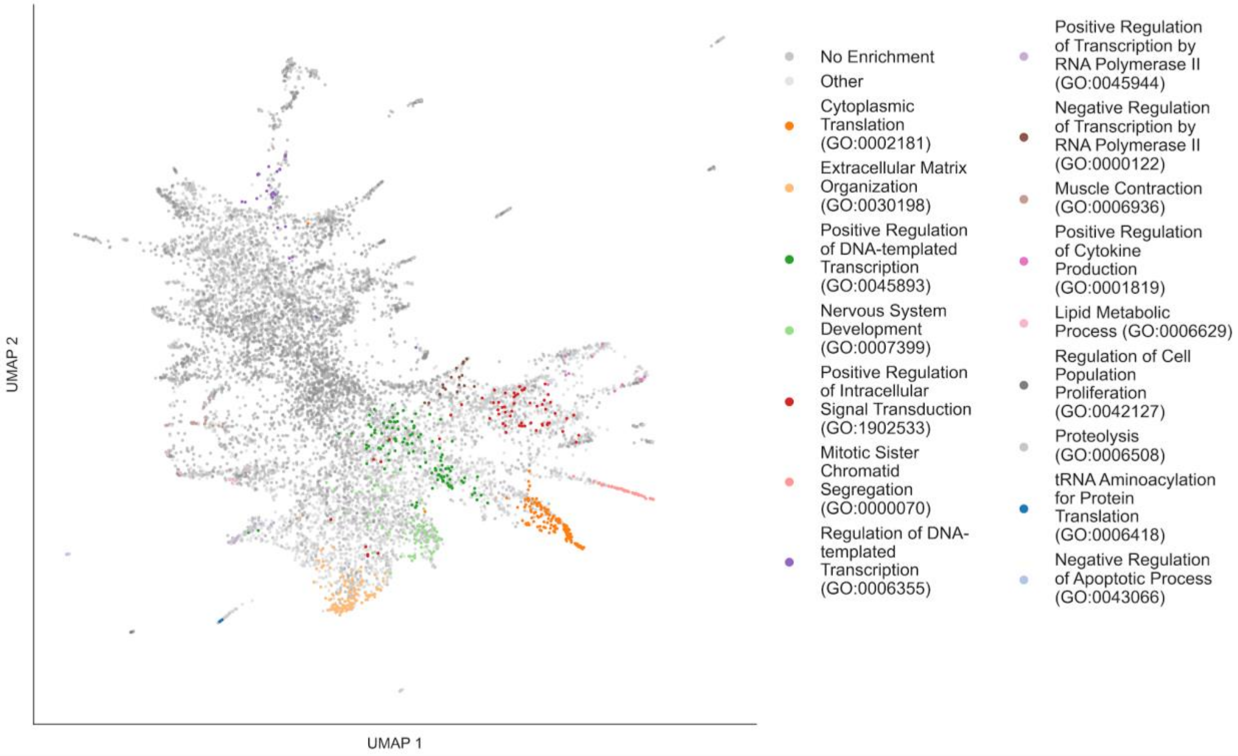


**Fig. S3-L1. Perturb Atlas Mouse (Enriched GO Term)**

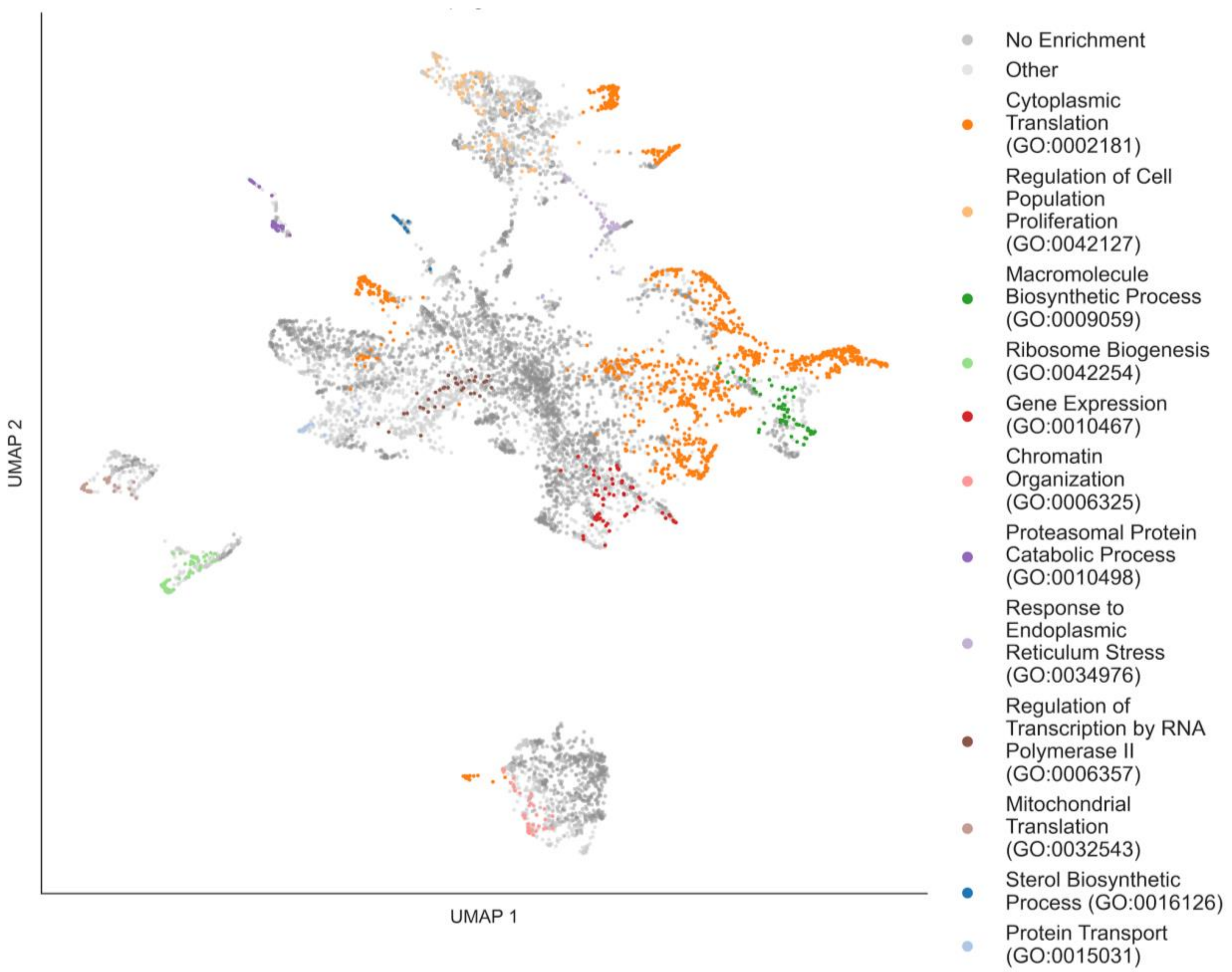


**Fig. S3-M1. Replogle et al. (Enriched GO Term)**

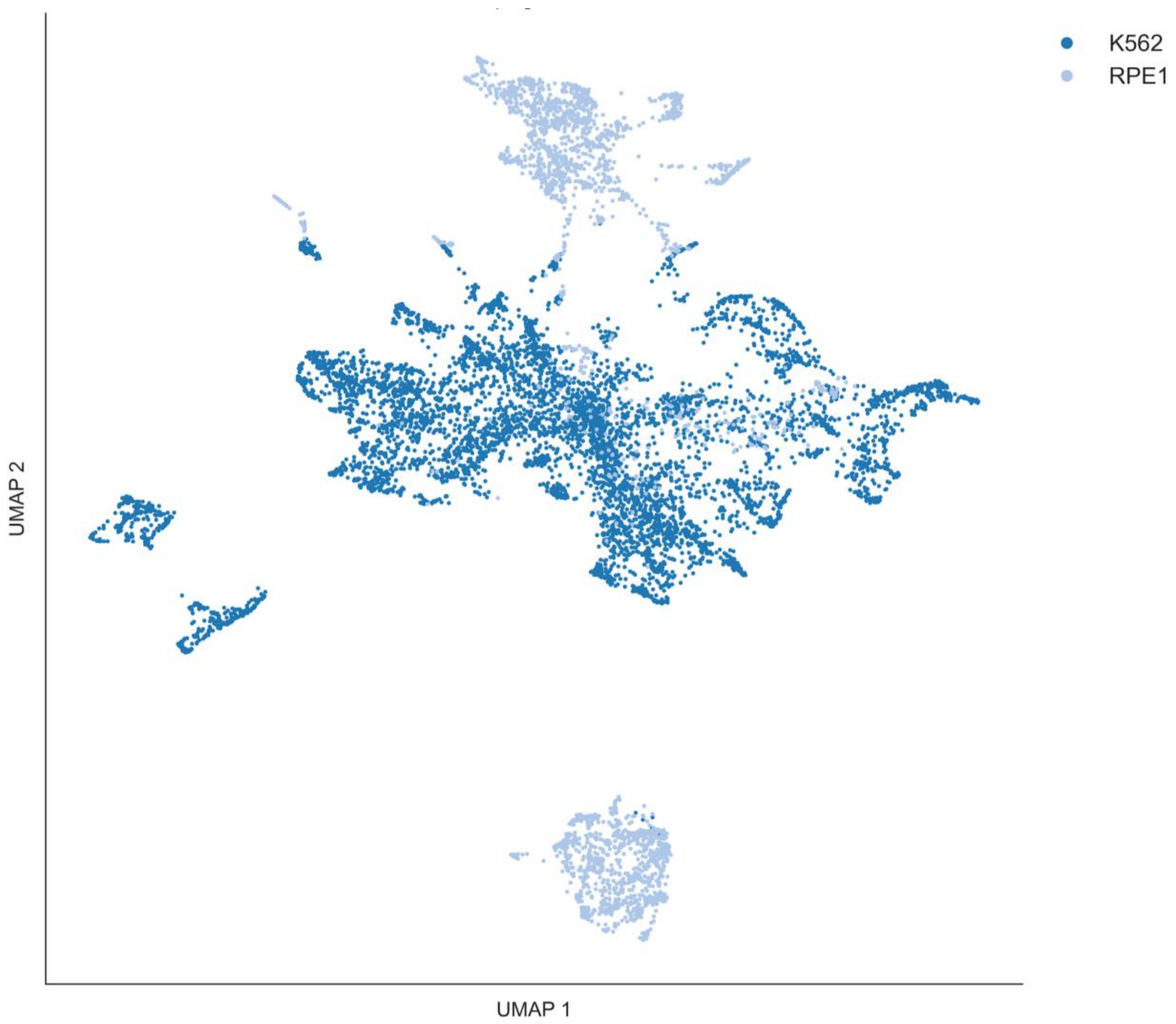


**Fig. S3-M2. Replogle et al. (Cell Line)**

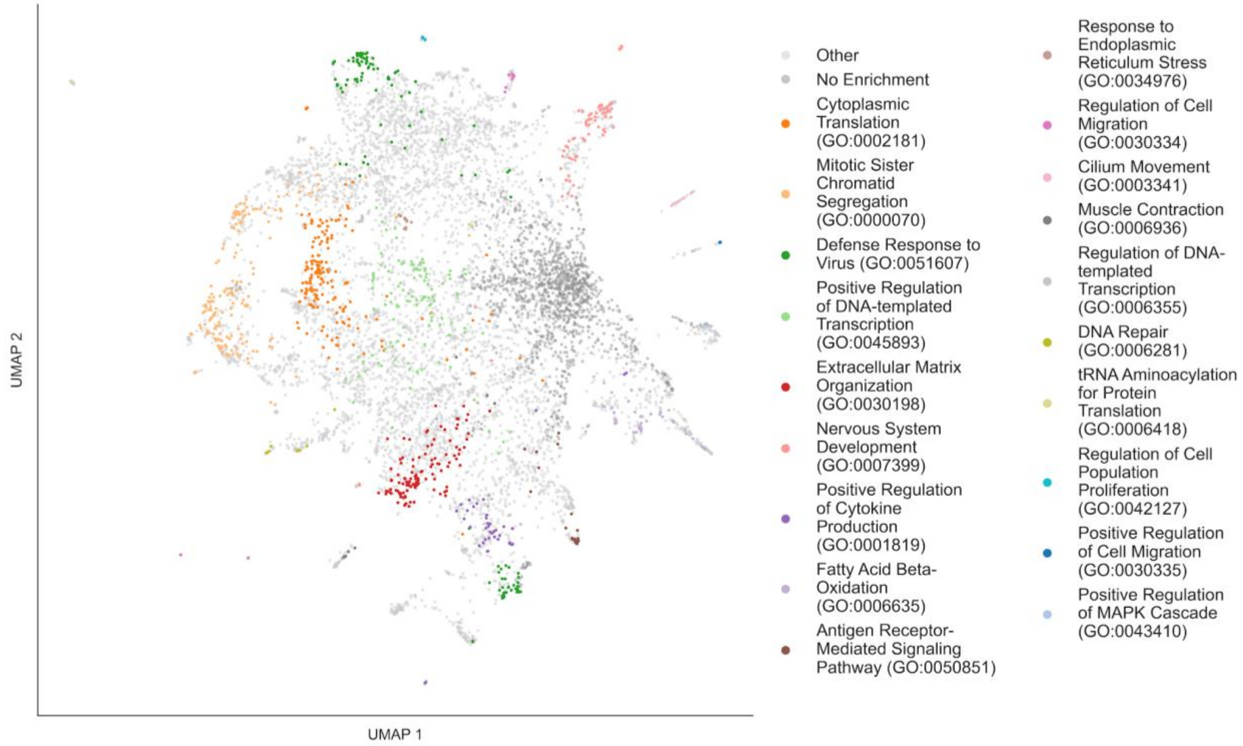


**Fig. S3-N1. RummaGEO Gene Perturbations (Enriched GO Term)**

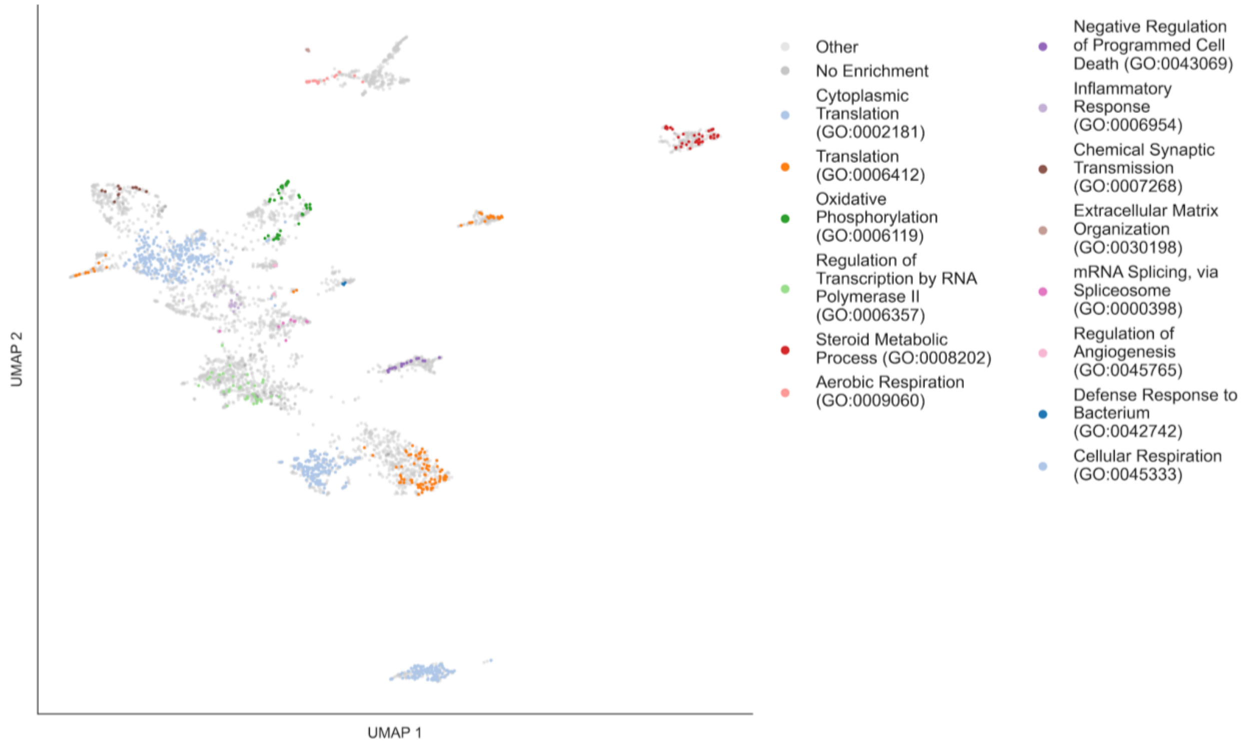


**Fig. S3-O1. CREEDS Gene Perturbations (Enriched GO Term)**

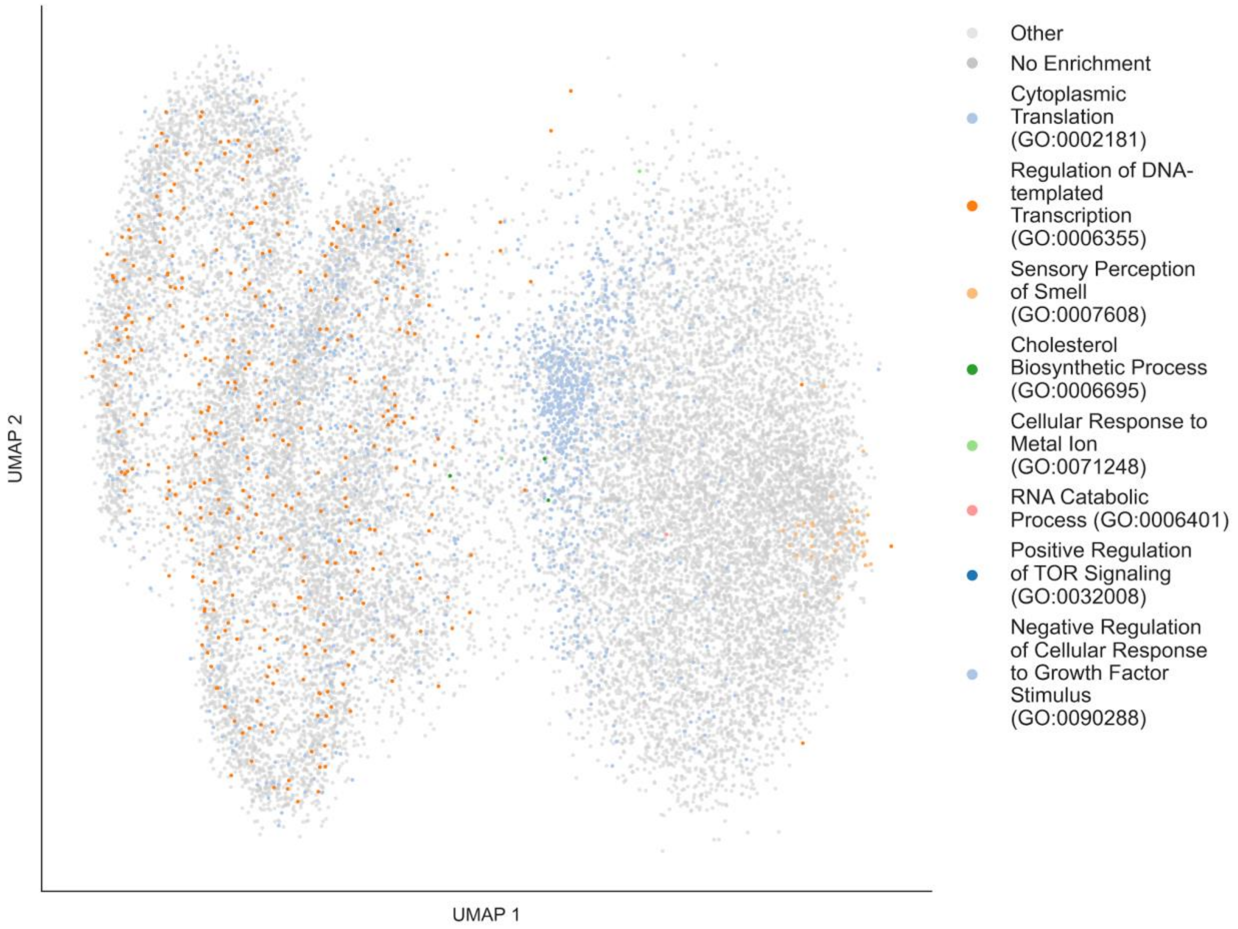


**Fig. S3-P1. CM4AI KOLF2.1J Genome-Wide Perturb-Seq (Enriched GO Term)**

**Fig. S3.** Drug perturbation (A-I) and gene perturbation (J-P) Connectivity Map UMAPs. Each gene set is colored by MoA (A1-I1) (drug perturbations), cell line (A2-I2, J2-M2) (drug and gene perturbations) if there is more than one cell line utilized in the resource, for the drug perturbations Connectivity Maps; and the most enriched GO terms (J1-P1) (gene perturbations Connectivity Maps).

A

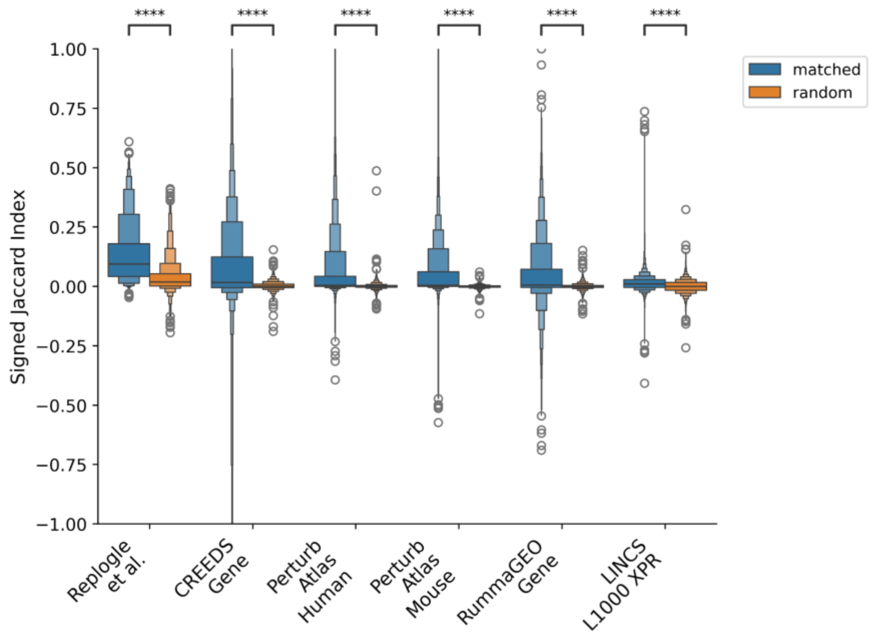

****
1.00
0.75
0.50
0.25
0.00
−0.25
−0.50
−0.75
−1.00
Signed Jaccard Index
matched
random
Replogle et al.
CREEDS Gene
Perturb Atlas Human
Perturb Atlas Mouse
RummaGEO Gene
LINCS L1000 XPR


B

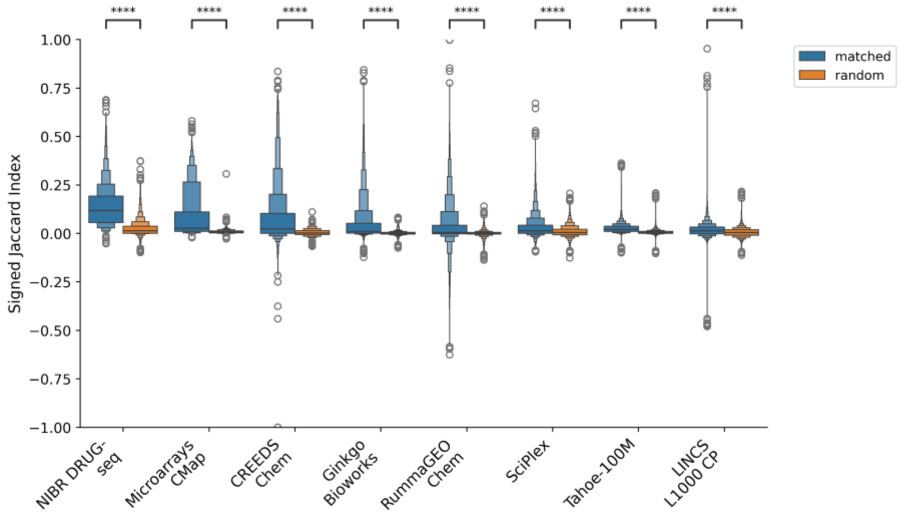

****
1.00
0.75
0.50
0.25
0.00
−0.25
−0.50
−0.75
−1.00
Signed Jaccard Index
matched
random
NIBR DRUG-seq
Microarrays CMap
CREEDS Chem
Ginkgo Bioworks
RummaGEO Chem
SciPlex
Tahoe-100M
LINCS L1000 CP

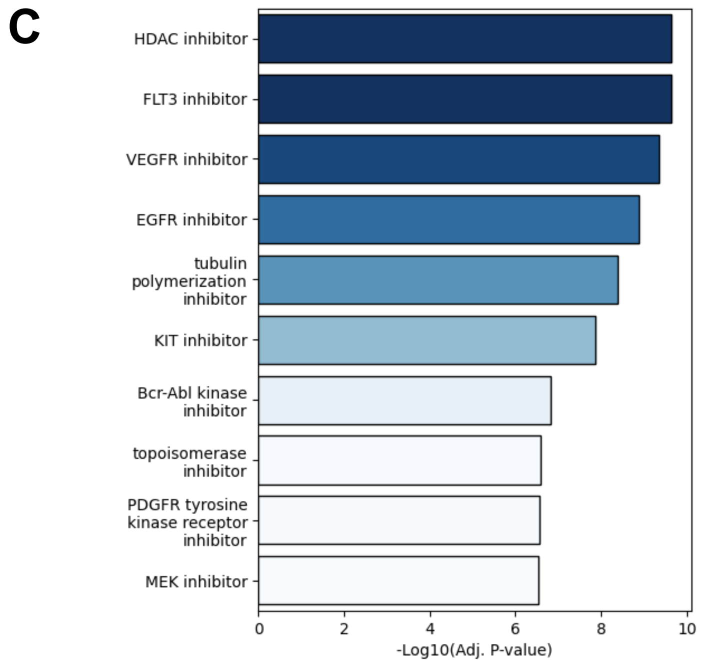


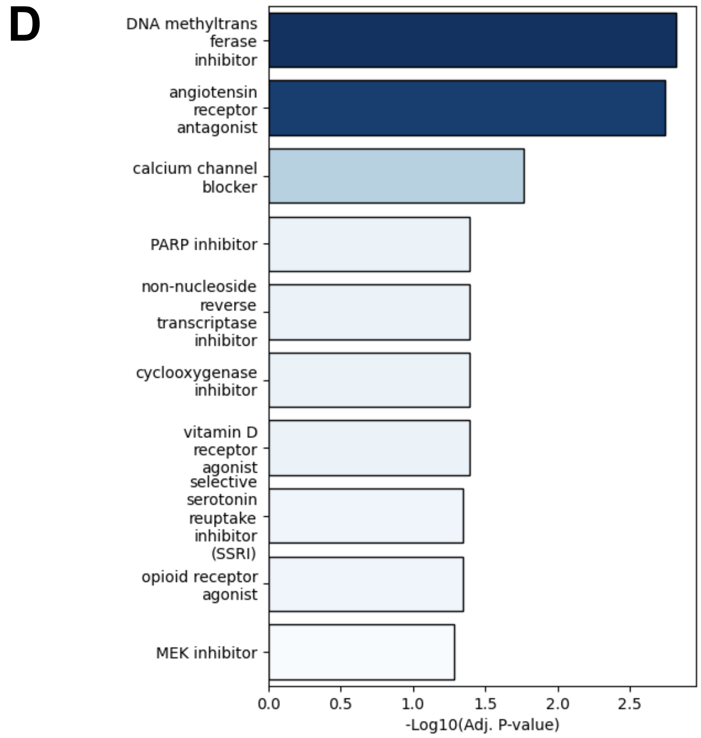


**Fig. S4.** Outlier perturbations by signed Jaccard index (A) for the gene perturbation Connectivity Maps, and (B) for the drug perturbation Connectivity Maps. (C, D) DrugEnrichr results for Drug Repurposing Hub MoA enrichment of outlier small molecule perturbagens for (C) high and (D) low signed Jaccard indices.

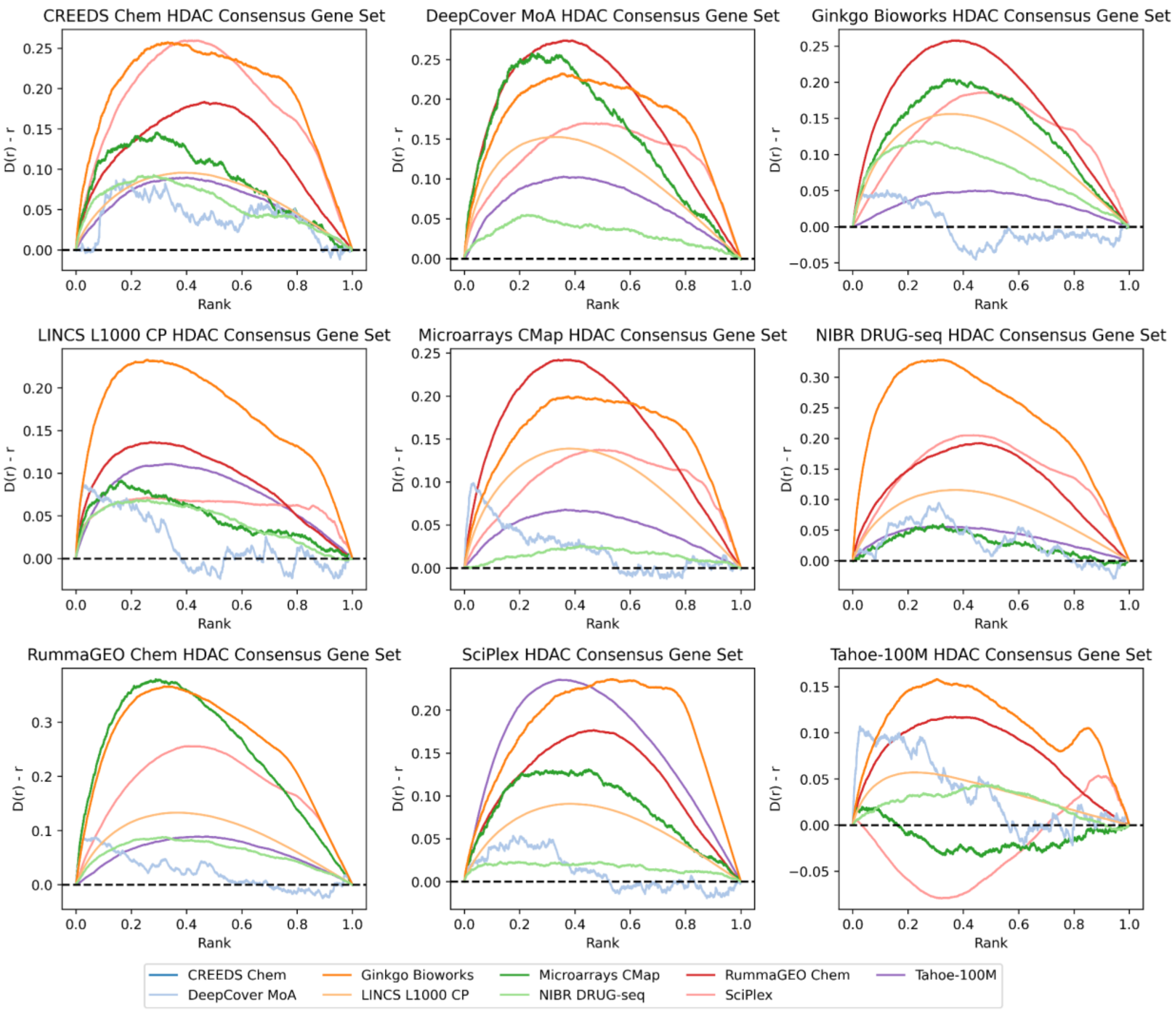


**Fig. S5.** Average enrichment analysis displayed as bridge plots of HDAC inhibitor signatures from each drug perturbation Connectivity Map resource against a consensus HDAC inhibitor gene set constructed from a left-out resource.

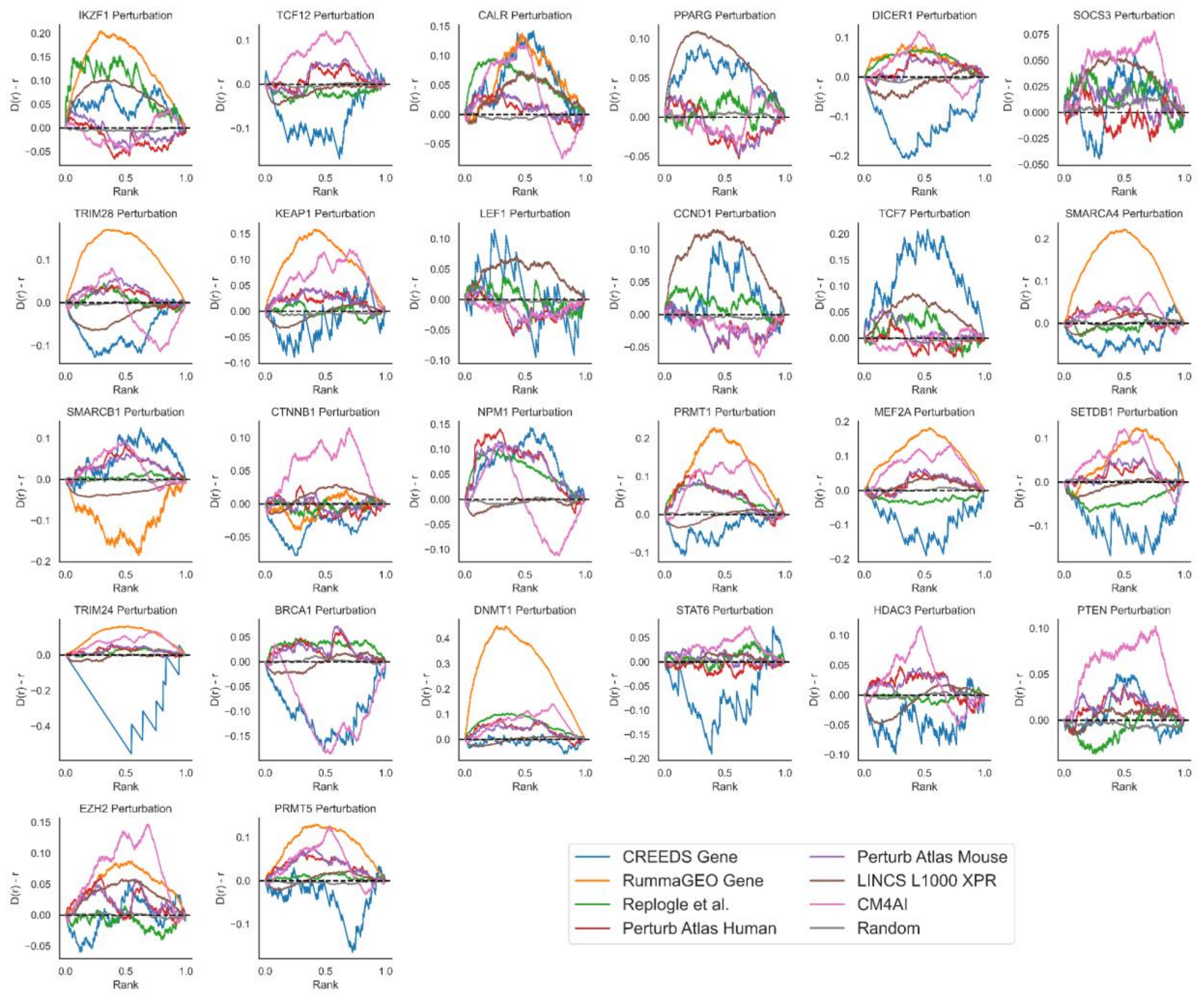


**Fig. S6.** Average enrichment analysis displayed as bridge plots of gene perturbation followed by expression signatures enriched against the 1000 most highly correlated genes with the perturbed gene based on mRNA gene-gene co-expression correlations computed from the ARCHS4 resource. Genes displayed have perturbation signatures in all gene perturbation Connectivity Map resources and in ARCHS4.

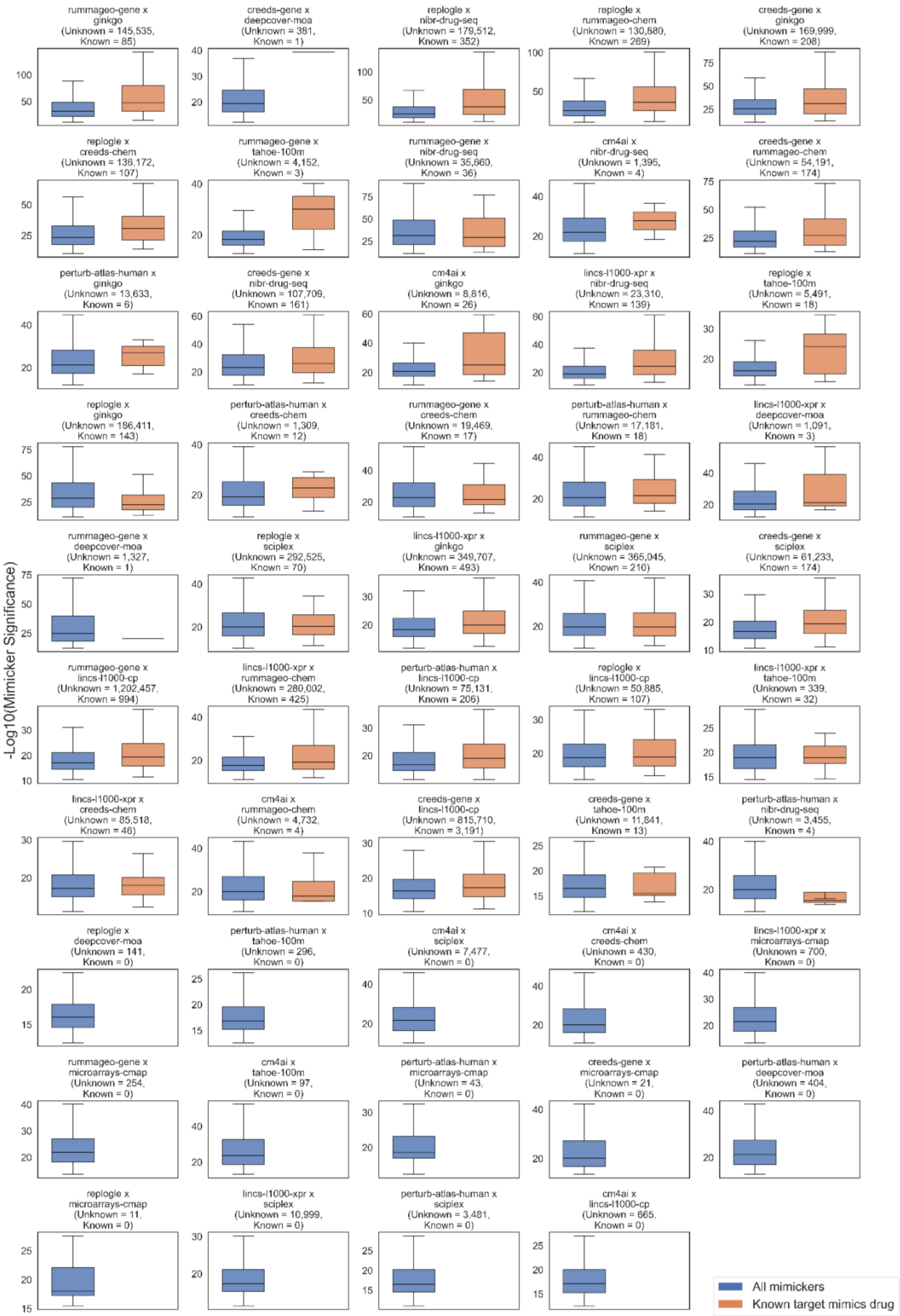

-Log10(Mimicker Significance)
rummageo-gene x ginkgo (Unknown = 145,535, Known = 85)
creeds-gene x deepcover-moa (Unknown = 381, Known = 1)
replogle x nibr-drug-seq (Unknown = 179,512, Known = 352)
replogle x rummageo-chem (Unknown = 130,880, Known = 269)
creeds-gene x ginkgo (Unknown = 169,999, Known = 208)
replogle x creeds-chem (Unknown = 136,172, Known = 107)
rummageo-gene x tahoe-100m (Unknown = 4,152, Known = 3)
rummageo-gene x nibr-drug-seq (Unknown = 35,660, Known = 36)
cm4ai x nibr-drug-seq (Unknown = 1,395, Known = 4)
creeds-gene x rummageo-chem (Unknown = 54,191, Known = 174)
perturb-atlas-human x ginkgo (Unknown = 13,633, Known = 6)
creeds-gene x nibr-drug-seq (Unknown = 107,709, Known = 161)
cm4ai x ginkgo (Unknown = 8,816, Known = 26)
lincs-l1000-xpr x nibr-drug-seq (Unknown = 23,310, Known = 139)
replogle x tahoe-100m (Unknown = 5,491, Known = 18)
replogle x ginkgo (Unknown = 186,411, Known = 143)
perturb-atlas-human x creeds-chem (Unknown = 1,309, Known = 12)
rummageo-gene x creeds-chem (Unknown = 19,469, Known = 17)
perturb-atlas-human x rummageo-chem (Unknown = 17,181, Known = 18)
lincs-l1000-xpr x deepcover-moa (Unknown = 1,091, Known = 3)
rummageo-gene x deepcover-moa (Unknown = 1,327, Known = 1)
replogle x sciplex (Unknown = 292,525, Known = 70)
lincs-l1000-xpr x ginkgo (Unknown = 349,707, Known = 493)
rummageo-gene x sciplex (Unknown = 365,045, Known = 210)
creeds-gene x sciplex (Unknown = 61,233, Known = 174)
rummageo-gene x lincs-l1000-cp (Unknown = 1,202,457, Known = 994)
lincs-l1000-xpr x rummageo-chem (Unknown = 280,002, Known = 425)
perturb-atlas-human x lincs-l1000-cp (Unknown = 75,131, Known = 206)
replogle x lincs-l1000-cp (Unknown = 50,885, Known = 107)
lincs-l1000-xpr x tahoe-100m (Unknown = 339, Known = 32)
lincs-l1000-xpr x creeds-chem (Unknown = 85,518, Known = 46)
cm4ai x rummageo-chem (Unknown = 4,732, Known = 4)
creeds-gene x lincs-l1000-cp (Unknown = 815,710, Known = 3,191)
creeds-gene x tahoe-100m (Unknown = 11,841, Known = 13)
perturb-atlas-human x nibr-drug-seq (Unknown = 3,455, Known = 4)
replogle x deepcover-moa (Unknown = 141, Known = 0)
perturb-atlas-human x tahoe-100m (Unknown = 296, Known = 0)
cm4ai x sciplex (Unknown = 7,477, Known = 0)
cm4ai x creeds-chem (Unknown = 430, Known = 0)
lincs-l1000-xpr x microarrays-cmap (Unknown = 700, Known = 0)
rummageo-gene x microarrays-cmap (Unknown = 254, Known = 0)
cm4ai x tahoe-100m (Unknown = 97, Known = 0)
perturb-atlas-human x microarrays-cmap (Unknown = 43, Known = 0)
creeds-gene x microarrays-cmap (Unknown = 21, Known = 0)
perturb-atlas-human x deepcover-moa (Unknown = 404, Known = 0)
replogle x microarrays-cmap (Unknown = 11, Known = 0)
lincs-l1000-xpr x sciplex (Unknown = 10,999, Known = 0)
perturb-atlas-human x sciplex (Unknown = 3,481, Known = 0)
cm4ai x lincs-l1000-cp (Unknown = 665, Known = 0)
All mimickers
Known target mimics drug

**Fig. S7.** Distribution of Fisher's exact test p-value for mimickers across drug and gene library crosses. A mimicker is a pair of signatures whose up-regulated and down-regulated gene sets overlap significantly, as determined by Fisher's exact test. For each drug-gene library cross, the significance of all drug-gene mimicking pairs is compared with the subset of pairs in which the gene is a known target of the drug. Some crosses yielded no significant mimickers between drugs and their known targets at $p < 1x10^{-5}$; for those crosses only the distribution of all pairs is shown. The number of mimicking signature pairs below this significant threshold is displayed in each subplot. Crosses were restricted to libraries from different Connectivity Map resources (e.g. LINCS drug perturbations were not crossed with LINCS gene perturbations).

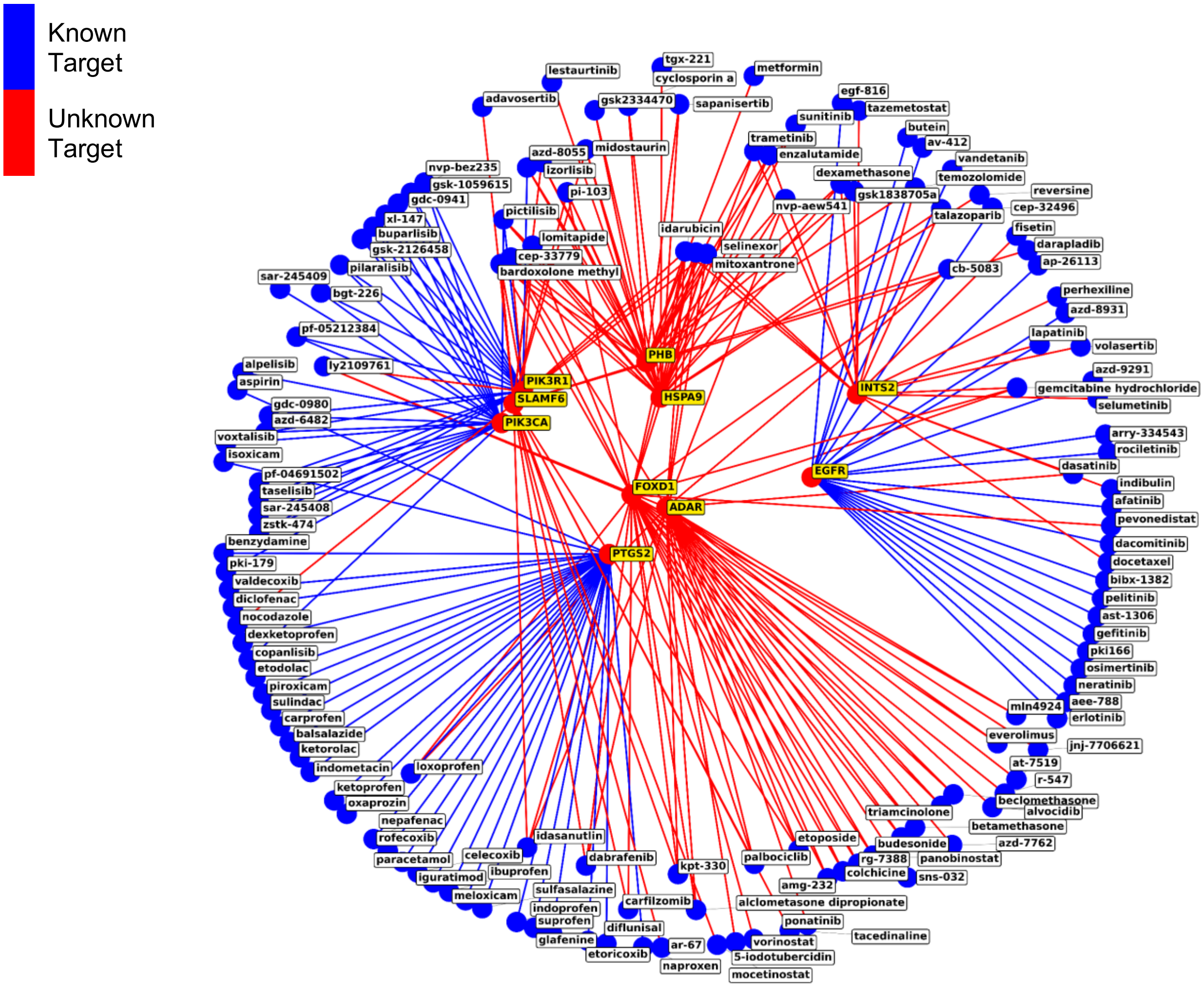


**Fig. S8.** Gene perturbations with the largest number of distinct small molecule mimickers at Fisher's exact test p-value $< 1x10^{-5}$. Gene perturbation nodes (red) are highlighted in yellow. Blue edges indicate the gene is listed as a target of the drug in OpenTargets and red edges indicate the gene is not listed as a target.

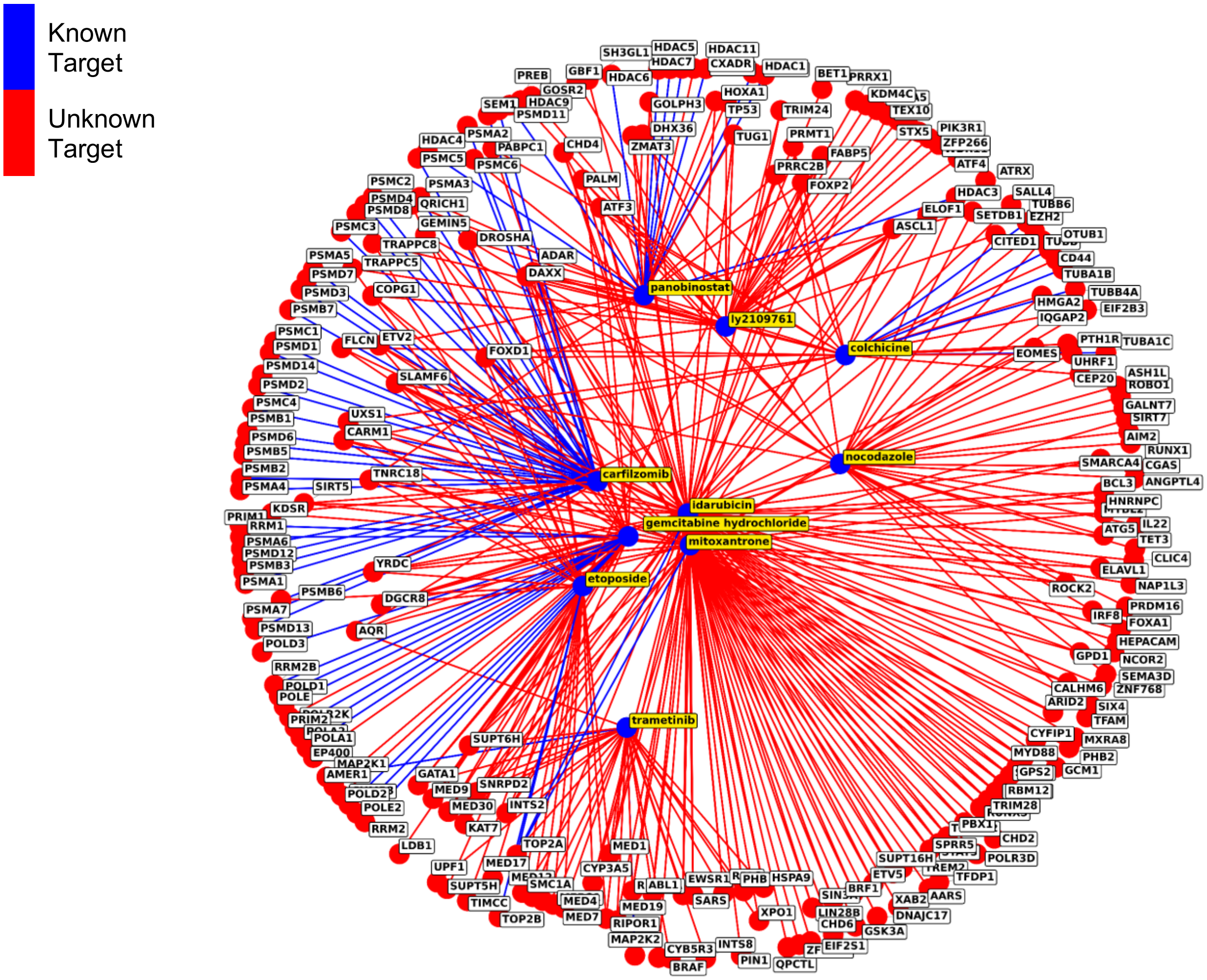


**Fig. S9.** Drug perturbations with the largest number of distinct gene mimickers at Fisher's exact test p-value < $1x10^{-5}$. Drug perturbation nodes (blue) are highlighted in yellow. Blue edges indicate the gene is listed as a target of the drug in OpenTargets and red edges indicate the gene is not listed as a target.

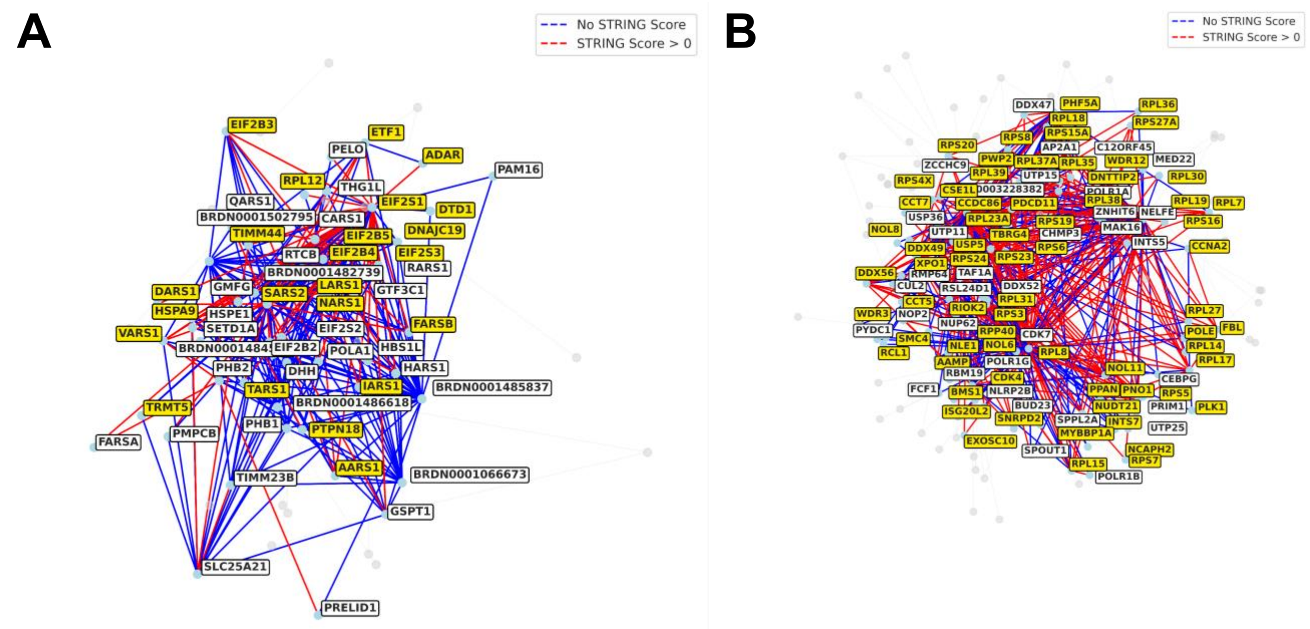


**Fig. S10.** Triangle subnetworks in Louvain communities 1 and 2 from Fig. 7. Genes in each community were submitted to Enrichr and the subset of genes in the community's most significantly enriched ChEA TF library are highlighted. Edge color indicates presence or absence of PPI evidence in the STRING database. (A) Community 1 with MYC TF targets highlighted for term *ChEA MYC 22102868 ChIP-Seq CA46 Human Blood Burkitts Lymphoma* (2314/23/48, ChEA term/overlap/community 1) and (B) community 2 with MYC TF targets highlighted for term *ChEA MYC 19030024 ChIP-ChIP MESCs Mouse* (2842/68/102, ChEA term/overlap/community 2). Full enrichment results for each of the eight identified communities are provided (Table S7).

### Supporting Tables

| Library | Perturbation | Signed Jaccard Index | Outlier Status |
|---|---|---|---|
| CREEDS Gene | E2A | 1.000 | outlier_hi |
| CREEDS Gene | TP53INP2 | 1.000 | outlier_hi |
| CREEDS Gene | PTHRP | 1.000 | outlier_hi |
| CREEDS Gene | FBXL10 | 1.000 | outlier_hi |
| CREEDS Gene | PFKL | 1.000 | outlier_hi |
| LINCS L1000 XPR | HSPA5 | 0.736 | outlier_hi |
| LINCS L1000 XPR | CTNNB1 | 0.699 | outlier_hi |
| LINCS L1000 XPR | SNRPA1 | 0.681 | outlier_hi |
| LINCS L1000 XPR | PSMA5 | 0.663 | outlier_hi |
| LINCS L1000 XPR | KPNB1 | 0.652 | outlier_hi |
| Perturb Atlas Human | UPF1 | 1.000 | outlier_hi |
| Perturb Atlas Human | IGF2BP1 | 1.000 | outlier_hi |
| Perturb Atlas Human | ERG | 1.000 | outlier_hi |
| Perturb Atlas Human | METTL14 | 1.000 | outlier_hi |
| Perturb Atlas Human | METTL3 | 1.000 | outlier_hi |
| Perturb Atlas Mouse | TRIM24 | 1.000 | outlier_hi |
| Perturb Atlas Mouse | NRF2 | 1.000 | outlier_hi |
| Perturb Atlas Mouse | KCNQ1OT1 | 1.000 | outlier_hi |
| Perturb Atlas Mouse | MYC | 1.000 | outlier_hi |
| Perturb Atlas Mouse | BMAL1 | 1.000 | outlier_hi |
| Replogle et al. | ACTR8 | 0.609 | outlier_hi |
| Replogle et al. | FCF1 | 0.566 | outlier_hi |
| Replogle et al. | KRR1 | 0.563 | outlier_hi |
| Replogle et al. | TSR2 | 0.559 | outlier_hi |
| Replogle et al. | WDR36 | 0.559 | outlier_hi |
| RummaGEO Gene | MTOR | 1.000 | outlier_hi |
| RummaGEO Gene | DNMT3A | 0.932 | outlier_hi |
| RummaGEO Gene | SRPK3 | 0.807 | outlier_hi |
| RummaGEO Gene | MERTK | 0.787 | outlier_hi |
| RummaGEO Gene | MYD88 | 0.753 | outlier_hi |

**Table S1.** The 5 gene perturbations with the most consistent transcriptomics effect for each gene perturbation Connectivity Map resource. A gene perturbation's signed Jaccard index measurement for set consistency is defined as an outlier at the high end of the distribution if it is > Q3 + 1.5 * IQR (interquartile range).

| Library | Perturbation | Signed Jaccard Index | Outlier Status |
|---|---|---|---|
| CREEDS Chem | Tretinoin | 0.835 | outlier_hi |
| CREEDS Chem | Perfluorooctanoic acid | 0.783 | outlier_hi |
| CREEDS Chem | Imatinib | 0.746 | outlier_hi |
| CREEDS Chem | Vancomycin | 0.682 | outlier_hi |
| CREEDS Chem | Decitabine | 0.652 | outlier_hi |
| Ginkgo Bioworks | Brefeldin-A | 0.844 | outlier_hi |
| Ginkgo Bioworks | Panobinostat | 0.699 | outlier_hi |
| Ginkgo Bioworks | Trichostatin A | 0.697 | outlier_hi |
| Ginkgo Bioworks | Dexamethasone | 0.693 | outlier_hi |
| Ginkgo Bioworks | Thapsigargin | 0.660 | outlier_hi |
| LINCS L1000 CP | PCI-24781 | 0.953 | outlier_hi |
| LINCS L1000 CP | I-BET-151 | 0.813 | outlier_hi |
| LINCS L1000 CP | vorinostat | 0.799 | outlier_hi |
| LINCS L1000 CP | GSK-461364 | 0.774 | outlier_hi |
| LINCS L1000 CP | sirolimus | 0.755 | outlier_hi |
| Microarrays CMap | trichostatin A | 0.581 | outlier_hi |
| Microarrays CMap | vorinostat | 0.517 | outlier_hi |
| Microarrays CMap | lanatoside C | 0.471 | outlier_hi |
| Microarrays CMap | cephaeline | 0.451 | outlier_hi |
| Microarrays CMap | MG-262 | 0.427 | outlier_hi |
| NIBR DRUG-seq | Cinobufagin | 0.689 | outlier_hi |
| NIBR DRUG-seq | Nannocystin A | 0.685 | outlier_hi |
| NIBR DRUG-seq | Arenobufagin | 0.674 | outlier_hi |
| NIBR DRUG-seq | (1R,2R,3E,7S,11Z,13S,15S)-2,15-dihydroxy-7-methyl-6-oxabicyclo[11.3.0]hexadeca-3,11-dien-5-one | 0.673 | outlier_hi |
| NIBR DRUG-seq | CID 56946209 | 0.655 | outlier_hi |
| RummaGEO Chem | morphine | 0.999 | outlier_hi |
| RummaGEO Chem | hiltonol | 0.997 | outlier_hi |
| RummaGEO Chem | ethanol | 0.853 | outlier_hi |
| RummaGEO Chem | oxymetholone | 0.837 | outlier_hi |
| RummaGEO Chem | talazoparib | 0.776 | outlier_hi |
| SciPlex | Panobinostat | 0.671 | outlier_hi |
| SciPlex | Quisinostat | 0.644 | outlier_hi |
| SciPlex | Dacinostat | 0.515 | outlier_hi |

| | | | |
|---|---|---|---|
| SciPlex | JQ1 | 0.436 | outlier_hi |
| SciPlex | Abexinostat | 0.403 | outlier_hi |
| Tahoe-100M | Panobinostat | 0.362 | outlier_hi |
| Tahoe-100M | Dexamethasone | 0.354 | outlier_hi |
| Tahoe-100M | Clobetasol propionate | 0.326 | outlier_hi |
| Tahoe-100M | Budesonide | 0.300 | outlier_hi |
| Tahoe-100M | Betamethasone dipropionate | 0.295 | outlier_hi |

**Table S2.** The 5 gene perturbations with the most consistent transcriptomics effect for each drug perturbation Connectivity Map resource. A drug perturbation's signed Jaccard index measurement for set consistency is defined as an outlier at the high end of the distribution if it is > Q3 + 1.5 * IQR (interquartile range).

| ID | N Genes | Likely Function Based on Enrichment | Enrichr Links |
|---|---|---|---|
| 0 | 34 | mRNA splicing | 28052956f81b3d2291bd4534c082442c |
| 1 | 75 | MYC targets, tRNA processing, translation | 6d05e6699f2a8a5bc6028dba10425953 |
| 2 | 158 | MYC targets, tRNA processing, translation | 969eba8126b26d5c127b2983e798a7d2 |
| 3 | 84 | Vesicle transport | 1827d26b23937e7fbff0cd23b1602857 |
| 4 | 3 | N/A | 2d64dda73c24bcca08533e550498fc08 |
| 5 | 18 | Hematopoiesis and skin diseases | 76838b1f1e63a393191badc8d06bb4a7 |
| 6 | 48 | Cell cycle | 7c73b9247ffd09068cc6e39e6d223310 |
| 7 | 63 | Members of the proteasome | 33abac4314eb154198dc80dfc33df2a8 |
| 8 | 32 | Mediator complex | 7986ab4e060920a88980f7459b942a60 |

**Table S7.** Enrichr enrichment results for each community of gene perturbation mimickers identified by crossing all the LINCS L1000 single gene perturbation signatures with all the Replogle et al. Perturb-Seq single gene perturbation signatures. Densely connected communities in the network of gene-gene mimicking pairs were identified with the Louvain community detection algorithm as implemented in the NetworkX Python package. The general function for each community is summarized by a few terms manually by scanning the Enrichr results.

*The remaining supporting tables are provided as Excel files.*

**Table S3.** The top 1000 drug and gene perturbation mimicking pairs, identified by those pairs that had the highest number of mimicking signatures significant at Fisher's exact test p-value < $1x10^{-5}$ for overlap among up-regulated genes and overlap among down-regulated genes.

**Table S4.** Mimicking drug-gene signature pairs among ARID1A knockouts and knockdowns and dasatinib perturbations across Connectivity Map libraries in Perturb-Seqr.

**Table S5.** Top 1000 drug and gene signature pairs by the product of Fisher's exact test p-value for overlap among up-regulated genes and overlap among down-regulated genes.

**Table S6.** Drug and INST2 perturbation signature pairs with Fisher's exact test p-value < $1x10^{-5}$ for overlap among up-regulated genes and overlap among down-regulated genes**.**

**Table S8.** Enrichr enrichment analysis results for gene sets extracted from communities of highly connected gene perturbation mimickers, identified using the Louvain algorithm on a network of mimicking pairs identified from crossing all signatures in the LINCS L1000 gene perturbation Connectivity Map with the Replogle et al. Connectivity Map. Significantly enriched terms from the following libraries are shown: KEGG pathways, GO Biological Process terms, ChEA TF gene sets, GWAS Catalog associations, and MGI Mammalian phenotypes.

**Table S9.** Significant drug perturbation and gene perturbation mimickers of endurance and resistance exercise signatures in muscle 3.5-4 hours after an acute bout of exercise. Significant mimickers were identified by querying the Perturb-Seqr database with the paired enrichment query, which compares an input signature's up- and down-regulated gene sets to the Perturb-Seqr collection.
